# Non-canonical open reading frames encode functional proteins essential for cancer cell survival

**DOI:** 10.1101/2020.03.10.981001

**Authors:** John R. Prensner, Oana M. Enache, Victor Luria, Karsten Krug, Karl R. Clauser, Joshua M. Dempster, Amir Karger, Li Wang, Karolina Stumbraite, Vickie M. Wang, Ginevra Botta, Nicholas J. Lyons, Amy Goodale, Zohra Kalani, Briana Fritchman, Adam Brown, Douglas Alan, Thomas Green, Xiaoping Yang, Jacob D. Jaffe, Jennifer A. Roth, Federica Piccioni, Marc W. Kirschner, Zhe Ji, David E. Root, Todd R. Golub

## Abstract

A key question in genome research is whether biologically active proteins are restricted to the ∼20,000 canonical, well-annotated genes, or rather extend to the many non-canonical open reading frames (ORFs) predicted by genomic analyses. To address this, we experimentally interrogated 553 ORFs nominated in ribosome profiling datasets. Of these 553 ORFs, 57 (10%) induced a viability defect when the endogenous ORF was knocked out using CRISPR/Cas9 in 8 human cancer cell lines, 257 (46%) showed evidence of protein translation when ectopically expressed in HEK293T cells, and 401 (73%) induced gene expression changes measured by transcriptional profiling following ectopic expression across 4 cell types. CRISPR tiling and start codon mutagenesis indicated that the biological effects of these non-canonical ORFs required their translation as opposed to RNA-mediated effects. We selected one of these ORFs, *G029442*--renamed *GREP1* (Glycine-Rich Extracellular Protein-1)**--**for further characterization. We found that *GREP1* encodes a secreted protein highly expressed in breast cancer, and its knock-out in 263 cancer cell lines showed preferential essentiality in breast cancer derived lines. Analysis of the secretome of GREP1-expressing cells showed increased abundance of the oncogenic cytokine GDF15, and GDF15 supplementation mitigated the growth inhibitory effect of *GREP1* knock-out. Taken together, these experiments suggest that the non-canonical ORFeome is surprisingly rich in biologically active proteins and potential cancer therapeutic targets deserving of further study.

---

Early analyses of the human genome sequence suggested the existence of 100,000 or more protein-coding genes, but further scrutiny revealed that the majority of those candidate genes were more likely producing non-coding RNAs, fragmented cDNA clones, or RNAs expressed at inconsequential levels^1–3^. The current Human Proteome Project NeXtProt database recognizes ∼17,600 proteins confirmed by mass spectrometry and ∼2,100 unconfirmed proteins^4^.

Nevertheless, a growing body of evidence utilizing high-throughput profiling of ribosome-associated RNAs suggests that additional, non-canonical translation exists in genes currently annotated as noncoding RNAs or pseudogenes, as well as 5’ and 3’ untranslated regions (UTRs) of protein-coding genes^5–8^. Yet, it is unclear whether such translation reflects proteins overlooked during the construction of reference genome databases^9–12^, leaky ribosome scanning, or confounded computational predictions^13, 14^, since stringent conservation-based analyses have added only a small number of new proteins to the human genome^15^. Indeed, systematic experimental evidence interrogating whether such predicted proteins are in fact stably translated and biologically functional is lacking.

To address this, we curated a list 553 high priority ORFs nominated in lncRNAs and regions upstream and downstream of known protein coding genes (uORFs and dORFs, respectively). These were selected based on published predictions of ORF translation, additional analyses to eliminate pseudogenes, and exclusion of ORFs representing variants of known protein coding regions^5, 6, 14, 16–19, 19–33^ (Supplementary Table 1, Methods). 203/553 (37%) were identified as translated by at least two independent studies. We annotated the 553 ORFs according to 12 metrics including evolutionary conservation, expression and structural features (Supplementary Tables 2-11, Methods). 518 of 553 selected ORFs (94%) scored highly for at least one metric, with 418 (76%) having multiple lines of evidence in support of relevance (Supplementary Fig. 1 and Supplementary Table 2).

We next asked whether systematic functional studies could test the predicted translation of these ORFs (Fig. 1a). The capacity for the ORFs to produce a stably translated protein was assessed by three independent methods. First, we queried publicly-available mass spectrometry databases (Methods) and observed 6,724 peptides (707 unique) supporting 174 of 553 ORFs (31%) (Supplementary Tables 12 & 13). Next, we designed an expression library of the 553 ORFs containing a V5 epitope tag and developed a scalable assay for individual protein evaluation by anti-V5 detection (Fig. 1b and Supplementary Fig. 2a-d). 257 ORFs (46%) yielded a V5-tagged protein detectable by in-cell visualization (Fig. 1c-e, Supplementary Fig. 2e-g and Supplementary Table 14). Lastly, we detected a protein for 10 of 30 ORFs tested by *in vitro* transcription and translation (Supplementary Fig. 2h). Taken together, experimental evidence of protein translation was obtained for 334/553 (60%) of the ORFs. Translatability was associated with evolutionary conservation, with ancient ORFs being more likely to be translated compared to evolutionarily recent ORFs as determined by phylostratigraphy (p<0.001, two-way ANOVA, Fig. 1g, Supplementary Table 15). ORFs predicted to encode proteins < 50 amino acids were less likely to yield a detectable protein, although this may be explained by the deleterious effect of fusing a 14-amino acid V5 tag to a very small protein.

**Figure 1:**
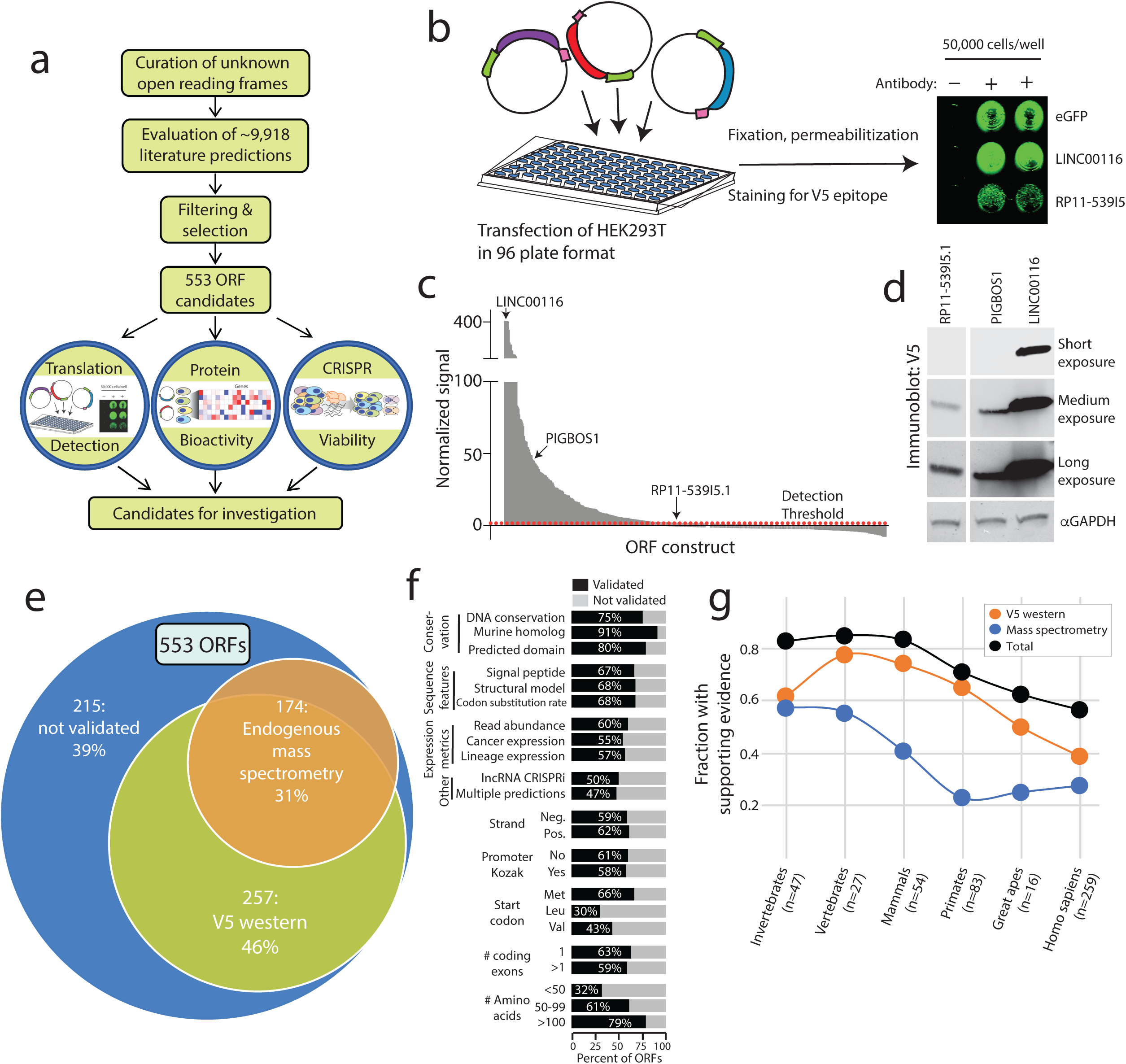
Identification of translated unannotated or unstudied open reading frames. **a)** A schematic overview of the research project. **b)** The experimental set-up for *in vitro* detection of protein translation by transfection of V5-tagged cDNAs into HEK293T cells followed by in-cell western blotting. **c)** In-cell western blot signal for each ORF. Values are the average of three replicates. **d)** Immunoblot correlates for three ORFs identified by in-cell western blotting, marked in panel **c**. **e)** An overview of biological support for translation of a subset of ORFs. **f)** Subgroup analyses of ORF biological features demonstrating fractions of ORFs supported by ectopic V5 translation assays, mass spectrometry or both. **g)** The fraction of ORFs supported by evidence of translation across major epochs in evolutionary time. Evidence of translation shown as the fraction of ORFs with V5 western blot signal, endogenous mass spectrometry peptides, and the summation of both.

Since the majority of non-canonical ORFs have evidence of translatability, we next asked whether such translation was associated with biological activity. To address this, we expressed the 553 ORFs in each of four cell lines (MCF7, A549, A375, HA1E), and then performed RNA expression analysis using the L1000 platform^34^ (Fig. 2a), which monitors the expression of 978 mRNAs. Ectopic expression of 401 ORFs (73%) yielded a reproducible gene expression consequence, of which 237 ORFs induced a high transcriptional activation score (*tas*) indicating marked cellular changes^34^ (Fig. 2b, Supplementary Fig. 3 and Supplementary Table 16). In comparison, 81% of 2,283 canonical protein-coding genes yielded a gene expression consequence in this assay, indicating that the frequency of biological activity of known genes and unannotated ORFs is similar (Fig. 2b). To exclude the possibility that the observed transcriptional signature was due simply to overexpression of the RNA, we mutated translational start sites and repeated the L1000 profiling. In 48 of 51 (94%) cases, the perturbational response was lost when translation was prevented, indicating that the biological effect was indeed mediated by a protein, and not a non-coding RNA (Fig. 2c-f, Supplementary Fig. 4, Supplementary Tables 17-18).

**Figure 2:**
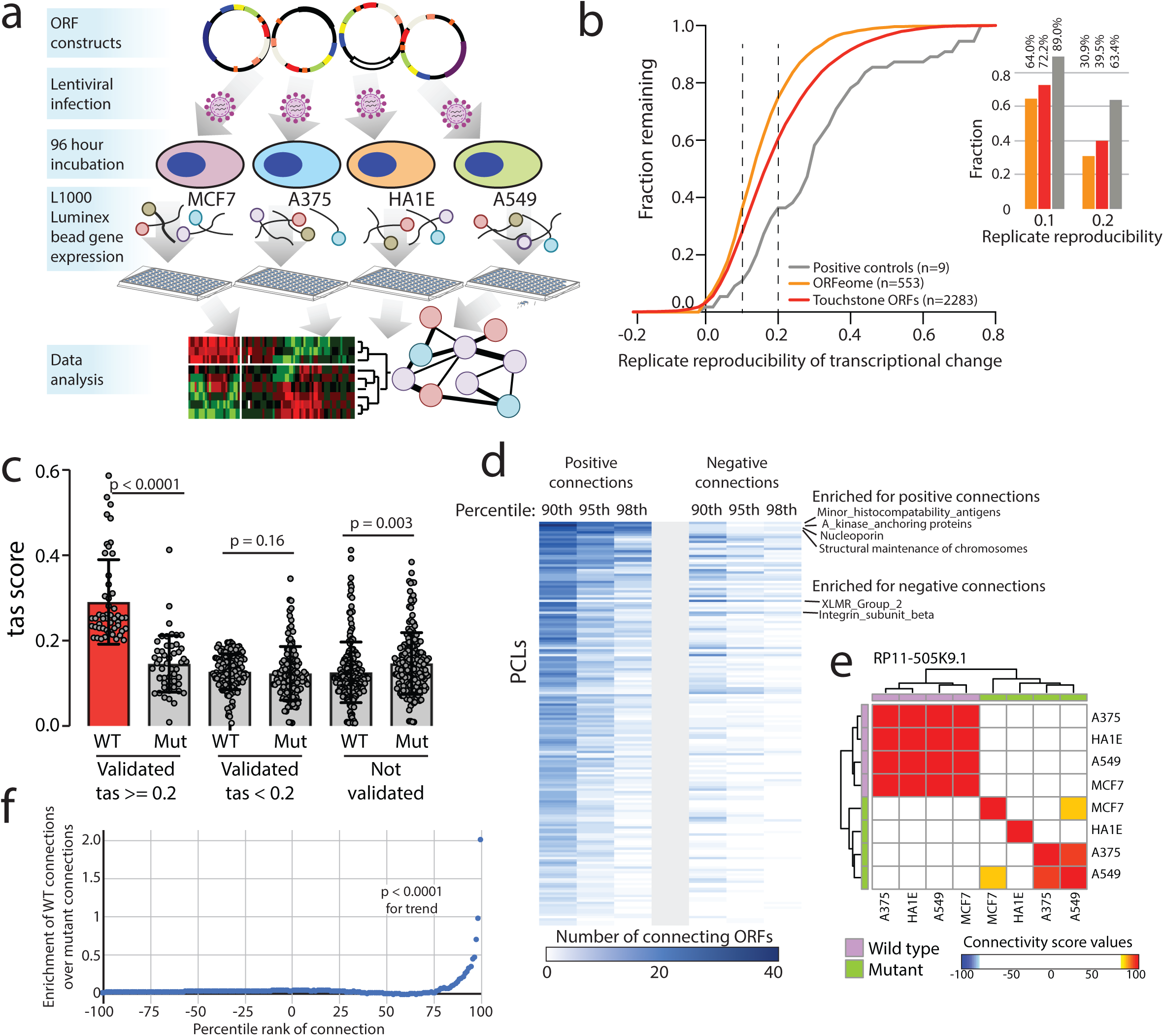
Defining bioactive ORFs through gene expression profiling. **a)** A schematic showing the experimental set-up. Briefly, ORFs were individually transduced into 4 cell lines and expression was profiled 96 hours after infection using the L1000 platform. **b)** The fraction of ORFs resulting in transcriptional perturbation compared to all profiled known genes and assay positive controls. *Inset at the right,* a barplot enumerating the percentage of ORFs in each group with a transcriptional signature above the indicated reproducibility threshold. **c)** A barplot showing the strength of transcriptional perturbation following expression of the indicated groups of wild-type or mutant ORF constructs. P value by Wilcoxon test. Error bars represent standard deviation. **d)** A heatmap showing the number of ORFs demonstrating positive or negative connections with individual Perturbational Classes (PCLs) at the indicated percentile rank. **e)** An example of *RP11-505K9.1* showing the high concordance of connectivity signatures when the wild type ORF is expressed compared to the ORF with mutated translational start sites. **f)** Bland-Altman analysis demonstrating enrichment of high-ranking connectivity values following expression of wild type ORFs compared to mutant ORFs. P value by Wilcoxon test.

The transcriptional responses observed following ORF expression could conceivably be a consequence of overexpression of the transgene. To address the functional relevance of endogenous expression of these ORFs, we performed CRISPR/Cas9 loss-of-function viability screens in 8 cancer cell lines using a guide RNA library targeting the 553 ORFs (Fig. 3a, Supplementary Fig. 5a and Supplementary Table 19). Knock-out of 57 of the 553 ORFs (10%) demonstrated a growth inhibitory effect (Fig. 3b,c, Supplementary Tables 20-21). Of these, 31 (54%) impaired survival of all 8 cell lines, whereas 26 (46%) displayed selective dependency (Supplementary Fig. 5b-e).

**Figure 3:**
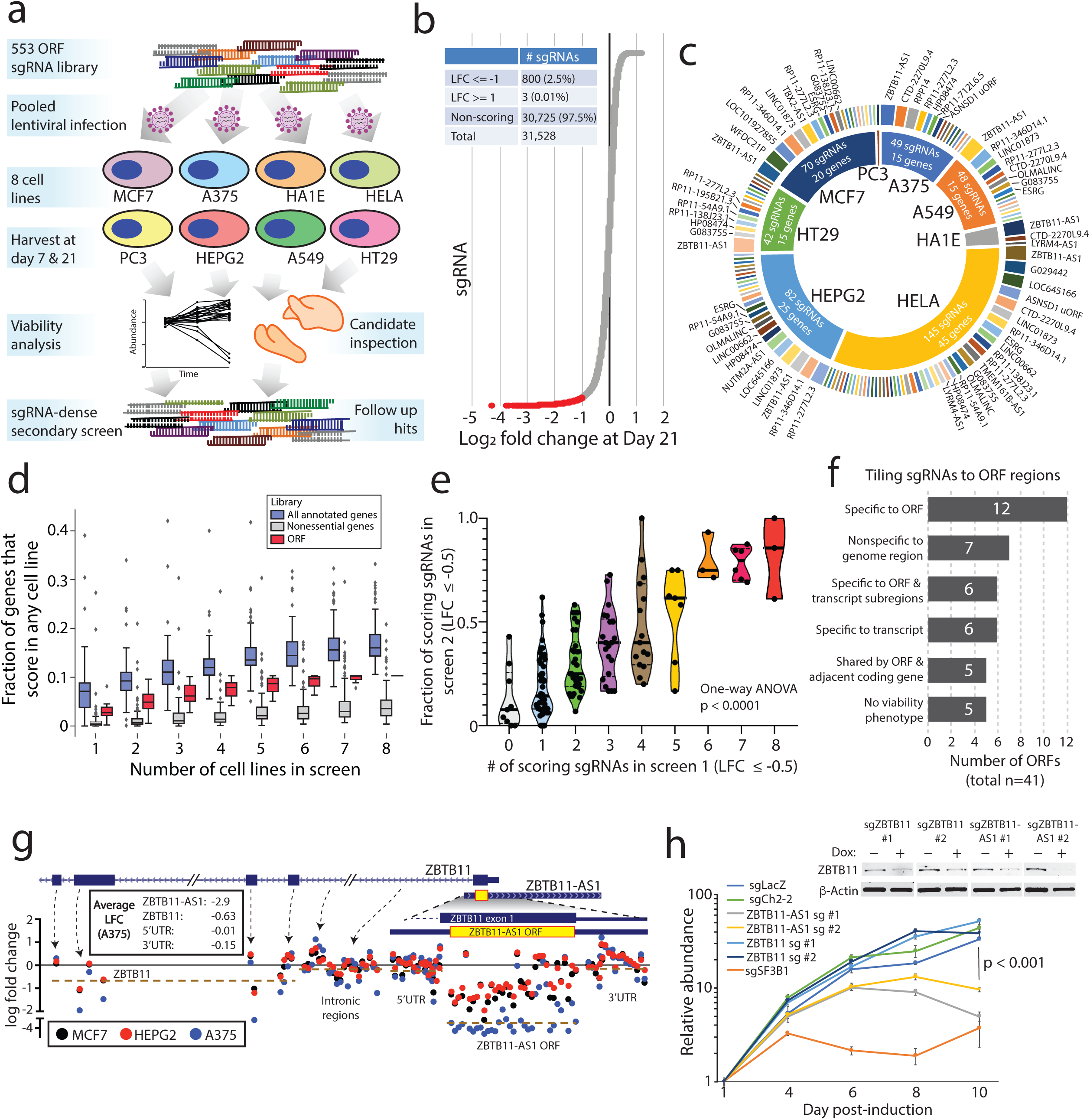
CRISPR screening to identify unknown ORFs implicated in cancer cell viability. **a)** A schematic showing the experimental design, including a primary screen in 8 cancer cell lines and a secondary screen in 3 cancer cell lines. **b)** The distribution of sgRNA depletion at day +21 following lentiviral infection in the CRISPR screen across 8 cell lines. 2.5% of sgRNAs were identified as depleted in a particular cell line with a log2 fold change of <= −1. **c)** The distribution of nominated ORFs. For each cell line, the inner circle, the number of sgRNAs with a log2 fold change of <= −1, and the number of nominated genes are shown. The outer circle shows the ORFs nominated in that cell line, with the ORFs ranked by the number of supporting sgRNAs. The thickness of the outer circle boxes reflects the number of sgRNAs supporting that ORF’s nomination. Only ORFs nominated with >= 2 sgRNAs are shown. **d)** A boxplot showing the fraction of annotated genes, new ORF genes, and RNAi-defined nonessential genes that score as a vulnerability gene in the indicated number of cell lines. Each data point represents a unique cell line. The cell lines for ORF genes represent the cell lines used in this study. For annotated genes, the randomly selected cell lines from the Dependency Map were used. Box plots represent median with interquartile ranges. **e)** The correlation between the number of sgRNAs producing a viability phenotype in the primary and the fraction of sgRNAs producing a viability phenotype in the secondary screen. P value by a one-way ANOVA. **f)** A barplot showing the number of ORFs with each category of viability phenotype in the tiling sgRNA CRISPR screens. **g)** An example of *ZBTB11* and *ZBTB11-AS1* for tiling CRISPR data, showing enhanced cell killing when the *ZBTB11-AS1* ORF is knocked-out. Each data point represents a sgRNA. Data points are color-coded for the indicated cell lines. **h)** Individual CRISPR knockout experiments in a doxycycline-inducible Cas9 HeLa cell line using two sgRNAs targeting exclusively *ZBTB11* or two sgRNAs targeting both the *ZBTB11-AS1* ORF and *ZBTB11*. The line plot shows cell viability measured by cellular ATP following induction of Cas9 activity with 2ug/mL doxycycline. sgLacZ and sgCh2-2 are non-cutting and cutting negative controls, respectively, and sgSF3B1 is a pan-lethal positive control. The inset western blot shows ZBTB11 protein abundance upon induction of Cas9. P value by a two-tailed Student’s t-test. Error bars represent standard deviation.

To compare these data to knock-out of canonical proteins, we analyzed the Cancer Dependency Map (www.depmap.org) for the viability effects of 553 randomly selected genes. Among canonical proteins, 17% demonstrated a viability effect in 8 randomly chosen cell lines, compared to approximately 10% for the non-canonical ORFs (Fig. 3d, Supplementary Fig. 5f-g), indicating a surprisingly similar frequency of dependencies between known genes and non-canonical ORFs. These results were validated both in a secondary CRISPR screen of 147 ORFs (Fig. 3e, Supplementary Fig. 5h-i, Supplementary Tables 22-24), as well as individually-performed CRISPR assays for selected ORFs (Supplementary Fig. 6 and Supplementary Table 27).

Because the viability effects from knock-out of non-canonical ORFs could be explained by loss of a regulatory region in the genome rather than the protein itself, we subjected 41 ORFs to dense tiling of sgRNAs across the genomic locus of each ORF. Only 7/41 (17%) genomic regions demonstrated non-specific viability loss suggestive of a regulatory region of the genome. For 18/41 ORFs (44%), the viability effect mapped exclusively to predicted coding exons or the coding region as well as adjacent nucleotides in the transcript, which may reflect sites of translational regulation or sgRNAs generating indels that also impact the ORF (Fig. 3f, Supplementary Fig. 7, Supplementary Table 29). Interestingly, in several cases, a novel ORF overlapped with an annotated protein-coding gene, but it is the novel ORF that best explained the knock-out phenotype (Fig. 3g). As examples, we observed that ORFs arising from *CTD-2270L9.4* and *ZBTB11-AS1*, which overlap coding exons of *COG7* and *ZBTB11*, respectively, both demonstrated markedly more dramatic viability phenotypes using sgRNAs that target the novel ORF compared to adjacent sgRNAs that target only the known, parent ORF (Fig. 3g-h, Supplementary Fig. 7b). These findings were supported by Cancer Dependency Map data in which sgRNAs targeting both the novel and the known ORFs had a more pronounced phenotype than sgRNAs targeting only the known ORF (Supplementary Fig. 8). Taken together, we conclude that a surprisingly high proportion of non-canonical ORFs exhibit a viability phenotype upon knock-out, and that prior CRISPR vulnerability screens may be confounded by cryptic, novel ORFs arising from the same genomic locus.

We next noted that 13 ORFs scored highly in all three high-throughput assays supporting translation, bioactivity and CRISPR vulnerability (Fig. 4a), suggesting that they may have particularly important biological roles. Among these, we especially focused on *G029442* (*LA16c-380H5.3* in GENCODE) because its knock-out resulted in selective cancer cell killing (1 of 8 cell lines killed), and it is highly expressed in several human cancer types (Fig. 4b and Supplementary Fig. 9). We subsequently renamed this gene *GREP1* (Glycine-Rich Extracellular Protein-1) for reasons elucidated below.

**Figure 4:**
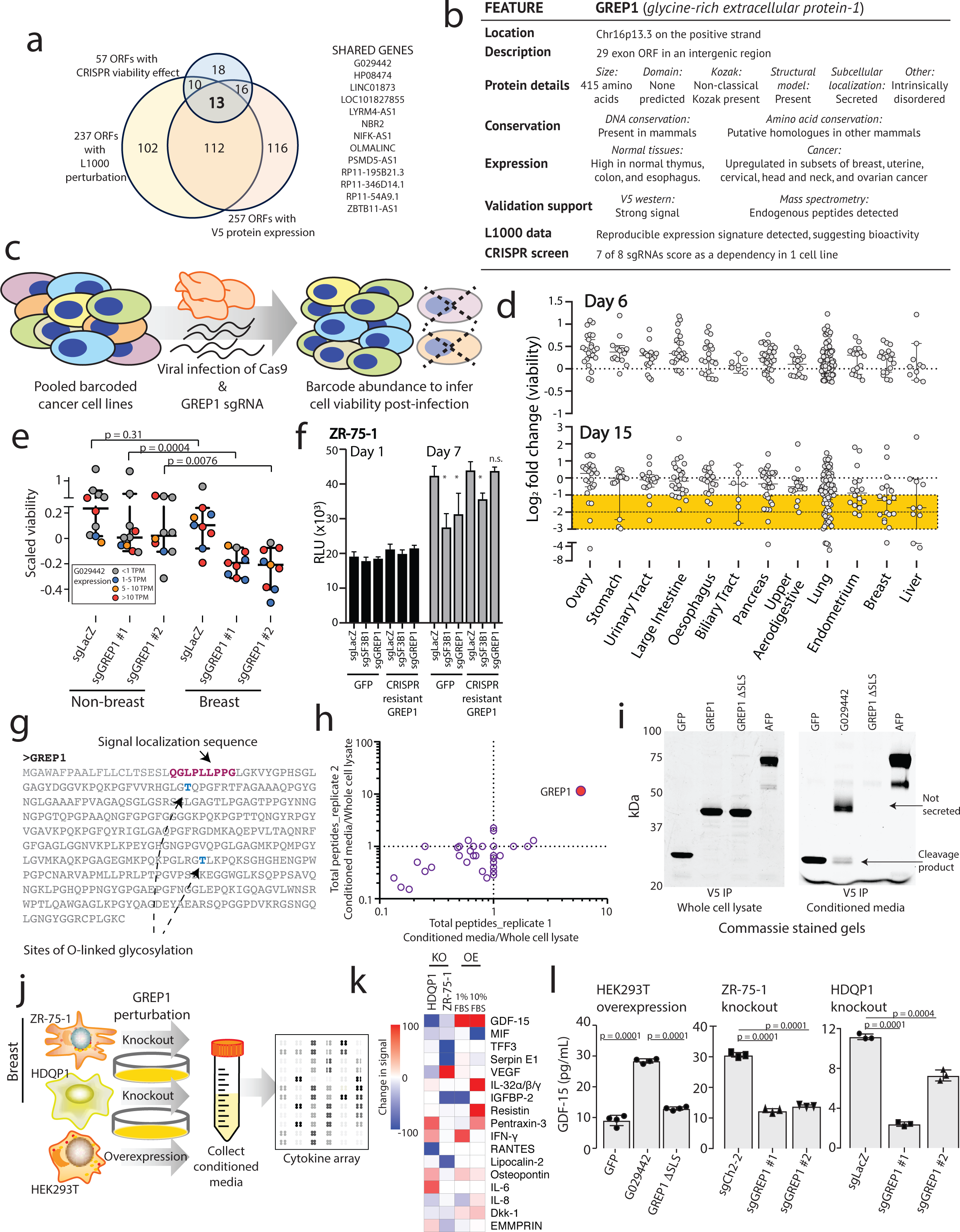
Characterizing *GREP1* as a cancer dependency gene in breast cancer. **a)** Nomination of candidate ORFs with evidence for protein translation, gene expression effect, and CRISPR phenotype. **b)** A table summarizing the characteristics of the *GREP1* gene. **c)** A schematic showing the overview of pooled CRISPR screening. **d)** Log2 fold change abundance of cancer cell lines at Day 6 and Day 15 following pooled CRISPR screening. Cell lineages are ranked based on the median log2 fold change at Day 15. Each data point represents a unique cell line. **e)** Individual CRISPR validation experiments for *GREP1* in a panel of non-breast (n=10) and breast (n=9) cell lines. Data are scaled so that 0 reflects the sgCh2-2 negative cutting control and −1 reflects the degree of viability loss from the sg*SF3B1* positive control. Data were obtained 7 days after lentiviral infection. P value by a Mann-Whitney test. **f)** Rescue of the CRISPR phenotype with overexpression of a CRISPR resistant *GREP1* construct and not GFP. An asterisk (*) indicates a P value < 0.05. P values by a two-tailed Student’s t-test. **g)** The GREP1 amino acid sequence with the signal localization sequence and the sites of glycosylation indicated. **h)** Immunoprecipitation followed by mass spectrometry of HEK293T conditioned media and whole cell lysate following ectopic expression of a pool of V5-tagged ORFs. The x and y axes represent the total number of MS peptides detected in two independent experiments. **i)** Expression of V5-tagged GREP1 or a truncated GREP1 lacking the N-terminal signal localization sequence in HEK293T cells. Cell lysates or conditioned media were subjected to V5 immunoprecipitation and then protein was visualized by Commassie stain. **j)** Experimental design for secreted cytokine profiling following *GREP1* perturbation. **k)** A heatmap showing individual cell line changes in cytokine abundance following *GREP1* perturbation. Cytokines are ranked according to the average of the absolute value of signal change for each cell line. **l)** Validation of *GDF15* modulation upon *GREP1* perturbation by ELISA in the indicated cell lines. P value by a two-tailed Student’s t-test. All error bars represent standard deviation.

To systematically explore the importance of *GREP1* in cancer cell viability, we infected a pool of 486 barcoded human cancer cell lines with a single lentivirus harboring both Cas9 and a guide RNA targeting *GREP1* (Fig. 4c, Methods). Because lentiviral infection rates vary across the cell lines, we focused our analysis on the 263 cell lines yielding highest quality data (Supplementary Fig. 10a-g, Supplementary Tables 30-31, Methods). *GREP1* knockout resulted in preferential loss of viability in certain cell lineages, most notably breast cancer (Fig. 4d). We validated these pooled screening results with knock-out and rescue experiments for *GREP1* in breast and non-breast cell lines, which confirmed a striking breast cancer viability phenotype that correlated with *GREP1* mRNA expression (Fig. 4e-f, Supplementary Fig. 10h, and Supplementary Fig. 11). Finally, *GREP1* expression was higher in human breast cancers compared to normal breast tissue (p = 1.4 x 10^-10^) (Supplementary Fig. 9c) and was associated with decreased patient survival in breast but not colon cancer patients (Supplementary Fig. 11c,d). Together, these data implicate *GREP1* as a previously unrecognized, prognostic breast cancer vulnerability gene.

To explore the function of GREP1, we noted the presence of a signal localization sequence for extracellular secretion, as well as sites of glycosylation documented by mass spectrometry (Fig. 4g, Supplementary Table 32). We confirmed that ectopic expression of a GREP1 fusion protein with a C-terminus V5 epitope tag, but not an N-terminal truncation mutant lacking the signal localization sequence, was indeed secreted as well as cleaved into a smaller product (Fig. 4h-i, Supplementary Fig. 11e-f, and Supplementary Table 33). Analyses of the GREP1 amino acid sequence revealed a conserved, glycine-rich, and intrinsically disordered protein (Supplementary Fig. 12a-c), characteristics that resemble some members of the extracellular matrix^35^. As expected, immunoprecipitation of ectopically expressed GREP1 from cell culture media followed by mass spectrometry revealed a strong enrichment for extracellular matrix proteins, including fibronectin and EMILIN2 (Supplementary Fig. 12d-h, Supplementary Table 34).

To establish the cellular consequence of GREP1 expression, we examined the impact of *GREP1* knock-out and overexpression on other secreted proteins by testing a panel of 102 secreted proteins by antibody arrays across 3 cell lines (Fig. 4j). The metabolic cytokine GDF15^36, 37^ demonstrated the most dramatic change, with *GREP1* knockout resulting in decreased GDF15 levels and GREP1 overexpression resulting in increased GDF15 levels (Fig. 4k,l and Supplementary Fig. 13a,b). Consistent with this, *GREP1* and *GDF15* expression were correlated across multiple tumor types in the TCGA database (Supplementary Fig. 13c,d). To determine whether GDF15 was functionally important in cancer cells’ requirement of GREP1 for survival, we tested the effect of *GREP1* knock-out in the presence and absence of recombinant GDF15. Remarkably, supplementation of recombinant human GDF15 rescued the loss of viability caused by *GREP1* loss of function (Supplementary Fig. 13e,f). The fact that GDF15 only partially rescues *GREP1* knock-out in some cell lines suggests that there may be additional mechanisms downstream of GREP1 that regulate cell survival (Supplementary Fig. 13g). While GDF15 has been previously implicated in a number of cancer phenotypes including chemoresistance^38, 39^, immune evasion^40^, cellular survival and invasiveness^41, 42^, its regulation by GREP1, which itself is a cancer dependency, is entirely new.

Despite the fact that the human genome was sequenced 18 years ago, the precise number of protein-coding genes in the genome remains a point of controversy. Our systematic sampling of over 550 uncharacterized ORFs provides the first experimental evidence that a substantial proportion of such ORFs are functional. Importantly, we establish that approximately 10% of the ORFs in our dataset were required for the survival of cancer cells, a rate only about half that observed for known, canonical proteins. Although our dataset represents a curated list of ORFs rather than a random sampling of all possible ORFs, these experiments suggest that further investigations of unannotated ORFs in cancer and other disease states will likely yield new insights. Since estimates of such ORFs now exceed 5,000^43^, our data predict the existence of thousands of functional, unannotated ORFs in the human genome. Overall, our work develops an approach for understanding uncharacterized ORFs through systematic interrogation and establishes that these non-canonical proteins represent a rich source of untapped biology that is deserving of further investigation.

## Methods

### Data statement

No statistical methods were used to predetermine sample size. The experiments were not randomized and the investigators were not blinded to allocation during experiments and outcome assessment.

### Cell lines and reagents

All parental cell lines were obtained from the American Type Culture Collection (ATCC, Manassas, VA). Cas9-derived cell lines were obtained from the Broad Institute. Cell lines were maintained using standard media and conditions. In brief, cell lines derived from ZR-75-1, HCC1806, HCC1954, HCC202, T47D, HT29, HCC15, KYSE410, KYSE510, SNU503, SW837, HCT116, AU565 and MDA-MB-231 were maintained in RPMI 1640 (Invitrogen, Carlsbad, CA, Carlsbad, CA) supplemented with 10% FBS and 1% penicillin-streptomycin in a 5% CO2 cell culture incubator at 37°C. Cell lines derived from HDQP1, BT-474, JIMT1, A375, A549, MIAPACA2, MCF7, HEK293T and MDA-MB-468 were maintained in DMEM supplemented with 10% FBS (Invitrogen, Carlsbad, CA) and 1% penicillin-streptomycin (Invitrogen, Carlsbad, CA) in a 5% CO2 cell culture incubator. GFP-positive Cas9-derived cell lines were drug-selected using 2ug/mL blasticidin.

For stable knockout cell lines, ZR-75-1 Cas9 and HDQP1 Cas9 expressing cells were infected with lentivirus for the indicated sgRNAs which had been cloned into the LentiGuide-Puro plasmid (plasmid #52963, Addgene) with 4ug/mL of polybrene. 16 hours after transduction, cells were selected with cell culture media containing 2ug/mL of puromycin. Cells were maintained in puromycin-containing media for 72 hours before transitioned back to standard culture media. Stable GREP1-overexpressing cell lines were generated in ZR-75-1 and CAMA-1 cells by infecting with a sgRNA-resistant *GREP1* cDNA construct and selecting with 350ug/mL of hygromycin for 96 hours, before transitioning back to standard culture media.

### RNA isolation; cDNA synthesis; and qPCR experiments

Total RNA was isolated using Qiazol and an miRNeasy Kit (Qiagen, Hilden, Germany) with DNase I digestion according to the manufacturer’s instructions. RNA integrity was verified on an Agilent Bioanalyzer 2100 (Agilent Technologies, Palo Alto, CA). cDNA was synthesized from total RNA using Superscript III (Invitrogen, Carlsbad, CA) and random primers (Invitrogen, Carlsbad, CA). Quantitative Real-time PCR (qPCR) was performed using Power SYBR Green Mastermix (Applied Biosystems, Foster City, CA) on a Thermo QStudio FLX Real-Time PCR System (Thermo Fisher Scientific, Waltham, MA). GAPDH was used the housekeeping control gene. The relative quantity of the target gene was completed for each sample using the ΔΔCt method by the comparing mean Ct of the gene to the average Ct of the geometric mean of the indicated housekeeping genes. The primer sequences are listed below:

GREP1 3’UTR-forward: AGCCTCCAAATGGCTATGGAC

GREP1 3’UTR-reverse: CTCGAGGCCACCATTAAAAC

GREP1 ORF-forward: CTGGATATCCGGCTGGAGATG

GREP1 ORF-reverse: ATTGCTGCCTCTCTTCACGTC

GAPDH-forward: TGCACCACCAACTGCTTAGC

GAPDH-reverse: GGCATGGACTGTGGTCATGAG

Beta-actin forward: AAGGCCAACCGCGAGAAG

Beta-actin reverse: ACAGCCTGGATAGCAACGTACA

Fibronectin forward: GAGAAAATGGCCAGATGATGA

Fibronectin reverse: AATGGCACCGAGATATTCCTT

Emilin2 forward: AACAAAGTGCTGGTGAACGAC

Emilin2 reverse: CTCTCCTGTACCCAGCGGTAT

ZBTB11-AS1 forward: CCGTTTTTACGTTTGAGACTCC

ZBTB11-AS1 reverse: ATGTAAATGGGCTGTCTCTGGT

HP08474 forward: GTGTAAAGAGGTCCTGGGACAG

HP08474 reverse: GCACTCCAGTCTAGACGACACA

RP11-54A9.1 forward: TTGGTGAGATGTTCCTTGAGC

RP11-54A9.1 reverse: CTCCACTTCACTGTCGGTCTC

G083755 forward: ATCCCATCTGAGTGCTTACCAA

G083755 reverse: CATGCATAATCTCCTTCCCTGC

OLMALINC forward: AGGAACATCTTGCCAATTTCA

OLMALINC reverse: TGTGGATCTTCAGTTGCTTCA CTD-2270L9.4

forward: AGTCGTTGGCCGTTACCATA CTD-2270L9.4

reverse: CTTCCCAGGCTCAAGCAAT

ASNSD1 uORF forward: ACAATTCGACCCCACACAAG

ASNSD1 uORF reverse: GGTTAGAAAGTTCATCCACCACA

RP11-277L2.3 forward: CTACGTGGGGCTGGAAATAC

RP11-277L2.3 reverse: CCCTTCCCAGTTCTCTGACC

### Selection process for candidate ORFs

Candidate ORFs were collated via manual curation from 25 published studies and one in-house analysis of ribosomal profiling data (Z. Ji, personal communication). Published studies are listed in Supplementary Table 1. Data types included were 14 studies with mass spectrometry data, 6 studies with ribosomal profiling data, 4 studies with computational ORF predictions, and 1 study with both mass spectrometry and ribosomal profiling data. In total, there were 9,918 candidate ORFs among which 4,433 unique Ensembl transcripts were represented. From these, pseudogenes, N-terminal extension ORFs, ORFs of known proteins with new predicted exons, and alternative reading frame ORFs located entirely within the genomic nucleotides of an annotated protein were removed from consideration. 553 high-priority ORFs were selected from the ORFeome. The 553 ORFs were then manually curated according to the following metrics as described. See Supplementary Figure 1 for an overview.

#### DNA conservation

An ORF was considered to have high DNA conservation if the average PhastCons score for 100 placental mammals was ≥0.9 for the entire ORF.

#### Murine homolog

Murine homologs were defined by the Slncky program^44^.

#### Cancer association

ORFs were searched in the Pubmed database for associations with the word “cancer”. Additionally, cancer-associated transcripts in the MiTranscriptome were also queried.

#### Lineage association

ORFs were searched in the NIH Roadmap Epigenome Project data^45^, which transcriptionally profiled human embryonic stem cells before and after differentiation into mesenchymal stem cells, neural progenitor cells, trophoblast-like cells, or meso-endoderm.

#### High read coverage

ORFs were stratified if they had a read/length ratio of ≥1.0 in available ribosomal profiling data

#### Codon substitution rate

ORFs were stratified is they had a codon PhyloCSF decibans score (29 mammal alignment) of ≥ 5.0 averaged across the whole ORF transcript.

#### Protein domain structure

ORFs were analyzed via the NCBI Conserved Domain finder. ORFs were domain structures were an e value confidence score of < 0.01 are indicated.

#### Multiple overlapping ORF predictions

Published ORF predictions from 25 large datasets were integrated^5, 6, 16–19, 22^ to nominate 203 ORFs with at least 2 publications supporting their existence (Supplementary Tables 1 & 2).

#### Predicted ORF CRISPR phenotype

Data from a CRISPR interference screen of lncRNAs were employed^46^. Of 492 lncRNA hits nominated in that study, there were 312 hits with GENCODE identifiers were could be further evaluated. Of those 312, there were 292 unique GENCODE identifiers, which were manually reviewed. GENCODE identifiers overlapping ORFs in this ORFeome are indicated.

#### Upstream and downstream ORFs

We used candidates from Ji et al.^5^ and considered conserved upstream and downstream ORFs between mouse and human, as defined by an inter-species alignment with an E value of < 0.0001. We evaluated ORFs with all of the following attributes: a Ka/Ks conservation ratio of < 0.5, an ORF length of ≥25 amino acids, an ATG start site, and the ORF was non-overlapping with the annotated ORF. 49 dORFs and 195 uORFs met these criteria and were manually reviewed to select candidates included in the ORFeome.

#### Signal peptide

All ORFeome ORFs were analyzed by SignalP version 4.1 using standard default settings^47^ and a D-score of ≥0.450 to nominate ORFs with a classical signal localization sequence.

#### Structural modeling

All ORFeome ORFs that are ≥40 amino acids were analyzed by the Phyre2 structural domain prediction software using default settings^48^. To distinguish ORFs enriched for structural models, we generated a random amino acid sequence library of 500 random 150-mer amino acid sequences with a methionine start codon. We derived a structural model score of (%ORF alignment to the structural model) * (%confidence of the model). A structural model score of 0.175 was used to maximally differentiate ORFeome ORFs from random amino acid sequences.

#### Overall ORF confidence score

Each criteria as above in addition to mass spectrometry peptide evidence (see below) was given a binary score of 1 if the criterion was met by the ORF or 0 if not met by the ORF. The ORF confidence score was the summation of these binary scores.

### Identification of small open reading frames in proteomics datasets

A fasta database containing the amino acid sequences of the 553 ORFs was appended to a reference protein database (UCSC RefSeq) and used to search peptide mass spectra of datasets acquired or analyzed in our laboratory. These datasets predominantly comprised of studies conducted by the Clinical Proteomics Tumor Analysis Consortium (CPTAC) (Supplementary Table 12). Raw mass spectrometry (MS) data were analyzed in Spectrum Mill MS Proteomics Workbench v6.0 (Agilent Technologies, Santa Clara, CA) employing search parameters specific for each project. Detailed descriptions of search parameters such as enzyme definition and specificity or the number of types variable modifications included in database search can be found in the corresponding publications (Supplementary Table 12).

Peptide-spectrum matches (PSMs) to the ORF database were identified by automatically parsing through database search results generated by Spectrum Mill Software using an inhouse developed R-script. Only PSMs validated by target-decoy based false-discovery (FDR) estimation were used for subsequent analysis. To further minimize the possibility of false positive identifications, we required a minimal Spectrum Mill PSM score of 8 which roughly translates to a minimum of eight identified fragment ions in the MS/MS spectrum. All PSMs meeting the criteria described above are listed in Supplementary Table 12.

### Phylostratigraphy analysis

All ORFs with an amino acid length of >= 40 amino acids was analyzed as described previously^49, 50^, using TimeTree^51^ to identify evolutionary strata. Using a BLASTP e-value threshold of 10^-3^ and a maximum number of 200,000 hits, we identified the phylostratum in which each ORF appeared. For clarity, we aggregated results into the following evolutionary eras: Invertebrates (phylostrata 1-9, including cellular organisms through Craniata, ∼540 millions of years ago (Mya)), Vertebrates (phylostrata 10-17, including Vertebrata through Amniota (312 Mya)), Mammals (phylostrata 18 - 22, including Mammalia through Euarchontoglires (95 Mya)), Primates (phylostrata 23-27, including Primates through Hominoidae (20 Mya)), Great apes (phylostrata 28-30, including Hominidae through Homo), and Humans (phylostratum 31, including Homo sapiens).

### Generation of the ORFeome library

Initial prototype plasmids were generated in the pLX_TRC307 vector backbone designed for prior ORF studies^52^, obtained from the Broad Institute Genomic Perturbation Platform (Broad Institute, Cambridge, MA, USA), by PCR-amplification from cell line cDNA (HeLa, HEK293T, K562, or MCF7). PCR products were gel-purified (Qiagen, Hilden, Germany), cloned into the non-directional Gateway PCR8 vector (Invitrogen, Carlsbad, CA) as an entry vector, and shuttling to pLX_TRC307 using LR clonase II (Invitrogen, Carlsbad, CA) according to the manufacturer’s instructions. pLX_307 is a Gateway-compatible expression vector where E1a is the promoter of the ORF and SV40 is the puromycin resistance gene (details available at https://portals.broadinstitute.org/gpp/public/resources/protocols). Following technical optimization of the insert sequence to include a barcode sequence following the V5 tag, the final ORF construct design is as follows:

vector backbone -> ORF sequence lacking stop codon -> c-terminus V5 sequence (GGTAAGCCTATCCCTAACCCTCTCCTCGGTCTCGATTCTACG) -> Triple stop codon (TAGTAATGA) -> P1 primer site (TCTTGTGGAAAGGACGA) -> Barcode sequence -> AC (linker sequence) -> vector backbone.

Following the ORF sequence, each construct therefore had the additional sequence:

GGTAAGCCTATCCCTAACCCTCTCCTCGGTCTCGATTCTACGTAGTAATGATCTTGTGGAAAGGACGA_BARCODE_AC

The ORFeome library was then generated via insert synthesis and cloning of unique plasmid inserts consisting unique barcodes (Supplementary Table 22) by a commercial vendor (GenScript, Piscataway, NJ) in arrayed barcoded tube format. Each plasmid therefore had a barcode sequence flanked by common PCR primer pair for amplification of a 233bp amplicon, where the sense primer was located in the ORF insert and the antisense primer was located in the plasmid backbone as follows:

P1 Sense primer: TCTTGTGGAAAGGACGA

P2 Antisense primer: TTAAAGCAGCGTATCCACATAGCGT

### Generation of paired mutant ORFs

The 85 mutant constructs employed and identical plasmid insert construct as detailed above with the following modifications: the putative ORF start codon was mutated to GCG (encoding alanine), and all internal in-frame ATG codons (encoding methionine) were mutated to GCG to reduce the chance of internal initiation of translation. Constructs were generated via commercial gene synthesis (GenScript, Piscataway, NJ).

### In cell western blotting

HEK293-T cells were plated at a density of 20,000 cells per well in a 96 well black plate format to minimize autofluorescence. 6 to 8 hours after plating, cells were transiently transfected with 0.1 ug of an individual plasmid with Fugene HD reagent (Promega, Madison, WI). 48 hours later, cell culture media was removed, and cells were fixed for 20 minutes with 150 uL of 3.7% formaldehyde solution in 1x phosphate-buffered saline at room temperature with no shaking. Fixing buffer was removed and cells were washed five times with 200 uL PBS containing 0.1% Triton X-100 (Sigma-Aldrich, St. Louis, MO) for permeabilization. Following this, cells were blocked with 150uL of Odyssey Blocking Buffer (LI-COR, Lincoln, NE) for 90 minutes at room temperature on a plate shaker. Cells were then treated with anti-V5 antibody at 1:200 concentration in Odyssey Blocking Buffer or no-antibody control wells. Cells were incubated with the primary antibody overnight at 4°C. The next day, the primary antibody was removed and cells were washed five times with 200uL PBS containing 0.1% Triton X-100 as above. Then, 50uL secondary antibody was applied at 1:1000 dilution and samples were incubated for 1 hour with gentle shaking and protection from light. Afterwards, wells were washed five times with 200uL PBS containing 0.1% Tween20 (Sigma-Aldrich, St. Louis, MO). After the final wash, plates were blotted on tissue paper to remove excess wash buffer and immediately scanned on a LI-COR Odyssey system using the 800nm light channel to image and quantify anti-V5 abundance.

### Analysis of in cell western data

First, a preliminary dilution series was performed with decreasing amounts of transfected plasmid and decreasing numbers of HEK293T cells plated per well (Extended Data Figure 3). This was performed for two high-expressing plasmids that were verified by western blot (eGFP and LINC00116), and one low-expressing verified plasmid (RP11-539I5.1). Using eGFP and RP11-539I5.1 we defined a dynamic range for the assay (Extended Data Figure 3) by normalizing V5 800nm light signal to the plate background. This defined a threshold above which signal was reproducibly detected even in low-expressing plasmids when transfected into 1,000 plated HEK293T cells.

Then, for the full ORFeome library, all plasmids were run in biological triplicate on 3 unique 96 well plates for in cell western analysis. Each plate was normalized by median-centering raw 800nm signals within each plate to minimize variance in plate background. Normalized 800nm signals were then averaged across replicates. Plasmids with averaged signal above the previously defined threshold based on RP11-539I5.1 expression were considered to generate a protein by V5 tag detection.

### *In vitro* transcription/translation

50 ORFs were selected at random from the ORFeome library for synthesis of the ORF insert lacking a V5 tag and containing a 5’ T7 promoter sequence. This tag-free insert was cloned into pUC57 plasmid. 1.0 mcg of linearized purified plasmid were subjected to wheat germ extract *in vitro* transcription/translation systems employing the non-radioactive Transcend tRNA system according to manufacturer’s instructions (Promega, Madison, WI). 10 of 50uL from the reaction volume was then heat-denatured in the presence of DTT and protein bands were detected by SDS-PAGE gel electrophoresis using a Tris-Glycine 10-20% gel (Thermo Fisher Scientific, Waltham, MA).

### Immunoblot Analysis

Cells were lysed in RIPA lysis buffer (Sigma-Aldrich, St. Louis, MO) and allowed to homogenize on ice for 30 minutes after lysis. Cell debris was removed by centrifugation for 15 minutes at 13,200 RPM and the debris pellet was discarded. 1x HALT protease inhibitor (Thermo Fisher Scientific, Waltham, MA) was added to lysate supernatants. Protein abundance was quantified by the bicinchoninic acid (BCA) method using and bovine-specific albumin standard curve for normalization of protein abundance. Aliquots of each protein extract were boiled in LDS sample buffer, size fractionated by SDS-PAGE at 4°C by Tris-Glycine 10-20% gels, and transferred onto nitrocellulose membranes with pre-cast gels via the iBlot-2 system (Thermo Fisher Scientific, Waltham, MA). The membrane was then incubated at room temperature for 1-2 hours in LICOR Odyssey blocking buffer and incubated at 4°C with the appropriate antibody overnight. Following incubation, the blot was washed 4 times with 1x TBS with 0.1% Tween20 and incubated with fluorophore-specific IRDye secondary antibodies (LI-COR, Lincoln, NE) and imaged on a LI-COR Odyssey machine.

For conditioned media western blots, conditioned media of GFP- or G029442-expressing HEK293T cells was concentrated using 3kDa size exclusion filter tubes (Millipore, Burlington, MA) by a factor of 5-fold. Then, 1x HALT protease inhibitor was added to the sample. Samples were kept at 4°C and not frozen to preserve protein fidelity. Western blots were then performed as detailed above.

### Antibodies used

| Antibody | Species | Monoclonal/<br>Polyclonal | Dilution | Catalogue<br>Number | Vendor | Conditions |
| --- | --- | --- | --- | --- | --- | --- |
| V5 (D3H8Q) | Rabbit | Monoclonal | 1:2000 | 13202S | Cell<br>Signaling<br>Technology | 4C<br>overnight |
| ZBTB11 | Rabbit | Polyclonal | 1:1000 | A303-240A-M | Bethyl<br>Laboratories | 4C<br>overnight |
| Beta-Actin | Mouse | Monoclonal | 1:4000 | A5316 | Sigma-Aldrich | 4C<br>overnight |
| Goat anti-mouse<br>secondary | Goat | N/A | 1:5000 | 926-32210 | LI-COR | 20C for 1<br>hour |
| Goat anti-rabbit<br>secondary | Goat | N/A | 1:5000 | 926-68021 | LI-COR | 20C for 1<br>hour |

### Non-denaturing western blot

Non-denaturing western analysis was performed using the NativePage system (Thermo Fisher Scientific, Waltham, MA). In brief, HEK293T cells were transfected with plasmid encoding GREP1. 72 hours after transfection, conditioned media was collected and cellular debris was removed via centrifugation and filtering of the media. Protease inhibitor was added to the conditioned media for preservation. Conditioned media was then prepared with 4x NativePAGE sample buffer without heat, detergents, or reducing agents. For comparison, conditioned media was also prepared using 4x NativePAGE sample buffer and also 1% SDS and 10% NuPAGE sample reducing agent (Thermo Fisher Scientific, Waltham, MA) followed by boiling at 95°C for 5 minutes. Samples were then run on a NativePAGE Novex Bis-Tris gel using NativePAGE running buffer and NativePAGE 20x Cathode Buffer according to manufacturer’s instructions. Proteins were transferred to a PVDF membrane after membrane activation with isopropanol using a semi-dry system of 7V for 30 minutes at room temperature. After blocking for 1 hour at room temperature in Odyssey Blocking Buffer, membranes were treated with rabbit anti-V5 antibody at a 1:2000 dilution (Clone D3H8Q, #13202S, Cell Signaling Technology, Danvers, MA) overnight at 4°C, then washed 4 times in 1x TBS-Tween, and treated with anti-rabbit HRP secondary antibody at a 1:10000 dilution. Chemilluminence was achieved with SuperSignal West Dura Extended Duration Substrate (Thermo Fisher Scientific, Waltham, MA), and images were developed with CareStream Kodak BioMax light film (Kodak, Rochester, NY).

### Lentivirus production for L1000 experiments

Complete details of standard virus production pipelines can be found at the Broad Institute Genetic Perturbation Platform website https://portals.broadinstitute.org/gpp/public/.

Virus was produced in arrayed 96 well plates via triple transfection of HEK293T cells with each packaging vector (100 ng), envelope plasmid (10 ng), and each respective pLX317 plasmid (100 ng). Lentiviral-containing supernatants were harvested at 32-56 hours post-transfection and stored in polypropylene plates at −80°C until use.

### Cell lines and lentiviral transduction for L1000 expression profiling

A549 and A375 cells were cultured in RPM1 media supplemented with 10% FBS and 1% penicillin/streptomycin. MCF7 and HA1E cells were cultured in DMEM media supplemented with 10% FBS and 1% penicillin/streptomycin. To perform L1000 HT gene expression profiling, cells were robotically seeded (40uL per well) into 384 well plates. Optimized seeding densities were 250 cells per well (MCF7), 400 cells per well (A549, A375 and HA1E). Twenty-four hours post-seeding cells were spin-infected in the presence of polybrene (4 ug/mL for A549 and HA1E and 8 ug/mL for MCF7 and A375 cells). The plates were then centrifuged for 30 minutes at 1,178 g at 37°C. The supernatant was robotically removed and replaced with fresh media 3 hours (A549) or 24 hours post-infection (A375, MCF7, HA1E) and cells cultured for an additional 72 hours till assay.

Infections were carried out in 5 replicates, 3 of which were used for L1000 assay and 2 used for assessing the infection efficiency. To assess infection efficiency, cells were treated with or without puromycin selection (1.5 ug/mL) 24 hours post-infection, and cell viability was determined using CellTiterGlo (Promega, Madison, WI) after 72 hours of selection. For the remaining plates, media was removed 96 hours post-infection, and the cells were lysed with the addition of TCL buffer (Qiagen, Hilden, Germany). Plates were then sealed and stored at −80 °C until gene expression profiling.

### L1000 experimental design

Two 384 well plates of perturbational ORFs were designed for cell treatment prior to L1000 profiling, each containing 352 unique ORFs, negative control ORFs, internal technical controls, and untreated wells. Plate format can be found in Extended Data Figure 4. In each plate, 346 wells were devoted to treatment ORFs, and ten to ORFs targeting L1000 landmark genes were included for positive control purposes. These positive control wells would later be assessed for targeted gene z-score (≥ 2) and targeted gene rank (computed relative to the expression levels of that same gene across the assay plate). Control genes included were *ACAA1*, *ACD*, *AURKB*, *BMP4*, *CBR1*, *CCDC90A*, *CDK6*, *CSNK1A1*, *ETV1*, and *SOX2*. Genes were selected for overall for high baseline expression levels in the lines profiled and previous reproducibility in the L1000 assay. Additionally, 16 wells of negative control ORFs targeting BFP, EGFP, or HCRED were added. Each plate also contained 12 untreated wells.

Cell lines MCF7, HA1E, A549, and A375 were chosen to represent a diversity of tissue types and also to match CMap cell lines that had been profiled extensively in the past and were constituents of the CMap reference database *Touchstone*^34^.

### L1000 data processing

Detailed protocols for the L1000 assay are provided at https://clue.io/sop-L1000.pdf. Each plate was profiled 96 hours after infection. Antibiotic selection was not employed, and each plate was processed using the standard L1000 data processing pipeline which has been described elsewhere^34^. Briefly, mRNA was initially captured using 384-well oligo dT-coated Turbocapture plates; after removing lysate and adding a reverse-transcription mix containing MLLV, the plate was washed and a mixture of both upstream and downstream probes (each containing a gene-specific sequence and a universal primer site) for each of the 978 (“Landmark”) genes measured was added. The probes were first annealed to cDNA over a six hour period, and then ligated together to form a PCR template. After ligation, Hot Start Taq and universal primers were added to the plate, the upstream primer was biotinylated to allow for later staining with streptavidin-pycoerythrin, and the PCR amplicon was hybridized to Luminex microbeads using the complementary and probe-specific barcode on each bead. After overnight hybridization, the beads were washed and stained with streptavidin-pycoerythrin and Luminex FlexMap 3D scanners were used to measure each bead independently, reporting bead colour, identity, and fluorescence intensity of the stain. Fluorescence intensity of the stain values were then converted into median intensity values for each of the 978 measured genes using a deconvolution algorithm (resulting in “GEX” level data). These GEX data were then normalized relative to a set of invariant genes, and subsequently quantile normalized (resulting in “QNORM”) level data. An inference model was applied to the QNORM data to infer gene expression changes for a total of 10,174 genes, which corresponds to the “BING” (Best INferred Genes) genes we report below. Next, expression values of each individual well was converted to robust z-scores by z-scoring gene expression relative to corresponding expression across the entire plate population; these z-scored differential expression gene signatures were lastly replicate collapsed to a single differential expression vector per treatment, which we term a signature (and “MODZ” level data).

### L1000 quality control

All samples profiled passed internal technical L1000 assay quality control measures described elsewhere^34^. Additionally, all samples included passed an internal fingerprinting algorithm that verifies the identity of cell lines on L1000 plates by comparing quantile-normalized gene expression data in each will to a ranked reference library of over 1000 cancer cell lines; samples are defined as passing if their Spearman correlation to their respective reference profile is higher than equivalent correlation values to all other reference cancer profiles. Additionally, 67% of positive control ORFs included had a replicate correlation of 0.25 or greater and induced a z-score of 2 or greater in their target gene. Notably, ORFs targeting *CNSK1A1* represented the majority of poorly performing positive control ORFs. Positive control ORFs that showed high transcriptional activity (TAS) also clustered together (Extended Data Figure 4c).

### Measures of L1000 signature bioactivity

Each perturbagen’s transcriptional activity was represented using a Transcriptional Activity Score (TAS), which has been described in depth elsewhere^34^. Briefly, TAS is computed as a geometric mean of signature strength (SS; or, the number of landmark (n=978) genes in a signature with absolute z-score greater than or equal to 2) and replicate correlation (RC; or, the 75th quantile of all pairwise Spearman correlations between replicate level z-score profiles):
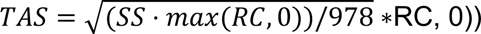. Signatures were considered to be bioactive if they had a TAS score of 0.2 or higher, which represents the value at which 95% of negative control wells fall below^34^.

### L1000 signature queries

Each MODZ-level signature profiled was queried both against the other L1000 signatures in the dataset and against the Connectivity Map dataset that has been published and described elsewhere^34^. Similarity values between these signatures was assessed using a percentile score derived from a normalized weighted connectivity score (WTCS). Briefly, WTCS is a similarity measure based on the weighted Kolmogorov-Smirnov enrichment statistic (ES) described previously^53^ and is computed as follows for a given query gene set pair (q_up, q_down) and a reference signature r:

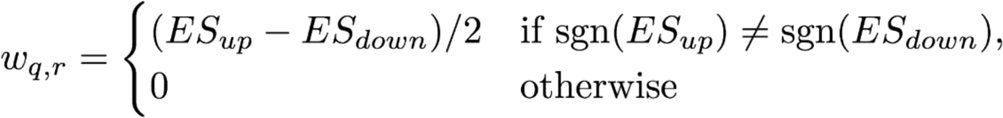

Where ES_up is the enrichment of q_up in r and ES_down is the enrichment of q_down in r. WTCS ranges between −1 and 1, and is positive for signatures that are positively related, negative for the converse, and near zero for unrelated signatures.

WTCS is then normalized to allow for comparison of connectivity scores across cell and perturbagen types; this normalization is similar to that used in Gene Set Enrichment Analysis and accounts for differences in connectivity that may occur across such covariates. Given a vector of WTCS values from a query, normalization occurs as follows:

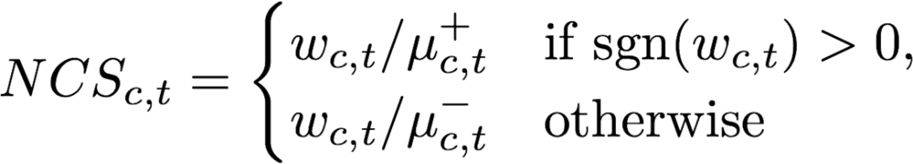

Where NCS_c,t_, w_c,t_, U+_c,t_, U-_c,t_ are the normalized connectivity scores, raw WTCS, and signed means (the mean of the positive and negative values evaluated separately) of the WTCS values within the subset of signatures corresponding to cell line c and perturbagen type t, respectively.

Lastly, NCS scores are converted to percentile scores accounting for whether the connectivity between the queried (“q”) and reference signature (“r”) are significantly different from that observed between r and other queries. This is done by comparing each observed NCS value ncs_q,r_ between the query q and a reference signature r to a distribution of NCS values representing the similarities between a reference compendium of queries (Q_ref_) and r. This procedure results in a standardized measure we refer to as Tau (τ) that ranges from −100 to +100 and represents the percentage of queries in Q_ref_ with a lower |NCS| than |ncs_q,r_|, adjusted to retain the sign of ncs_q,r_ and relies on the following formula:

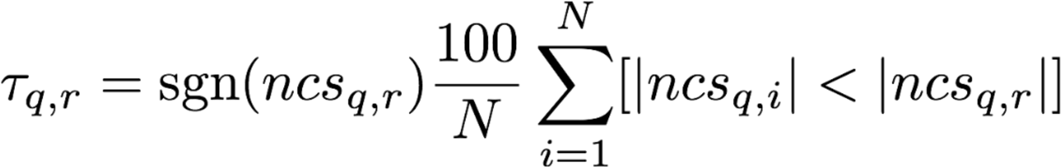

Where ncs_q,r_ is the normalized connectivity score for signature r w.r.t query q, ncs_i,r_ is the normalized connectivity score for signature r relative to the i-th query in Q_ref_ (a set of query signatures obtained from exemplar signatures of perturbagens matching the cell line and perturbagen type of signature r) and N is the number of queries in Q_ref_.

### L1000 software packages used

L1000 data were analysed using the ‘tidyverse’ suite^54^ of R packages (v1.2.1) and the ‘cmapR’ package^55^ (v1.0.1) in R v3.5.0 (R Core Team 2018).

### CRISPR sgRNA design

sgRNAs for the ORFs in this study were designed using the Broad Institute GPP sgRNA designer for S. Pyogenes Cas9 against genome coordinates for the GRCh38 assembly of the human genome (https://portals.broadinstitute.org/gpp/public/analysis-tools/sgrna-design). Only exonic coding regions for the ORFs were used. A maximum of 8 unique sgRNAs were employed per gene. If fewer than 8 were nominated due to small gene size and lack of available PAM sites, then all nominated sgRNAs were used. If more than 8 sgRNAs were nominated, then the top 8 ranked sgRNAs were used according the the Broad Institute GPP sgRNA designer pick analysis. For the secondary CRISPR screen, 147 ORFs were tested. These were chosen to include all ORFs with a viability phenotype in the primary screen in the appropriate cell lines (A375, MCF7, HEPG2), as well as additional ORFs that did not have viability phenotype.

For tiling sgRNA analyses, additional nominated sgRNAs for each ORF were selected. Also, we selected sgRNAs to putative 3’UTR, 5’UTR, and promoter regions (defined as within 1000 basepairs of the transcript start site). A maximum of 16 sgRNAs were designed for each region. If there were multiple UTR exons, then a maximum of 16 sgRNAs were designed for each UTR exon. Intronic sgRNAs were used were available and limited to 6 sgRNAs per intron. sgRNAs for adjacent protein coding genes were also employed as indicated, and designed in an identical manner. The number of sgRNAs for adjacent coding genes and various genome regions is detailed in Supplementary Tables 23 and 24.

### Determination of infection conditions for CRISPR pooled screens

Optimal infection conditions were determined in each cell line in order to achieve 30-50% infection efficiency, corresponding to a multiplicity of infection (MOI) of ∼0.5 - 1. Spin-infections were performed in 12-well plate format with 3 × 10^6^ cells each well. Optimal conditions were determined by infecting cells with different virus volumes with a final concentration of 4 ug/mL polybrene. Cells were spun for 2 hours at 1000 g at 30°C. Approximately 24 hours after infection, cells were trypsinized and 2×10e5 for A375, HT-29, and PC-3 cells; 1.5×10e5 for A549 and HeLa cells; 3×10e5 for HepG2 cells; 5×10e5 for MCF-7 cells from each infection were seeded 2 wells of a 6-well plate, each with complete medium, one supplemented with the appropriate concentration of puromycin (1.5 ug/mL for A375; 2 ug/mL for A549, MCF7 and PC-3; 1 ug/mL for HeLa, HA1E, HepG2, and HT-29). Cells were counted 4-5 days post selection to determine the infection efficiency, comparing survival with and without puromycin selection. Volumes of virus that yielded ∼30 - 50% infection efficiency were used for screening.

### Primary and secondary CRISPR pooled proliferation screens

The lentiviral barcoded library used in the primary screen contains 5235 sgRNAs, which includes an average of 8 guides per gene and 500 non –targeting control guides. The validation library contains 6996 sgRNAs targeting selected regions of the smORFs. Genome-scale infections were performed in three replicates with the pre-determined volume of virus in the same 12-well format as the viral titration described above, and pooled 24 h post-centrifugation. Infections were performed with enough cells per replicate, in order to achieve a representation of at least 1000 cells per sgRNA following puromycin selection (∼1.5×10e7 surviving cells). Approximately 24 hours after infection, all wells within a replicate were pooled and were split into T225 flasks. 24 hours after infection, cells were selected with puromycin for 7 days to remove uninfected cells. After selection was complete, 1.5-2×10e7 of cells were harvested for assessing the initial abundance of the library. Cells were passaged every 3-4 days and harvested ∼21 days after infection. For all genome-wide screens, genomic DNA (gDNA) was isolated using Midi or Maxi kits for the validation screens gDNA was isolated using and Midi kits according to the manufacturer’s protocol (Qiagen, Hilden, Germany). PCR and sequencing were performed as previously described^56, 57^. Samples were sequenced on a HiSeq2000 (Illumina, San Diego, CA). For analysis, the read counts were normalized to reads per million and then log_2_ transformed. The log2 fold-change of each sgRNA was determined relative to the initial time point for each biological replicate.

### Analysis of CRISPR screening data

CRISPR data was analyzed as log2 fold change values computed between the day 21 timepoint and the input plasmid DNA. A log2 fold change of <= −1 was defined as a scoring sgRNA which was depleted in the analysis. In the primary screen, a gene with at least 2 sgRNAs with a log2 fold change of <= −1 in at least 1 cell line was defined a putative vulnerability hit. Because the vast majority of genes in the primary screen had 8 sgRNAs per cell line, genes could be compared against each other with this metric. In the secondary screen, because the number of sgRNAs for each gene varied, a scoring candidate was defined as a gene in which at least 10% of the sgRNAs had a log2 fold change of <= −1, and there were at least 2 sgRNAs with a log2 fold change of <= −1 in at least 1 cell line. sgRNAs were also analyzed via STARS and CERES scores as previously described^56, 58^.

### Analysis of CRISPR tiling screen

Log2 fold change values for each sgRNA at day 21 of the screen were considered as above. sgRNAs were then grouped into their respective genomic region (e.g. UTR, ORF exon, adjacent gene exon, intron). The mean log2 fold change for each region was computed. A mean log2 fold change of <= −1 was considered to be a scoring hit. Genes were then classified in the following manner according to the viability affect of the sgRNAs: “specific to ORF” if only the ORF region sgRNAs scored; “specific to ORF and transcript subregion” if the ORF sgRNAs and sgRNAs to only one other region of the RNA transcript scored; “specific to transcript” if sgRNAs to any part of the ORF or RNA transcript scored, but not sgRNAs to introns or genomic regions; “shared with adjacent gene” if the ORF and an annotated adjacent protein coding gene both scored; “nonspecific to the genome” if sgRNAs to any part of the genomic region, intron, RNA transcript or ORF all demonstrated depletion.

### Comparison of CRISPR screen data with Project Achilles

For each gene of ORF in each of the eight cell lines used in the primary ORF CRISPR screen, knockout was determined to produce depletion if at least two guides produced at least 50% depletion from initial abundance after RPM normalization. The file “Achilles_logfold_change” in DepMap_public_19Q4 was used for Achilles screens (available at https://depmap.org/portal/download). To determine the expected number of genes or ORFs that deplete in any cell line given N cell lines, all possible subsets of N lines were selected and the number of genes with at least one depleted line were counted. For a negative control, this process was repeated in Achilles screens using only genes proposed as non-essential by previously published RNA interference data^59^, to generate a null distribution.

### GREP1 annotation analysis and expression data

*GREP1* annotation status was evaluated using the indicated historical versions of the GENCODE database with graphic visualization of the locus. In cell lines, *GREP1* expression was evaluated through Cancer Cell Line Encyclopedia data for *LINC00514* (NR_033861.1), a RefSeq annotation which incorporates the first portion of *GREP1*. CCLE data was downloaded from https://portals.broadinstitute.org/ccle.

### Pooled GREP1 knockout

For the pooled *GREP1* CRISPR knockout assay, we used a pool of 486 barcoded, adherent human cancer cell lines developed at the Broad Institute^60^. The cell line pool was grown in RPMI1640 media supplemented with 10% FBS. sgRNAs used for this experiment were non-cutting control sgLacZ (AACGGCGGATTGACCGTAAT), cutting control sgChr2 (GGTGTGCGTATGAAGCAGTGG), sg*GREP1* #1 (ACTCAAAATGGCTATAGACC), and sg*GREP1* #2 (AGGCTTTAGAGGGGACATGA). On Day 0, the cell line pool was plated in 6 well plates at 400,000 cells per well in 3mL of cell culture media. 24 hours later, using an all-in-one Cas9/sgRNA plasmid, the cell line pool was infected with each lentivirus at an MOI of 10; lentivirus was concentrated prior to use to obtain a concentration of >1e7 particle/ml. Cells were also treated with 4ug/mL polybrene in 2mL/well for the lentiviral infection, and spun at 2250rpm for 1 hour at 37°C. 24 hours after transduction, cells were split from 1 well in a 6 well plate into two T25 flasks; at this time the baseline cell DNA lysate was harvested as a “no infection” control. 72 hours after infection, cell culture media was changed and puromycin selection was started at a concentration of 1ug/mL puromycin. Thereafter, cell culture media was changed every 72 hours and cells were expanded as needed into T75 and T175 flasks. Pooled cell line DNA was collected from the input plasmid pool, on day +6 as an early timepoint, and day +15 as a late timepoint to assess for dropout of cell line. At each sample timepoint, cells were counted and 2e6 cells were removed for lysis for DNA. For lysis, cells were pelleted, washed in PBS, and genomic DNA was extracted with the DNA Blood and Tissue Kit according to manufacturer’s instructions (Qiagen, Hilden, Germany). The remainder to the cells not taken for lysis were re-seeded into T75 and T175 flasks for continuing cell growth.

For sequencing, timepoint DNA was subjected to PCR using universal barcode primers. PCR products were run on a 2% agarose gel to confirm amplicon size. Then 10uL from each PCR product was pooled, purified with AMPur beads (Beckman Coulter, Brea, CA). DNA concentration was measured via Qubit fluorometric quantification (Thermo Fisher Scientific, Waltham, MA) and DNA was sequenced on a NovaSeq (Illumina, San Diego, CA) at the Genomics Platform at the Broad Institute.

### Analysis of pooled *GREP1* knockout sequencing data

Cell line abundance was calculated based on cell line barcode detection by next generation sequencing as previously described^60^. To analyze the pooled *GREP1* CRISPR knockout data, we first calculated the theoretical number of cells in each well at each timepoint based on the experimental measurements of the total number of cells and the number of cells removed for sequencing. We accounted for these removed cells by scaling the measured number of cells at a given timepoint by the ratio of the total number of cells at the previous timepoint to the number of reseeded, or continued, cells from the previous timepoint.

Next, for quality control, we computed the purity of each sample as the percentage of the read counts mapping to cell lines not in the pool. We removed samples with lower than 95% purity. We also filtered out cell lines with fewer than 12 reads in more than one replicate of either of the two negative control conditions, LacZ and Chr2. The conservative threshold of 12 was determined from the minimum number of counts at which we are able to distinguish between that number of counts and half that number, at a confidence level of 0.05, under a Poisson distribution.

Then, we added a pseudocount of 1 to each of the read counts and normalized the updated read counts by the library size and the theoretical total cell count. We define the log fold-change of a cell line in a sample as the log2-transform of the ratio of the normalized read count of the cell line in the sample to the normalized read count of the cell line at day 0. Finally, we define the viability as the difference between the log fold-change in the cell line and treatment of interest and the average of the log fold-changes in the cell line and the two negative controls.

Next, we developed a series of data processing steps to empirically improve the quality of the dataset (see Supplementary Fig. 10). First, we excluded cell lines believed to be puromycin resistant based on the criterion of positive viability in the puromycin, no-virus condition. These filters resulted in a viability dataset of 400 out of 486 cell lines. Then, we removed cell lines that exhibited excessive lentiviral toxicity given the high MOI used for this experiment. This left 320 cell lines. Next, we eliminated cancer type cohorts with less than or equal to 5 cell lines, due to insufficient numbers for analysis, leaving 294 cell lines. Lastly, we calculated the number of cell lines per cancer cohort that expressed *GREP1* above a minimal threshold, and excluded cohorts with insufficient expression as any change in those cohorts may be spurious due to population shifts in the cell line pool or off-target effects.

### Patient outcomes analysis for *GREP1*

Expression data for *GREP1* in the TCGA samples was acquired from the MiPanda publicly available tool using the LA16c-H380H5.3 gene annotation as a query^61^. Data for the GDC TCGA Breast Cancer and GDC TCGA Colon Cancer datasets were used. LINC00514 expression was extracted as a proxy for *GREP1* given that LINC00514 is a fragment of the longer gene. Overall survival was also extracted for these datasets. Kaplan-Meier curves and statistical significance via Log-rank P value were generated using GraphPad Prism8 software, with a p value of < 0.05 being considered statistically significant.

### GREP1 copy number analysis

CCLE copy number data from the 2013-12-03 segmentation was downloaded from https://depmap.org. Data for LINC00514 (283875) was used as a proxy for *GREP1* given overlapping genomic regions. Copy number data was then aggregated by cell line lineage.

### CRISPR validation experiments

Cells were plated in 96-well plates and allow to grow for 4-8 hours prior to infection with the indicated sgRNA or treatment condition. 1,000 - 5,000 cells per well were plated depending on the cell line. sgRNAs were obtained from the Broad Institute Genomic Perturbation Platform (Broad Institute, Cambridge, MA, USA) or from direct synthesis into the BRDN0003 backbone via commercial vendor (GenScript, Piscataway, NJ). sgRNA sequences are listed below:

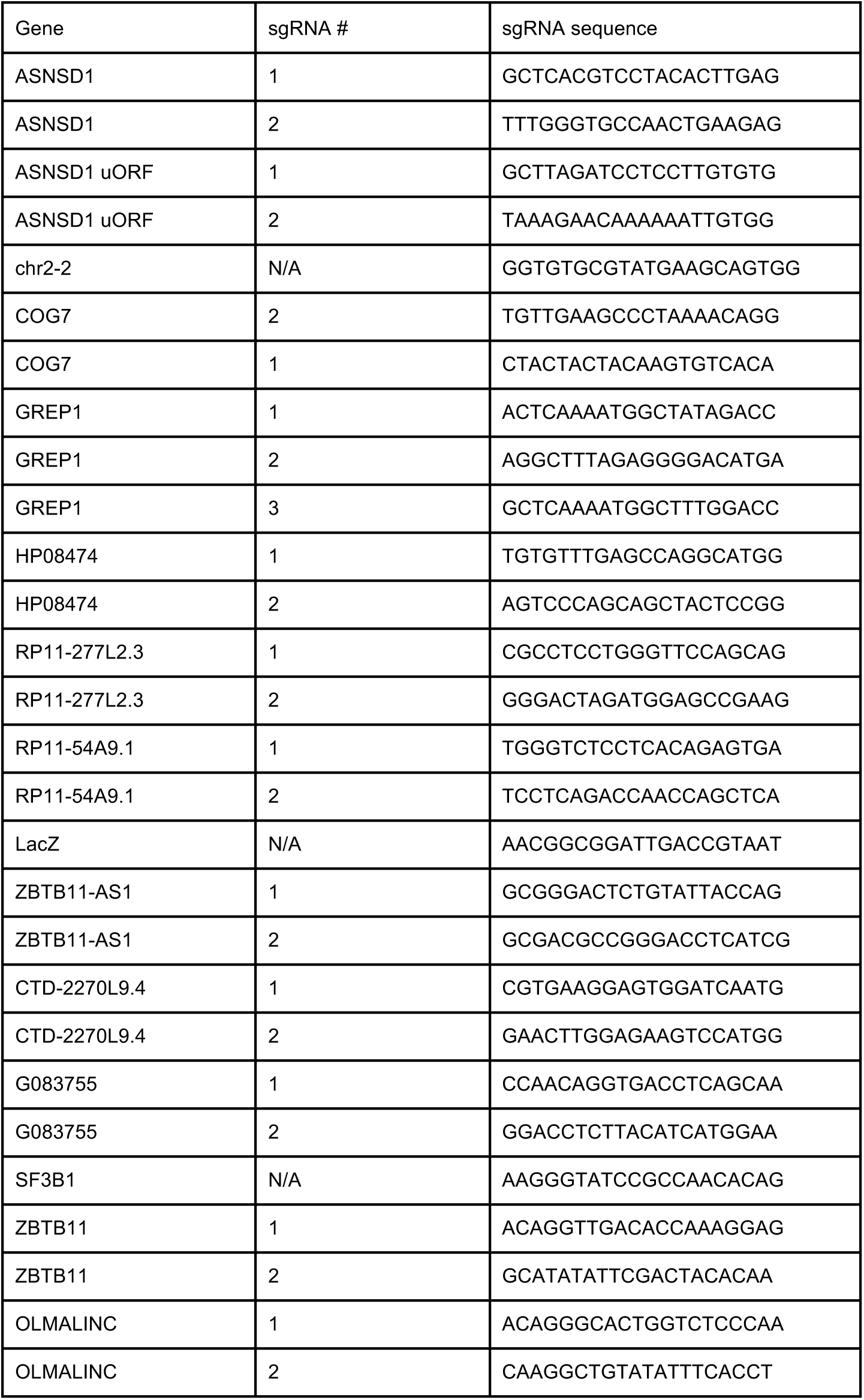

All sgRNAs were sequenced and verified. After sequence verification, constructs were transfected with packaging vectors into HEK-293T with Fugene HD (Sigma-Aldrich, St. Louis, MO). After plating, cells were then infected with sgRNA lentivirus to achieve maximal knockout but without viral toxicity. 16 hours after infection, cells were selected with 2ug/uL puromycin (Invitrogen, Carlsbad, CA) for 48 hours. Cell viability was measured CellTiter-Glo reagent (Promega, Madison, WI) was measured at 16 hours post-transfection for a baseline assessment, and additional timepoints as needed. For stable knockout cell lines, cells were plated at equal densities and cell viability was measured by CellTiter-Glo every 24 hours as indicated.

### GREP1 overexpression rescue experiments

For CRISPR rescue experiments, Cas9-derivatized cell lines were infected with lentivirus GFP or GREP1 coding plasmids cloned into the pLX_TRC313 vector, which has EF1a promoter and hygromycin resistance (see https://portals.broadinstitute.org/gpp/public/vector for details). Cells were selected in 350ug/mL of hygromycin for 72 hours prior to transitioning back to standard culture media.

In 96 well plates, 5000 ZR-75-1 derived cells were plated and infected with the indicated sgRNA lentivirus 4-6 hours after plating. 16 hours after infection, cells were selected with 2ug/mL puromycin for 48 hours and grown for 7 days prior to cell viability analysis using CellTiter-Glo reagent.

### Conditioned media rescue experiments

First, on day −2 HEK293-T cells were plated and transiently transfected with GFP and GREP1 with Fugene HD reagent. On day −2, 5000 ZR-75-1 derived cells or 2500 AU565-derived cells were plated in wells of a 96-well plate. On day −1, ZR-75-1 and AU565 cells were switched to serum-free media. On day 0, conditioned media from GFP- or GREP1-expressing HEK293-T cells was cleared of cellular debris via centrifugation and then 100uL of conditioned media was applied to each well. Conditioned media was then refreshed daily and cell viability was determined with the CellTiter-Glo reagent at the indicated time points.

### Immunoprecipitation

HEK293-T cells were transiently transfected with GFP-V5 or GREP1-V5 fusion proteins using OptiMem and Fugene HD (Sigma-Aldrich, St. Louis, MO). 72 hours after transfection, cell culture media was harvested and cell debris was sedimented by centrifugation at 1500 rpm x 5 minutes twice. Resulting cell culture media was concentrated in a 10:1 manner using 3kDa size-exclusion filter (Millipore, Burlington, MA). Concentrated culture media was treated with HALT protease inhibitor. Next, all immunoprecipitation steps were performed on ice or in a 4°C cold room. First, culture media was cleared with Pierce magnetic A/G beads (Thermo Fisher Scientific, Waltham, MA) for 1 hour while rotating at 18-20 rpm. Beads were then discarded and 10% of the media was removed as an input sample and kept at 4°C without freezing. The remained of the culture media was then treated with 50uL of magnetic anti-V5 beads (MBL International, Woburn, MA) and rotated at 18-20 rpm overnight at 4°C. The next day, the supernatant was discarded and beads were washed four times in IP wash buffer (50nM TricHCl, pH 8.0, 150nM NaCl, 2mM EDTA, pH 8.0, 0.2% NP-40, and 1ug/mL PMSF protease inhibitor) with rotation for 10 minutes per wash. After the final wash, beads were gently centrifuged and residual wash buffer was removed. Then, proteins were eluted twice with 2 ug/uL V5 peptide in water (Sigma-Aldrich, St. Louis, MO) at 37°C for 15 minutes with shaking at 1000rpm. The two elution fractions were pooled and samples were prepared with 4x LDS sample buffer and 10x Sample Reducing Agent (Thermo Fisher Scientific), followed by boiling at 95°C for 5 minutes.

One-third of the eluate was then run on a 10-20% Tris-Glycine SDS Page gel and stained with SimplyBlue Commassie stain (Thermo Fisher Scientific, Waltham, MA) for 2 hours. Gels were destained with a minimum of 3 washes in water for at least 2 hours per wash. Bands were visualized using Commassie autofluorescence on the LI-COR Odyssey in the 800nM channel.

Gel lanes were then cut into 6 equal-sized pieces using a sterile razor in sterile conditions, and stored in 1mL of DEPC-treated water prior to mass spectrometry analysis.

### Methods for Protein Sequence Analysis by LC-MS/MS

LC-MS/MS was performed in the Taplin Biological Mass Spectrometry Facility at the Harvard Medical School. Briefly, excised gel bands were cut into approximately 1 mm^3^ pieces. Gel pieces were then subjected to a modified in-gel trypsin digestion procedure^62^. Gel pieces were washed and dehydrated with acetonitrile for 10 min. followed by removal of acetonitrile. Pieces were then completely dried in a speed-vac. Rehydration of the gel pieces was with 50 mM ammonium bicarbonate solution containing 12.5 ng/µl modified sequencing-grade trypsin (Promega, Madison, WI) at 4°C. After 45 min., the excess trypsin solution was removed and replaced with 50 mM ammonium bicarbonate solution to just cover the gel pieces. Samples were then placed in a 37°C room overnight. Peptides were later extracted by removing the ammonium bicarbonate solution, followed by one wash with a solution containing 50% acetonitrile and 1% formic acid. The extracts were then dried in a speed-vac (∼1 hr). The samples were then stored at 4°C until analysis.

On the day of analysis the samples were reconstituted in 5 - 10 µl of HPLC solvent A (2.5% acetonitrile, 0.1% formic acid). A nano-scale reverse-phase HPLC capillary column was created by packing 2.6 µm C18 spherical silica beads into a fused silica capillary (100 µm inner diameter x ∼30 cm length) with a flame-drawn tip^63^. After equilibrating the column each sample was loaded via a Famos auto sampler (LC Packings, San Francisco CA) onto the column. A gradient was formed and peptides were eluted with increasing concentrations of solvent B (97.5% acetonitrile, 0.1% formic acid).

As peptides eluted they were subjected to electrospray ionization and then entered into an LTQ Orbitrap Velos Pro ion-trap mass spectrometer (Thermo Fisher Scientific, Waltham, MA). Peptides were detected, isolated, and fragmented to produce a tandem mass spectrum of specific fragment ions for each peptide. Peptide sequences (and hence protein identity) were determined by matching protein databases with the acquired fragmentation pattern by the software program, Sequest^64^ (Thermo Fisher Scientific, Waltham, MA). All databases include a reversed version of all the sequences and the data was filtered to between a one and two percent peptide false discovery rate. Glycosylated peptides were defined using the A score method as described^65^.

### IP-MS and gene ontology analysis

We analyzed IP-MS data from two independent experiments for V5 immunoprecipitation for GFP-V5 and GREP1-V5 conditioned media. IP-MS data was merged for the two experiments and all proteins with < 2 total peptides were removed to exclude technical artifacts. To the remaining proteins, a pseudocount of 1 was added to ensure a non-zero denominator. Next, fold change of (GREP1+1)/(GFP+1) peptide count was calculated and log10-transformed. Enriched peptides with a (GREP1+1)/(GFP+1) ratio of >= 2 were further analyzed using the Gene Ontology database (http://geneontology.org) for cellular component analysis and corrected false discovery rates were plotted as shown.

### GREP1 disorder analysis

The GREP1 primary amino acid sequence was analyzed via the DISOPRED3 package^66^ on the PsiPred server (http://bioinf.cs.ucl.ac.uk/psipred/) using default settings. Disorder scores were plotted as indicated.

### GREP1 evolutionary analysis

The GREP1 amino acid sequence (ENST00000573315.2_prot) was aligned to non-redundant protein sequences using the NCBI BlastP suite as well as manually aligned to the genomes of the common rat (RGSC 6.0/rn6, July 2014 assembly) and domestic dog (Broad CanFam3.1/canFam3 assembly). Resulting protein hits were then ranked by E-value value and the most significant result was used for each organism. Predicted proteins and low-quality protein assemblies were included in this analysis. Resultant species-specific amino acid sequences were then aligned by the Clustal Omega sequence aligner (https://www.ebi.ac.uk/Tools/msa/clustalo/) and percent similarity to human GREP1 was plotted.

### GREP1 codon usage analysis

We calculated the triplet codon frequency for all triplet codons for the GREP1 amino acid sequence, the whole ORFeome in total, and GENBANK genes by collating all mRNA sequences within these respective groups and calculating the codon usage per group. Each codon usage was normalized to a standard rate of codon usage per 1000 codons. Triplet codons were then collapsed into single amino acids by scaling the codon usage rate to the relative frequency of usage for each codon per amino acid. Aggregate frequency of amino acid representation was then calculated and compared across groups.

### Cytokine profiling array

Cytokine profiling was performed simultaneously using the Human XL Cytokine Array (R&D Systems, ARY022, Minneapolis, MN). Briefly, cell culture media were cleared of cellular debris and Halt protease inhibitor was added as above. Then, cytokine arrays were blocked in 2mL of Array Buffer 6 (blocking buffer) each for 1 hour on a shaker at room temperature. Samples were prepared with 300uL of culture media and diluted with 1200uL of Array Buffer 6. Cytokine arrays were then removed from blocking buffer and incubated with samples overnight at 4°C on a rocker. The next morning array membranes were washed in 20mL 1x Wash Buffer for a total of 3 washes. Then, arrays were placed in 1.5mL of 1x Array Buffer 4/6 (a 1:2 mixture of Array Buffer 4 and Array Buffer 6), and 30uL of reconstituted detection antibody cocktail was added. Samples were incubated for 1 hour at room temperature on a shaker. Subsequently, membranes were washed in 20mL 1x Wash Buffer for a total of 3 washes, and then transferred to 2.0mL of 1x streptavidin-HRP for 30 minutes at room temperature on a shaker, followed by three more washes in 20mL of 1x Wash Buffer. Afterwards, the membranes were blotted on tissue paper to remove excess buffer, and signal was developed with chemiluminescent reagent mix. Images were developed with CareStream Kodak BioMax light film (Kodak, Rochester, NY).

### Cytokine profiling analysis

Immunoblot images of the cytokine arrays were scanned and the signal intensity of all array antibody spots was determined using ImageJ (https://imagej.nih.gov/ij/index.html). Raw data values were then inverted using the formula y = 255 - x, where x is the raw signal intensity. Inverted values were then normalized according to knockout or overexpression experiments. For knockout experiments, signal was analyzed as sgControl - sgGREP1. For overexpression experiments, signal was analyzed as GREP1 - GFP. Then, the absolute value of signal change was averaged across experiments and rank-listed according to the magnitude of average change.

### GDF15 ELISA

The GDF15 Quantikine ELISA kit (R&D Systems, Minneapolis, MN) was used. In brief, cell culture media samples were diluted 1:3 by volume in Diluent RD5-20. To prepare microplate wells, 100uL of Assay Diluent RD1-9 was added to each well. Then, 50uL of standards, controls, or diluted samples were added to a given well. The plates were incubated at 2 hours at room temperature on a horizontal orbital microplate shaker at 500rpm. Wells were then washed four times with 400uL of 1x Wash Buffer for five minutes per wash; after the final wash, plates were inverted and blotted on tissue paper to remove excess. Then, 200uL of Human GDF-15 conjugate was added to each well and the plate was incubated for 1 hour at room temperature on an orbital shaker. Following this, wells were then washed four times with 400uL of 1x Wash Buffer for five minutes per wash; after the final wash, plates were inverted and blotted on tissue paper to remove excess. Then, 200uL of Substrate Solution was added per well, and plates were incubated for 30 minutes at room temperature without shaking and protected from light. Then, 50uL of Stop Solution was added per well and samples were mixed with gentle tapping. The optical density of samples at 450nM and 570nM was determined on a microplate reader within 15 minutes of completion of the protocol. For analysis, background signal from 570nM was subtracted per well from the 450nM signal. Samples were then calculated based on a standard curve to obtain GDF-15 concentration values.

### Correlation of *GREP1* and GDF15 expression

*GREP1*, *GDF15*, *FN1*, and *EMIL2* expression was downloaded via the MiPanda portal^61^ as TPM values. GTex and TCGA samples were used. Spearman rho correlation coefficients and Spearman p values were calculated using GraphPad Prism8 and plotted as shown.

### Recombinant GDF15 experiments

Recombinant human GDF15 (R&D Systems, Minneapolis, MN, catologue number 957-GD-025) was resuspended in water at 10ug/uL. Knockout with sgGREP1 #2 or controls in ZR-75-1 was performed as described above. 24 hours after infection with lentiviral sgRNA, cell culture media was refreshed to contain puromycin as above for antibiotic selection, and GDF15 or vehicle control was supplemented at the following concentrations: 0.01 pg/mL, 0.1 pg/mL, 1pg/mL, 10pg/mL, 100pg/mL. Thereafter cell culture media and recombinant GDF15 was refreshed every 24 hours. Cell viability was measured 7 days after lentiviral infection using the CellTiter-Glo reagent (Promega, Madison, WI).

### Statistical analyses for experimental studies

All data are expressed as means ± standard deviation. All experimental assays were performed in duplicate or triplicate. Statistical analysis was performed by a two-tailed Student’s t-test, one-way or two-way analysis of variance (ANOVA), Kolmogorov-Smirnov test, log-rank P value, or other tests as indicated. A p value <0.05 was considered statistically significant.

### Data availability statement

Processed data for CRISPR screens (in Figure 3 and Figure 4d) are available in Supplementary Tables 20, 25, and 29. Raw data will be made available upon request. Mass spectrometry data relating to Figure 1 are available in Supplementary Table 12. Raw MS spectra will be made available upon request. L1000 data relating to Figure 2 and Supplementary Figures 3 & 4 is available through the NIH LINCS program and at https://clue.io/data. Data will be made publicly available upon publication. The website lincsproject.org provides information about the LINCS consortium, including data standards.

### Code availability statement

L1000 data analysis code and pre-processed data are available via GitHub https://github.com/cmap/cmapM. There is additional information about this database and tools at http://clue.io/connectopedia. Additional in-house code generated for the ORFeome dataset will be made available upon request.

## Acknowledgements

We thank Daniel Bondeson, Peter Tsvetkov, Steven Corsello, Uri Ben-David and Tamara Ouspenskaia for helpful discussions and critical reading of the manuscript. We thank Mengdan Zhong for technical assistance. We thank David Nusinow and Steven Gygi for insights in identifying small peptides in proteomics datasets. We thank Ross Tomaino for assistance with mass spectrometry at the Talpin Biological Mass Spectrometry Facility at the Harvard Medical School. We thank Jennifer Chen for assistance with the Slncky algorithm. We thank Joshua Gould for assistance with gene datasets. We thank Iain Cheeseman for providing doxycycline-inducible HeLa Cas9 cells. J.R.P. was supported by the Harvard K-12 in Central Nervous System tumors (5K12 CA 90354-18). V.L and M.W.K. were supported by the National Institutes of Health (R01 HD073104 and RO1 HD091846 to M.W.K.). Correspondence and requests for materials should be addressed to T.R.G at

## Author Contributions

J.R.P. and T.R.G. conceived the project, designed experimental approaches, supervised the study, and analyzed data. J.R.P. selected ORFs for screening and developed ORF prioritization methods. J.R.P. and X.Y. designed and generated the ORF cDNA library. J.R.P performed ORF library screening, *in vitro* CRISPR experiments, western blots, cell culture assays, and all GREP1 functional experiments. B.F. executed arrayed ORF screen for L1000. O.M.E. and N.J.L. performed gene expression profiling and analyzed L1000 gene expression data. Z.J. contributed ORF predictions and assisted in analysis of ORF candidates. V.L., A.K., M.K. and J.R.P. performed protein evolutionary analyses and analyzed phylostratigraphy data. K.K., K.R.C., and J.D.J. performed proteomic identification of ORFs from datasets. J.R.P., F.P., and D.E.R. designed and analyzed CRISPR screens. T.G., D.A., and A.B. assisted with sgRNA design. A.G. and Z.K. performed cell line CRISPR screens. L.W., K.S., G.B. and J.A.R. performed pooled CRISPR screening. V.M.W. and J.M.D. analyzed pooled CRISPR screen data. J.M.D. performed comparative analyses of ORF CRISPR data with publicly available CRISPR screens. J.R.P. and T.R.G. wrote the manuscript draft and all authors contributed to editing the manuscript draft.

## Supplementary Figures

**Supplementary Figure 1:**
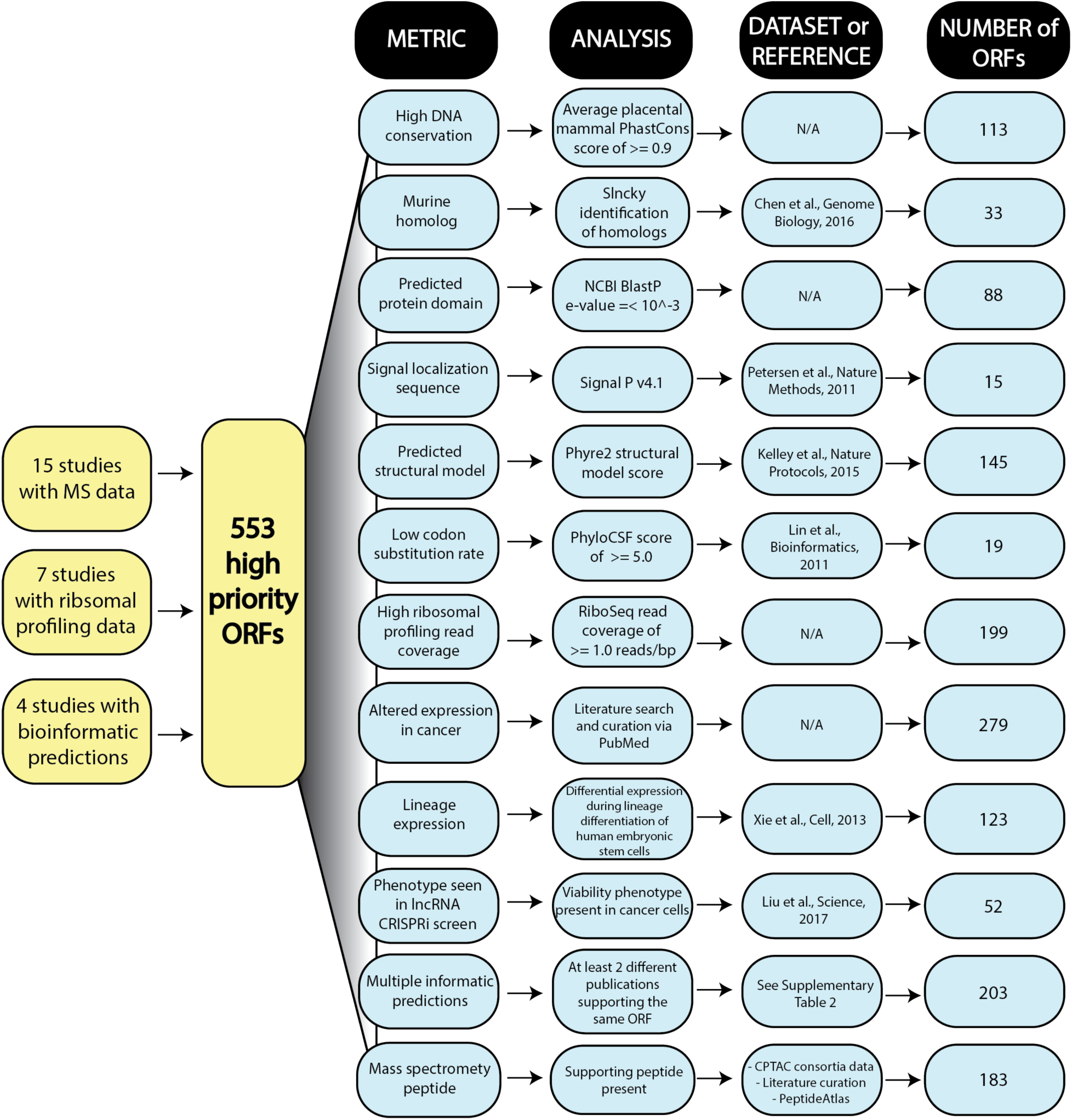
A flowchart of ORF support. Manual curation of ∼9900 ORF loci from the indicated dataset sources were then filtered using the indicated biological attributes and selection criteria. After selection, the 553 ORFs were then evaluated by additional metrics as shown. Please see the Methods for additional details on selection criteria.

**Supplementary Figure 2:**
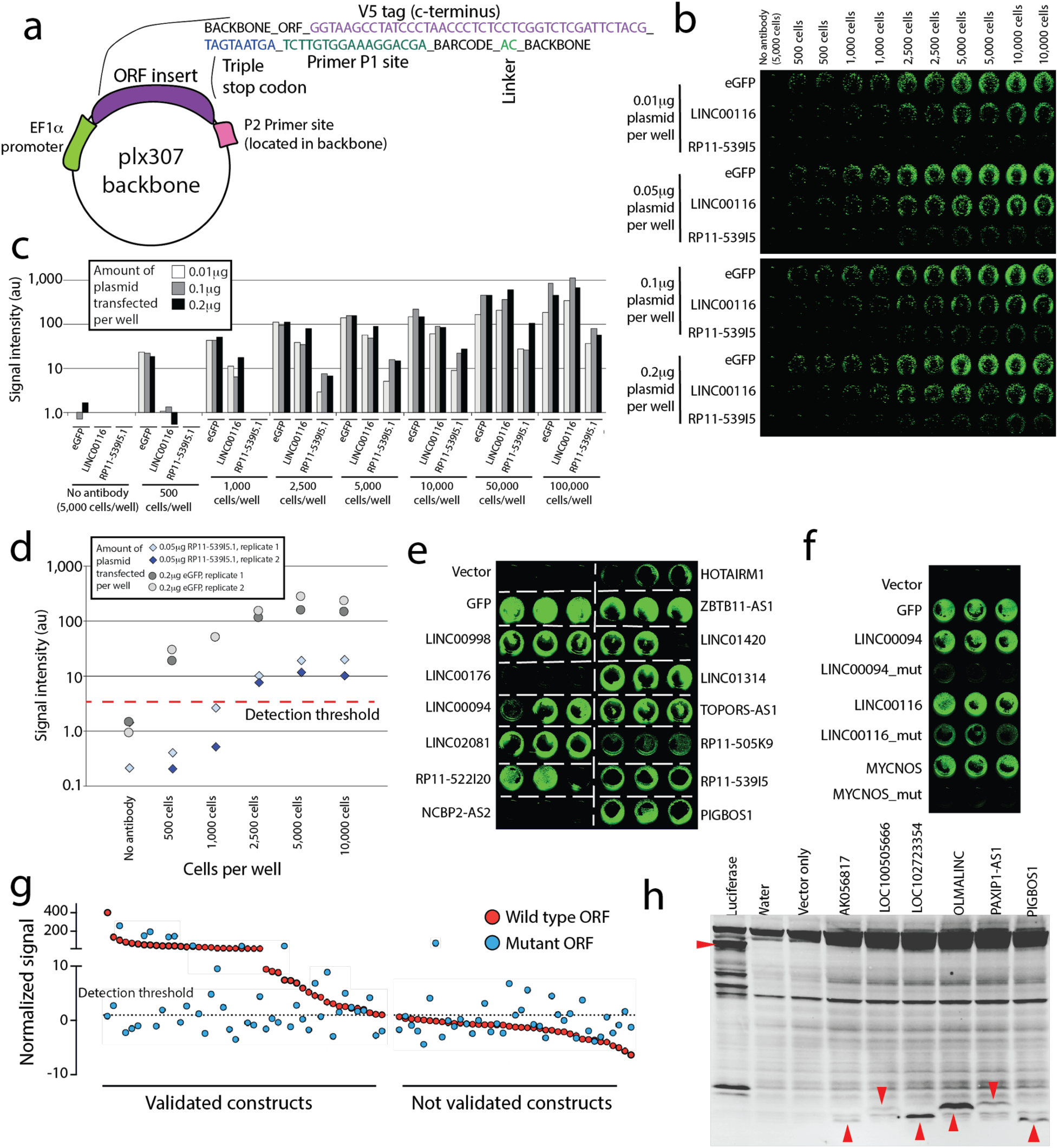
Validation of ORF proteins. **a)** Vector design and sequence details for the ORF library. The vector used is a modified version of the plx307 vector developed by the Genomic Perturbation Platform at the Broad Institute. **b)** Titration analyses of in cell western experiments. Three ORFs were chosen: eGFP (positive control), LINC00116 (high-expressing ORF), and RP11-539I5 (low expressing ORF). Increasing amounts of plasmid were transfected into increasing numbers of HEK293T cells as shown. **c)** Quantification the in cell western titration shown in **b**, demonstrating signal detection over noise and signal plateau. Signal was quantified using pixel density in the 800nM green color channel. **d)** Replicate experiments assessing signal-to-noise thresholds for a low-expressing ORF transfected into HEK293T cells with a low DNA plasmid concentration, as well as a high-expressing ORF (eGFP) transfected into HEK293T cells at a high DNA plasmid concentration. **e)** Example in cell western data in triplicate experiments for selected ORFs. **f)** Abrogation of protein translation via mutation of the ORF for selected examples. **g)** A systematic evaluation of in cell western signal for wild type and mutant ORFs for all pairs. ORFs are separated into those with signal above the baseline threshold, and those without reproducible signal. **h)** An immunoblot showing *in vitro* transcription/translation of selected tag-free ORFs using a wheat germ lysate system. Red arrows indicate the translated ORFs.

**Supplementary Figure 3:**
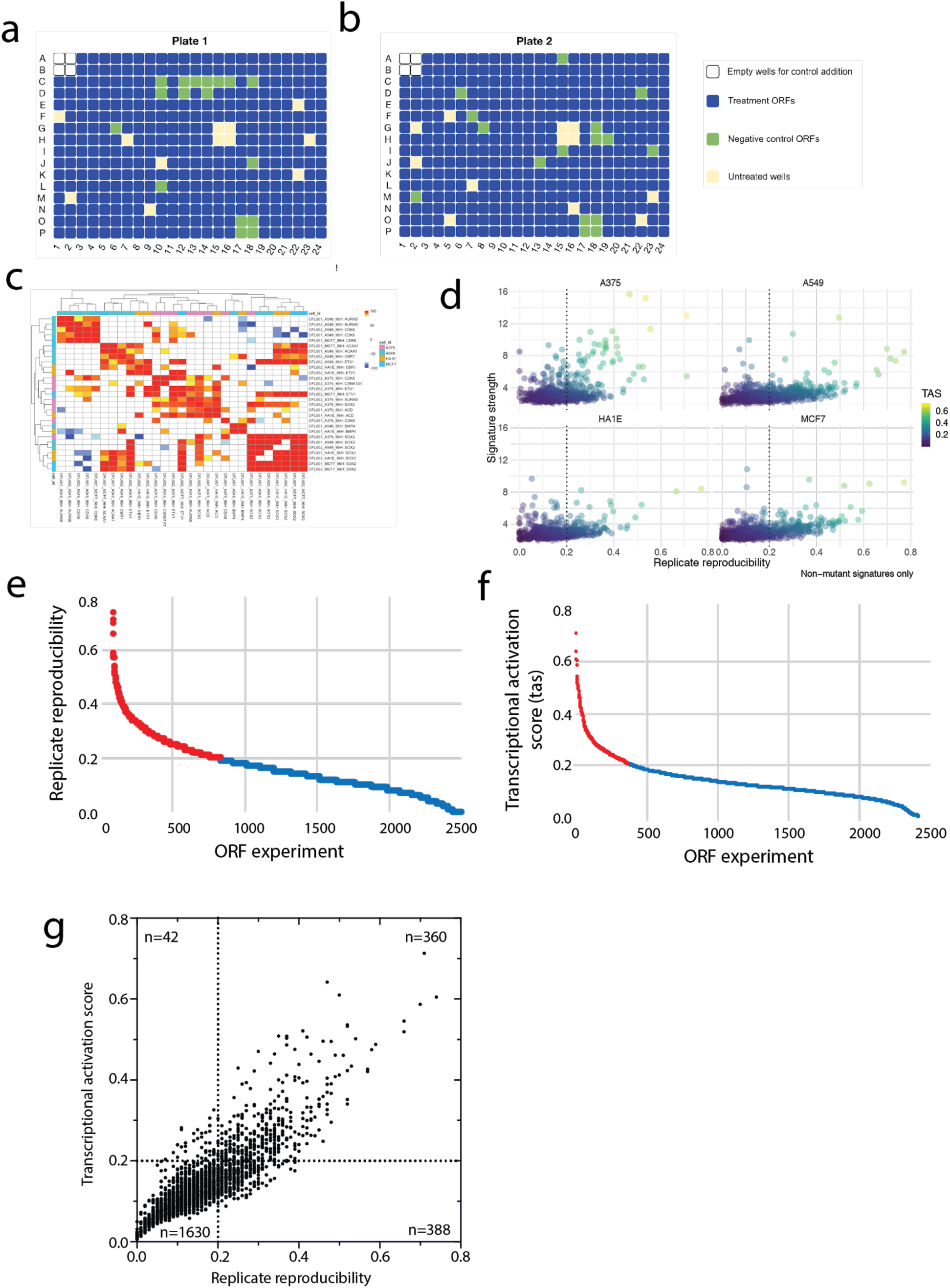
ORF gene expression data on the L1000 platform. **a)** A L1000 perturbational plate layout showing locations of treatment ORFs, non-human proteins, untreated wells, and technical positive control ORFs. **b)** A second L1000 perturbational plate layout showing locations of treatment ORFs, non-human proteins, untreated wells, and technical positive control ORFs. **c)** Level 5 L1000 data processing (“MODZ” score) and clustering of L1000 signatures for positive control ORFs with a TAS score of >= 0.2. Color red in cells denotes a connectivity score of 95 percentile or greater (similar signatures); blue denotes <= −95 percentile (dissimilar signatures). **d)** Scatter plots of L1000 data for experimental ORFs. The Y axis represents signature strength and the X axis represents reproducibility, the two metrics used to calculate the TAS score. Each TAS score is indicated by the color code of each individual ORF. Each data point represents one ORF. **e)** The distribution of replicate reproducibility scores across all L1000 experiments. Red denotes signatures >= 0.2, which indicated that a signature was present. Blue denotes signatures < 0.2, which denotes that a signature was not detected. **f)** The distribution of transcriptional activation scores (TAS) across all L1000 experiments. Red denotes signatures >= 0.2, which indicated that a signature was present. Blue denotes signatures < 0.2, which denotes that a signature was not detected. **g)** Intersection of replicate reproducibility and TAS scores shows a high degree of correlation. 360 signatures were considered positive for both replicate reproducibility and high TAS score.

**Supplementary Figure 4:**
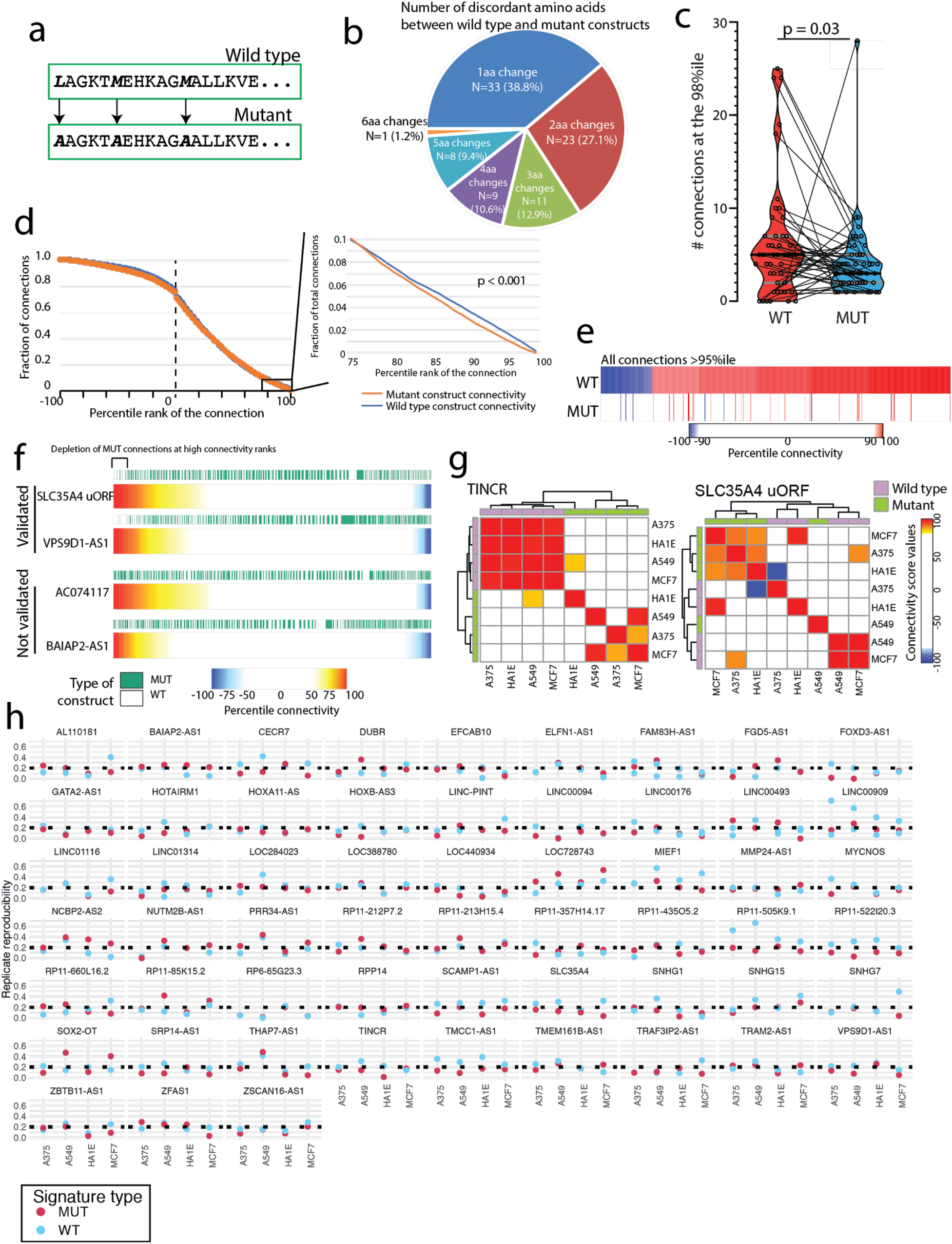
Analysis of paired wild-type and mutant constructs in L1000 data. **a)** An example of ORF mutagenesis strategy in which the start codon and downstream methionines were mutated to alanine. The shown amino acid sequence is fictional and does not represent an ORF in this study. **b)** A pie chart showing the number and percentage of amino acids changed per ORF from the mutagenesis. **c)** A violin plot showing the number of PCL connections made at the 98th percentile for matched mutant and wild type constructs. P value by a Wilcoxon matched pairs rank test. **d)** *Left*, the overall distribution of PCL connections across all ranks in wild type and mutant constructs. *Right*, an inset image of distribution of PCL connections at high connectivity, showing a bias in connections made with wild type constructs. P value by a Wilcoxon matched pairs rank test. **e)** All PCL connections in wild type constructs at either the >=95th percentile or <= -95th percentile, with the matched percentile connectivity in the mutant constructs. **f)** The distribution of percentile connectivity results in wild type or mutant constructs for the indicated genes. In brief, all ORF L1000 signatures were queried against all PCL classes and a percentile connectivity was generated for each individual cell line and for both wild type and mutant constructs. Cell line and construct data was then aggregated and ranked from highest to lowest connectivity. The rank positions of wild type and mutant ORFs were then plotted to reveal a depletion of mutant constructs at high connectivity scores. **g)** Two example heatmaps for the TINCR and SLC35A4 uORF genes showing clustering of PCL connectivity among wild type constructs that is not shared with mutant constructs. Purple bars denote wild type ORF experiments and green bars denote mutant ORF experiments. **h)** L1000 signature replicate reproducibility for all wild type and mutant pairs across all cell lines. All ORF signatures with at least one reproducible wild type signature are shown.

**Supplementary Figure 5:**
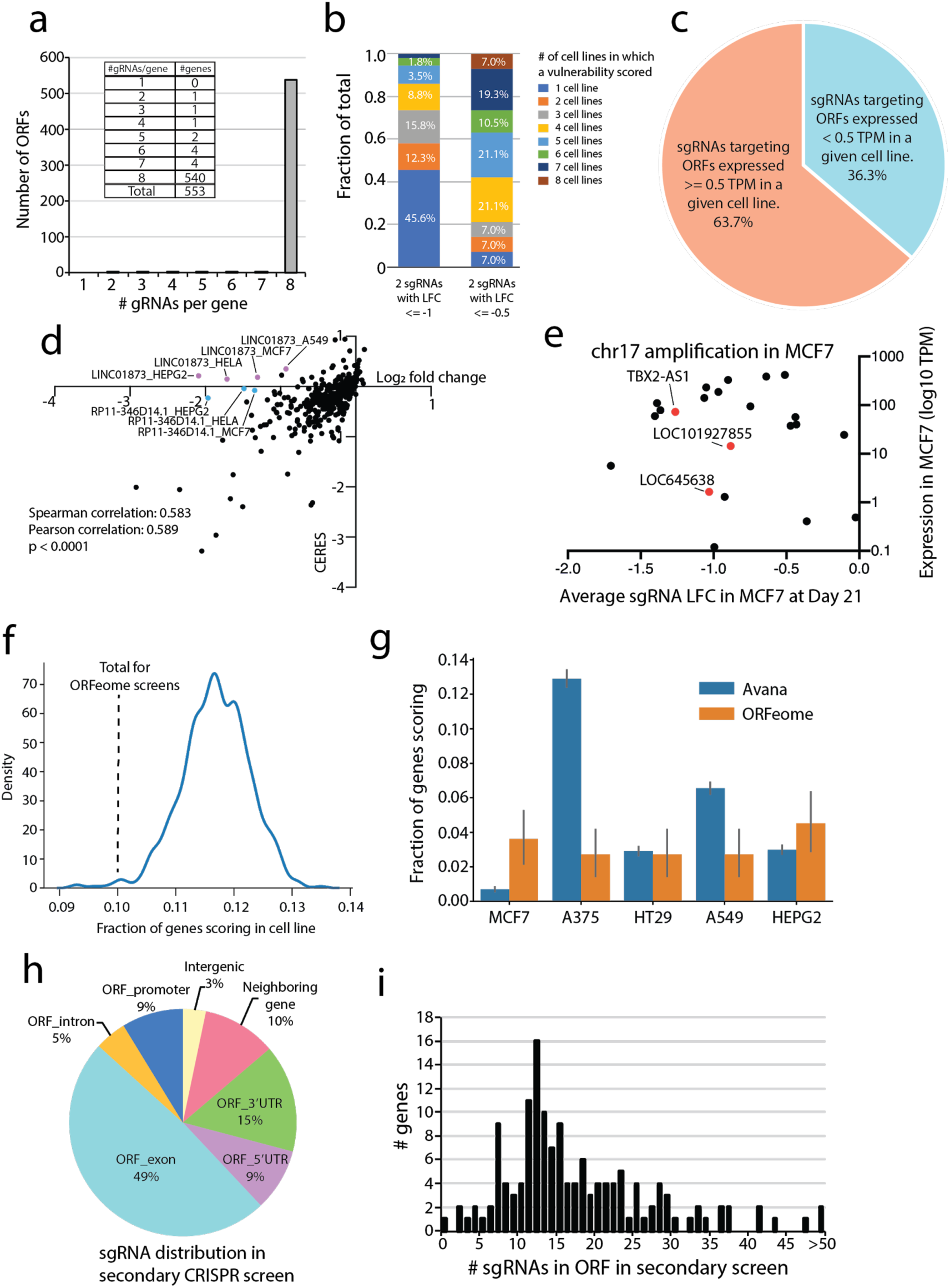
CRISPR screens for new ORFs. **a)** A barplot and inset table showing the number of sgRNAs per ORF in the primary CRISPR screen. **b)** Frequency distribution of putative CRISPR hits using a viability threshold of log fold change of <= −1 or <= -0.5 in the primary CRISPR screen. **c)** The percentage of nominated CRISPR hits which had minimal detectable expression or expressed above the threshold of >= 0.5 TPM. **d)** The correlation between log fold change values for nominated CRISPR hits and the CERES score for each gene, which integrates copy number data for each cell line. Spearman and Pearson correlations are shown with Spearman’s p value shown. **e)** An example of the chr17q23 amplification locus in MCF7 cells. CRISPR knockout of genes in the locus result in nonspecific cell death due to excessive genomic cutting, regardless of gene expression level. Three putative ORFs were located in this genomic region, indicated with red dots in the figure. **f)** A histogram showing the fraction of genes that would score as a vulnerability gene from a randomly selected set of 500 annotated genes from cell lines in the Cancer Dependency Map. The ORFeome CRISPR screen result is indicated. **g)** The rate of genes scoring as viability genes in the canonical Avana gene library and the ORFeome sgRNA library for the five cell lines shared between both screens. **h)** The distribution of sgRNAs across various genome regions in the secondary CRISPR screen. **i)** A histogram showing the number of sgRNAs per ORF in the secondary CRISPR screen.

**Supplementary Figure 6:**
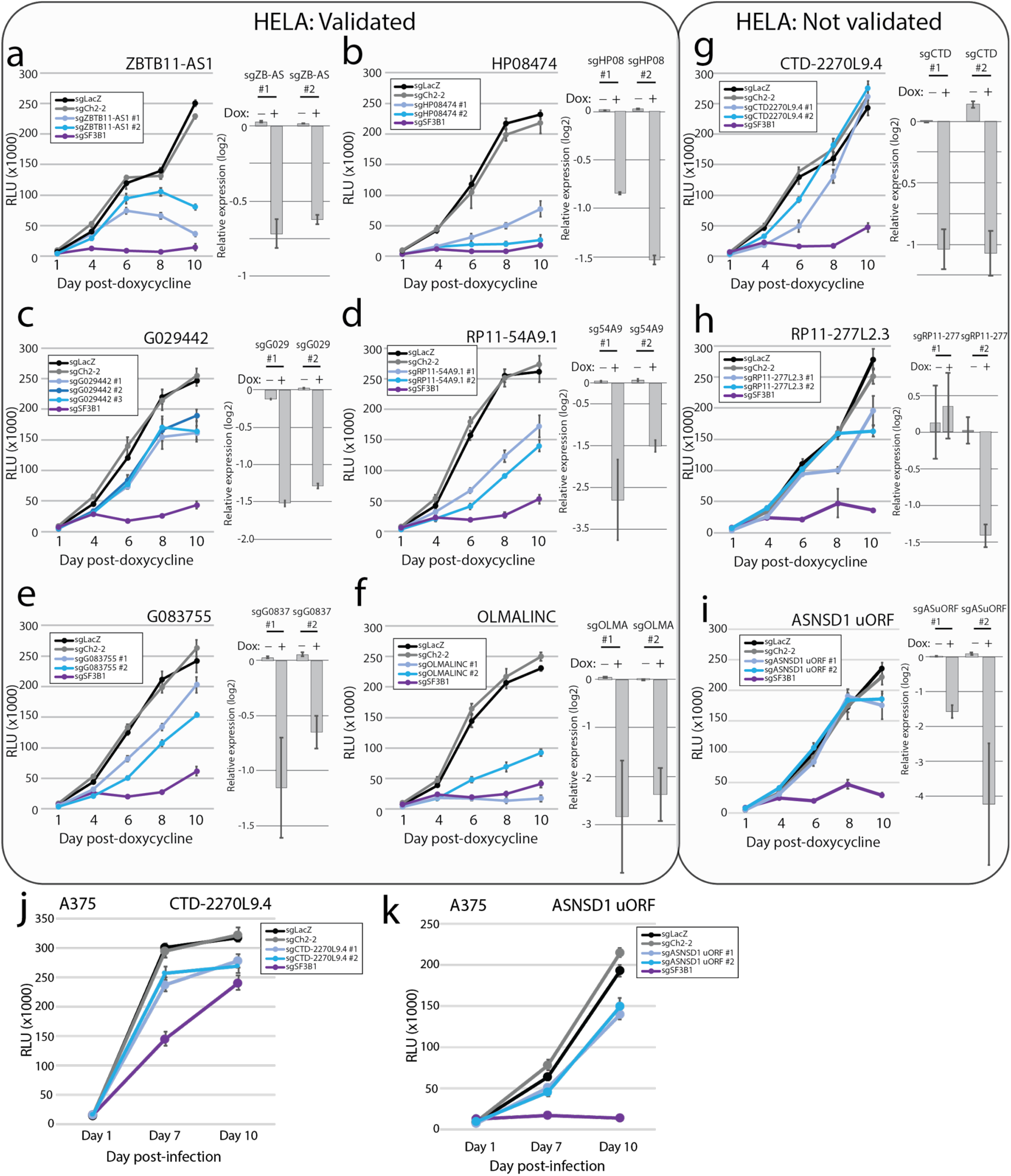
Validation of CRISPR hits via manual assays. **a-i)** CRISPR assays using doxycycline-inducible Cas9 in HeLa cells. Targets are divided in ones that validated and ones that did not. For each experiment, the right-set panel is qPCR data of expression 96 hours after induction of Cas9 with doxycycline. **a)** ZBTB11-AS1 **b)** HP08474 **c)** G029442 **d)** RP11-54A9.1 **e)** G083755 **f)** OLMALINC **g)** CTD-2270L9.4 **h)** RP11-277L2.3, **i)** ASNSD1 uORF. **j-k)** CRISPR assays using stably-expressing A375 Cas9 cells. **j)** CTD-2270L9.4 **k)** ASNSD1 uORF. Error bars represent standard deviation.

**Supplementary Figure 7:**
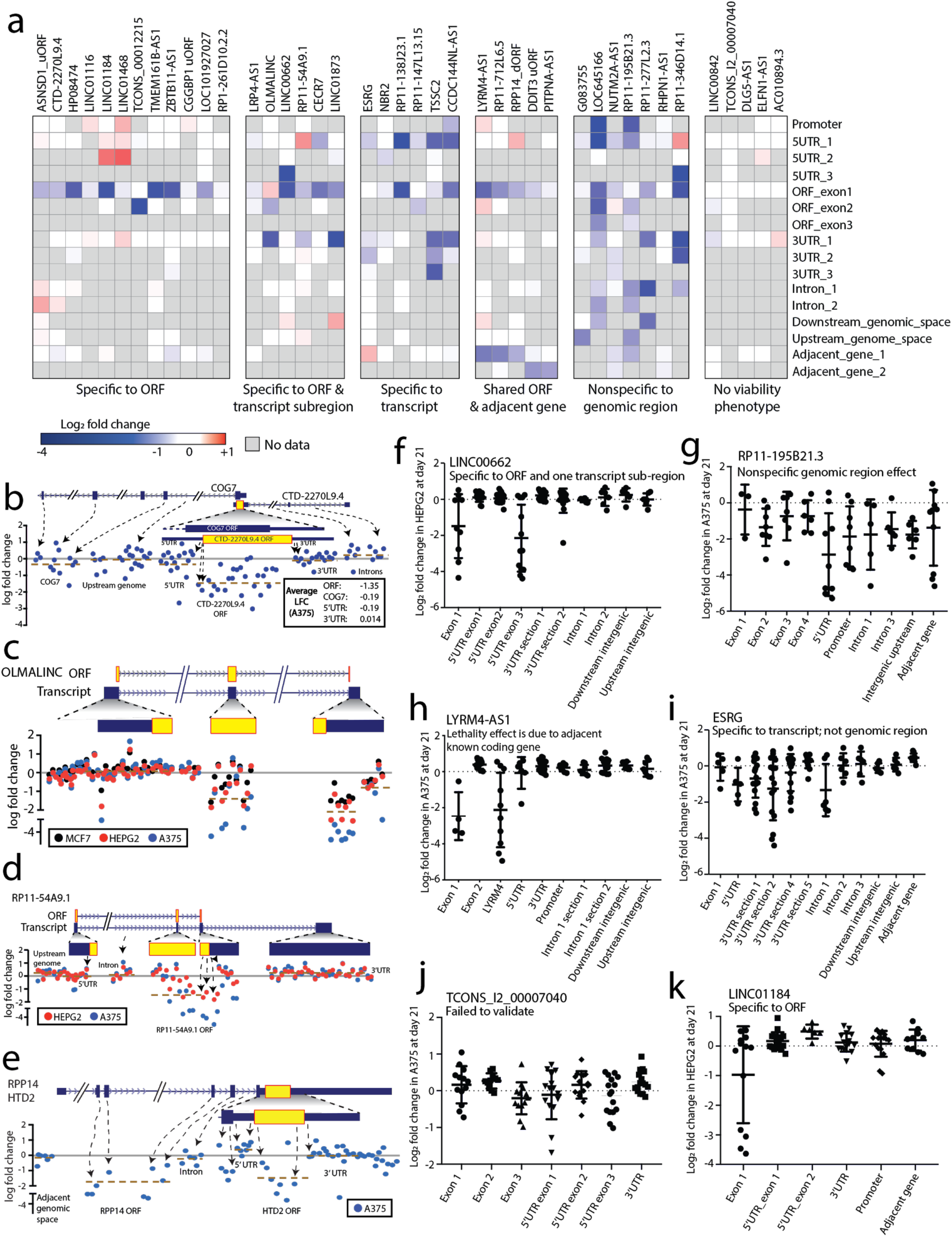
Tiling CRISPR to elucidate functional ORFs. **a)** A heatmap showing log fold change viability loss at Day +21 for the indicated ORFs tested by multiple tiling sgRNA regions. **b-e)** Examples of ORFs with a CRISPR tiling phenotype. **b)** CTD-2270L9.4 **c)** OLMALINC **d)** RP11-54A9.1 **e)** RPP14 dORF / HTD2. **f - k)** Representative sgRNA log fold change data for the indicated transcripts. Each tiling experiment is classified as indicated. **f)** LINC00662 **g)** RP11-195B21.3 **h)** LYRM4-AS1 **i)** ESRG **j)** TCONS_I2_00007040 **k)** LINC01184.

**Supplementary Figure 8:**
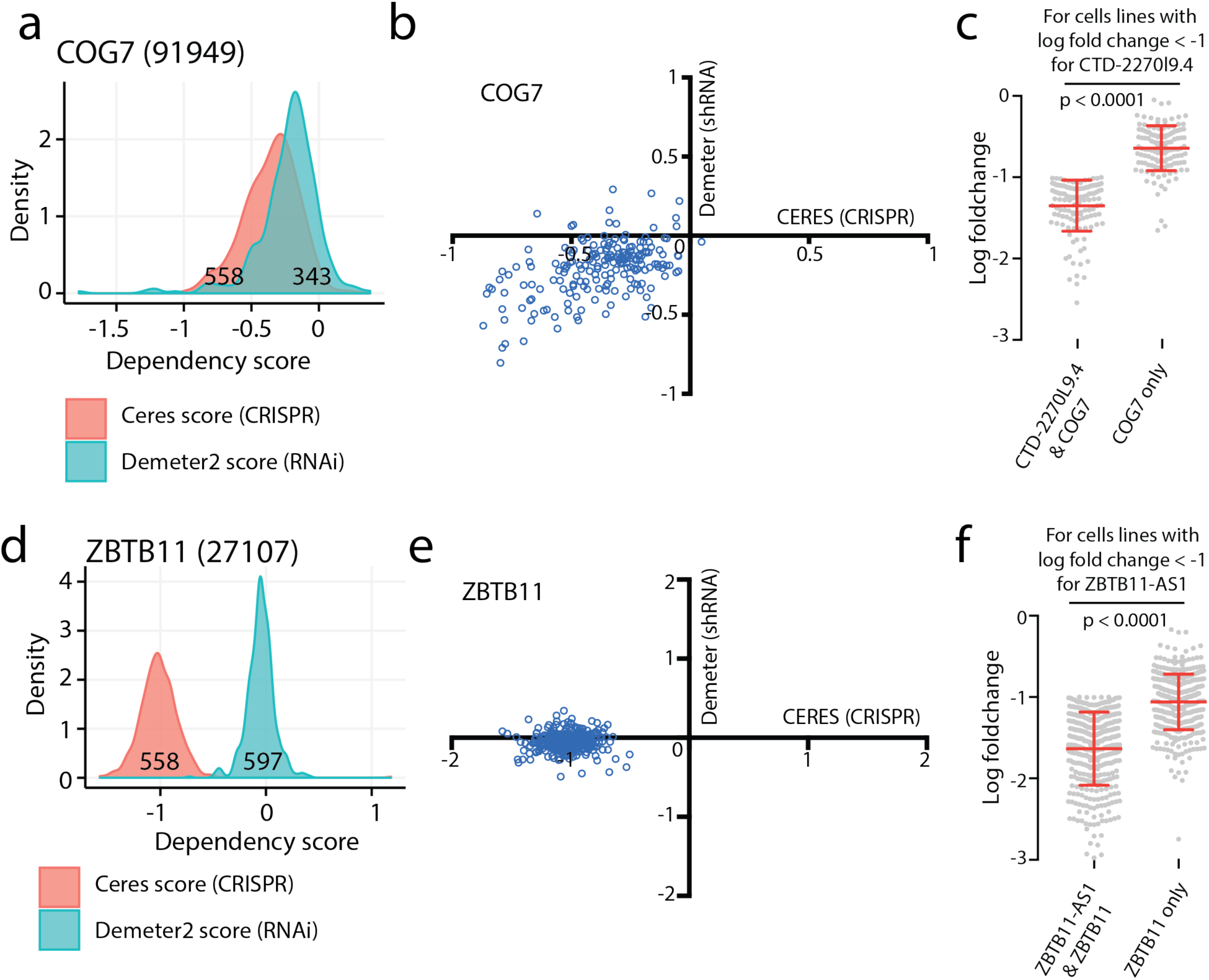
Discordant RNAi and CRISPR data for two overlapping ORFs. **a)** The dependency profile for COG7 using RNAi or CRISPR data. **b)** A scatter plot comparing the magnitude of dependency phenotype for individual cell lines in RNAi or CRISPR data. **c)** A comparison of the log fold change in cell abundance using the average LFC of the two sgRNAs targeting CTD-2270L9.4 and COG7, compared to two sgRNAs targeting COG7 alone. Only cell lines with a viability phenotype in the CTD-2770L9.4 targeting sgRNAs are shown. P value by a two-tailed Student’s t test. **d)** The dependency profile for ZBTB11 in RNAi or CRISPR data. **e)** A scatter plot comparing the magnitude of dependency phenotype for individual cell lines in RNAi or CRISPR data. **f)** A comparison of the log fold change in cell abundance using the average LFC of the two sgRNAs targeting ZBTB11 and ZBTB11-AS1, compared to two sgRNAs targeting ZBTB11 alone. Only cell lines with a viability phenotype in the ZBTB11-AS1 targeting sgRNAs are shown. P value by a two-tailed Student’s t test.

**Supplementary Figure 9:**
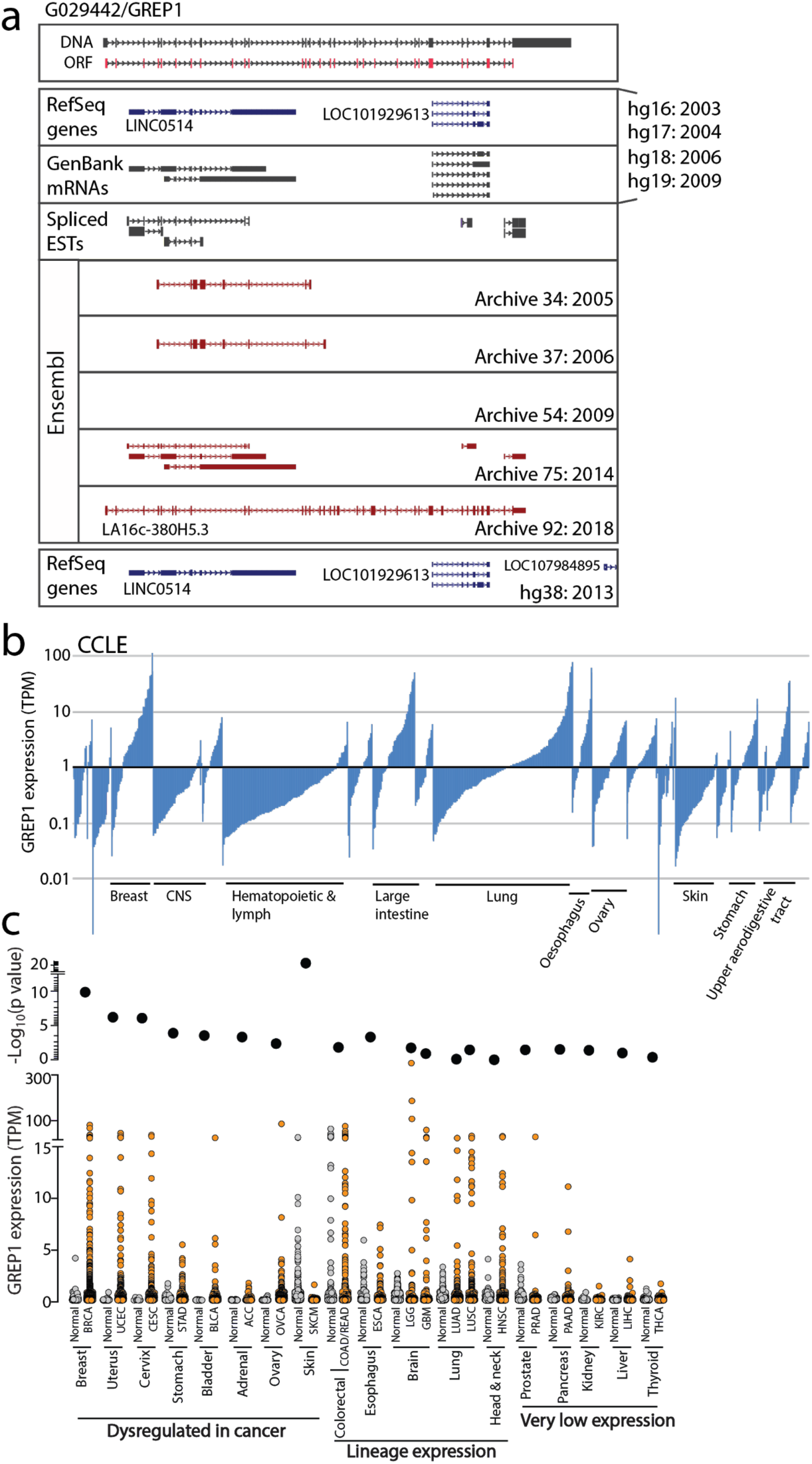
The GREP1 locus and expression. a) A schematic representation of the GREP1 gene structure and the annotation of this locus in the indicated databases. The year of release for each database is indicated. **b)** mRNA expression level of GREP1 across tumor lineages in the Cancer Cell Line Encyclopedia. The Y axis is in a log10 scale. **c)** mRNA expression of GREP1 across tumor types using TCGA and GTex data. A two-tailed Student’s t-test was used to calculate significance of change between normal and cancer tissues. Cell lineages are grouped according to whether GREP1 expression is specifically modulated in cancer, universally expressed as a lineage gene, or not robustly expressed in the indicated lineage.

**Supplementary Figure 10:**
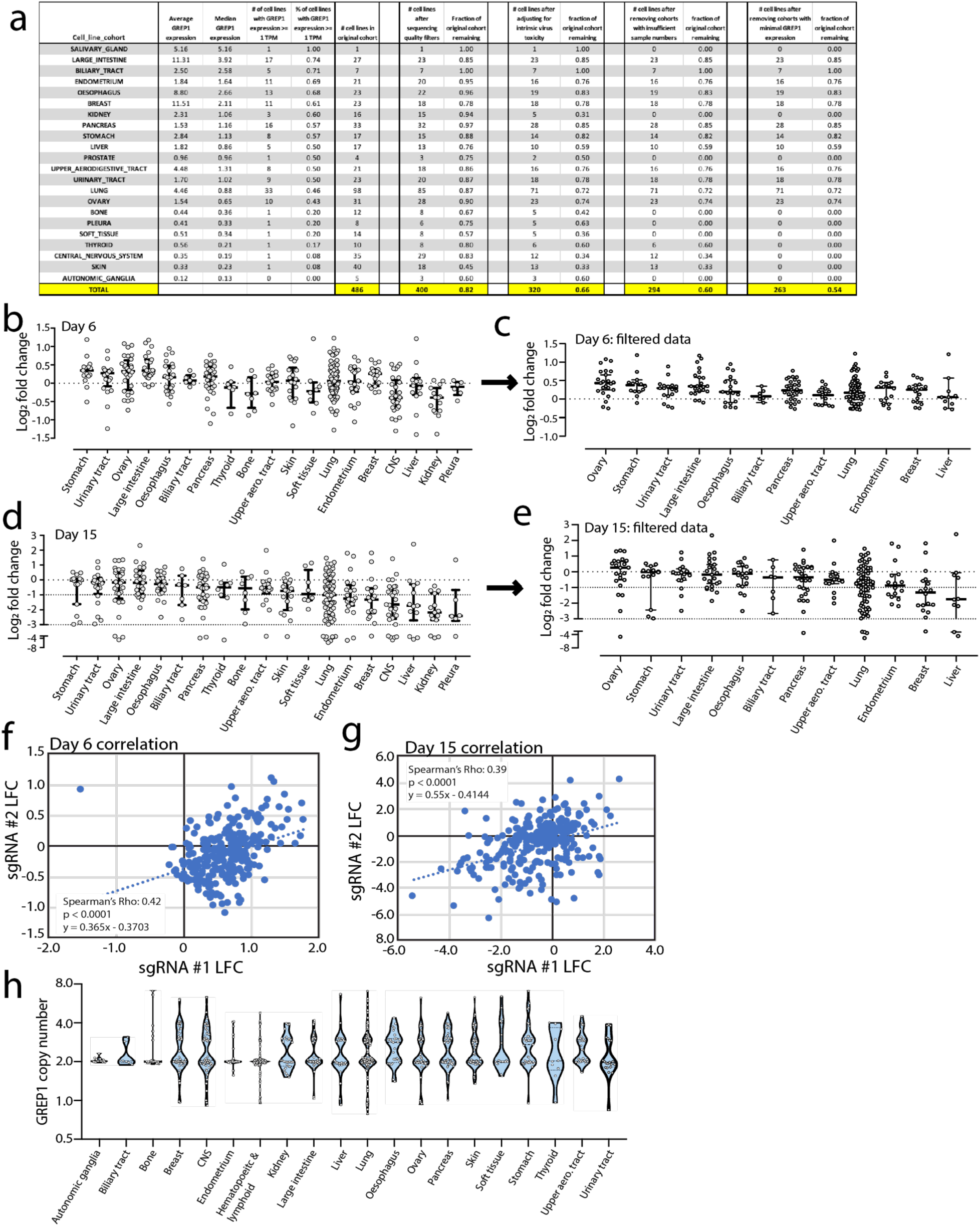
Pooled GREP1 knockout across cell lines. **a)** A table summarizing all input cell lines in the pool and filters applied to the data for final analysis. **b)** All raw cell line viability data at Day +6 prior to data filtering. **c)** Cell line viability data at Day +6 after data filtering. **d)** All raw cell line viability data at Day +15 prior to data filtering. **e)** Cell line viability data at Day +15 after data filtering. **f)** Correlation of G029442 sgRNAs at Day +6 using filtered data. **g)** Spearman’s correlation of G029442 sgRNAs at Day +15 using filtered data. P value for the Spearman’s rho is shown. **h)** G029442 locus copy number profile across cell line tumor types using Cancer Cell Line Encyclopedia data. No cell lineage harbors high-level amplifications.

**Supplementary Figure 11:**
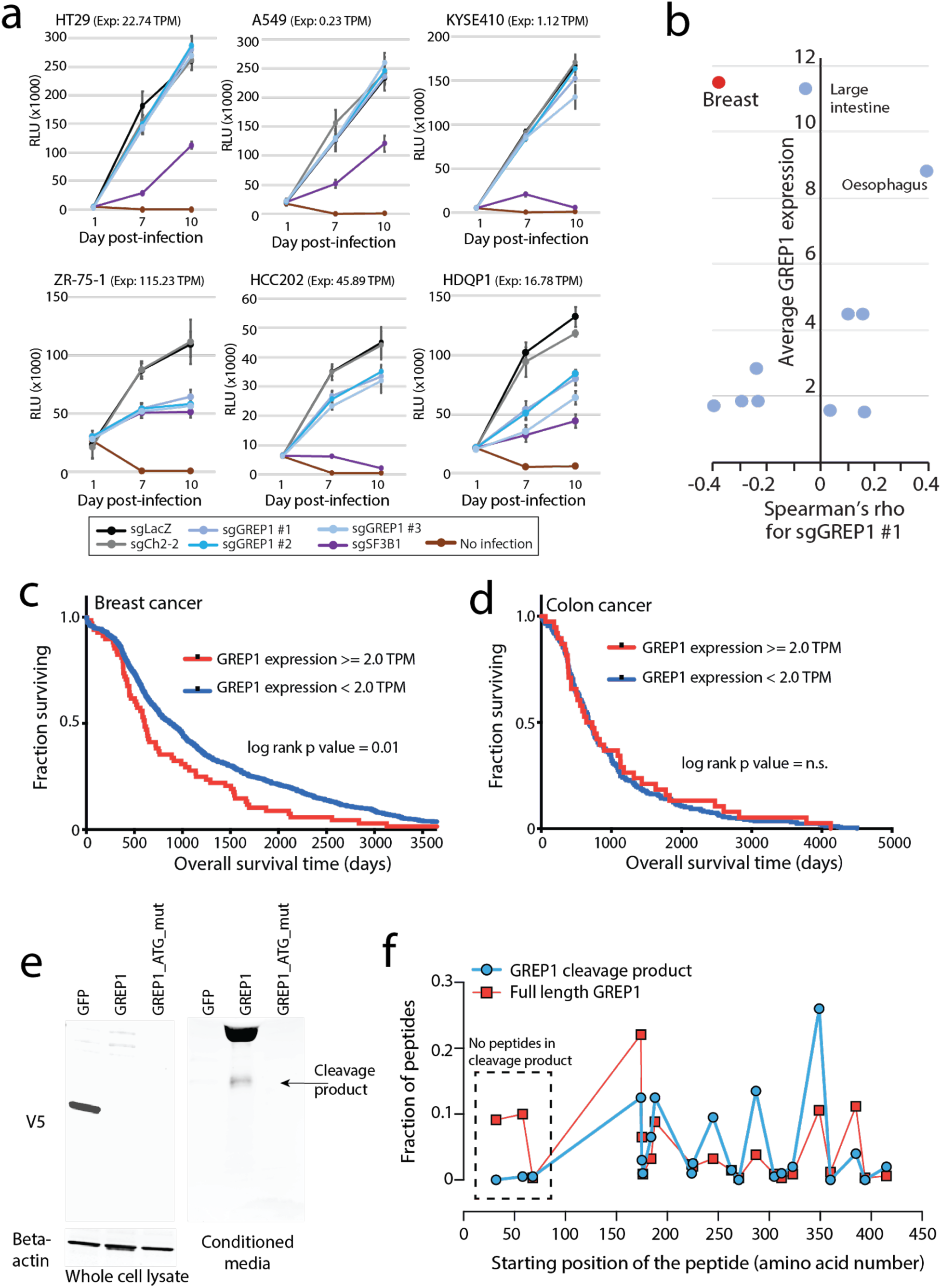
GREP1 is implicated in cell proliferation. **a)** Cell growth curves by CellTiter-Glo following GREP1 knockout in three sensitive and three insensitive cell lines. **b)** A scatter plot showing lineage-specific correlation between cell viability and GREP1 mRNA expression on the X axis with the average GREP1 expression level on the Y axis. **c)** Overall survival for breast cancer patients in the TCGA database stratified by GREP1 expression. Significance by log-rank P value. **d)** Overall survival for colon cancer patients in the TCGA database stratified by GREP1 expression. Significance by log-rank P value. **e)** Immunoblot of V5-tagged GREP1 or GFP in HEK293T cells in both whole cell lysate and conditioned media. A mutant GREP1, in which translational start sites were mutated to alanine, lacks protein translation initiation ability. **i)** Abundance of mass spec peptides detected in the full length GREP1 or cleavage product GREP1 proteins. Peptide abundance is represented as a fraction of total peptides detected. All error bars represent standard deviation.

**Supplementary Figure 12:**
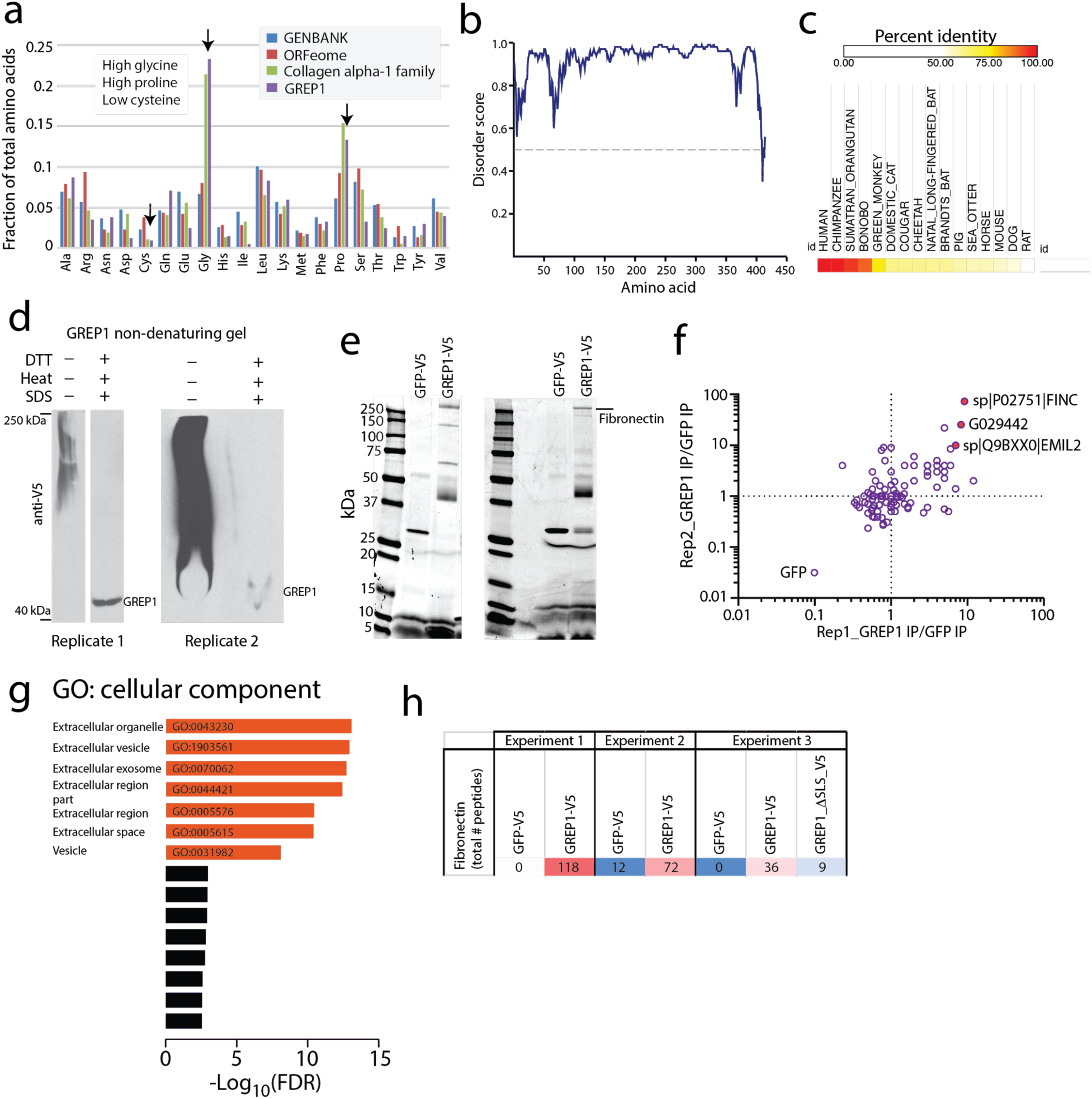
GREP1 is associated with the extracellular matrix. **a)** Total fraction of amino acid usage in the ORFeome, GENBANK, GREP1, and the Collagen alpha-1 family. Sequence similarities between GREP1 and the collagen family are indicated. **b)** Predicted disorder score for the GREP1 amino acid sequence. **c)** Amino acid conservation for detected homologs of GREP1 in the indicated species. **d)** Non-denaturing native western blot of GREP1 in conditioned media from HEK293T cells expressing V5-tagged GREP1. **e)** Representative Commassie-stained gels for immunoprecipitation of GREP1 from the conditioned media of HEK293T cells. Two representative biological replicates are shown. **f)** Enrichment of extracellular matrix proteins in the IP-MS data for GREP1 compared to IP-MS data for GFP. **g)** Gene Ontology Cellular Component analysis of proteins >= 2 fold enriched in GREP1 immunoprecipitation compared to GFP immunoprecipitations. **h)** IP MS total peptide count for fibronectin shown for three separate experiments.

**Supplementary Figure 13:**
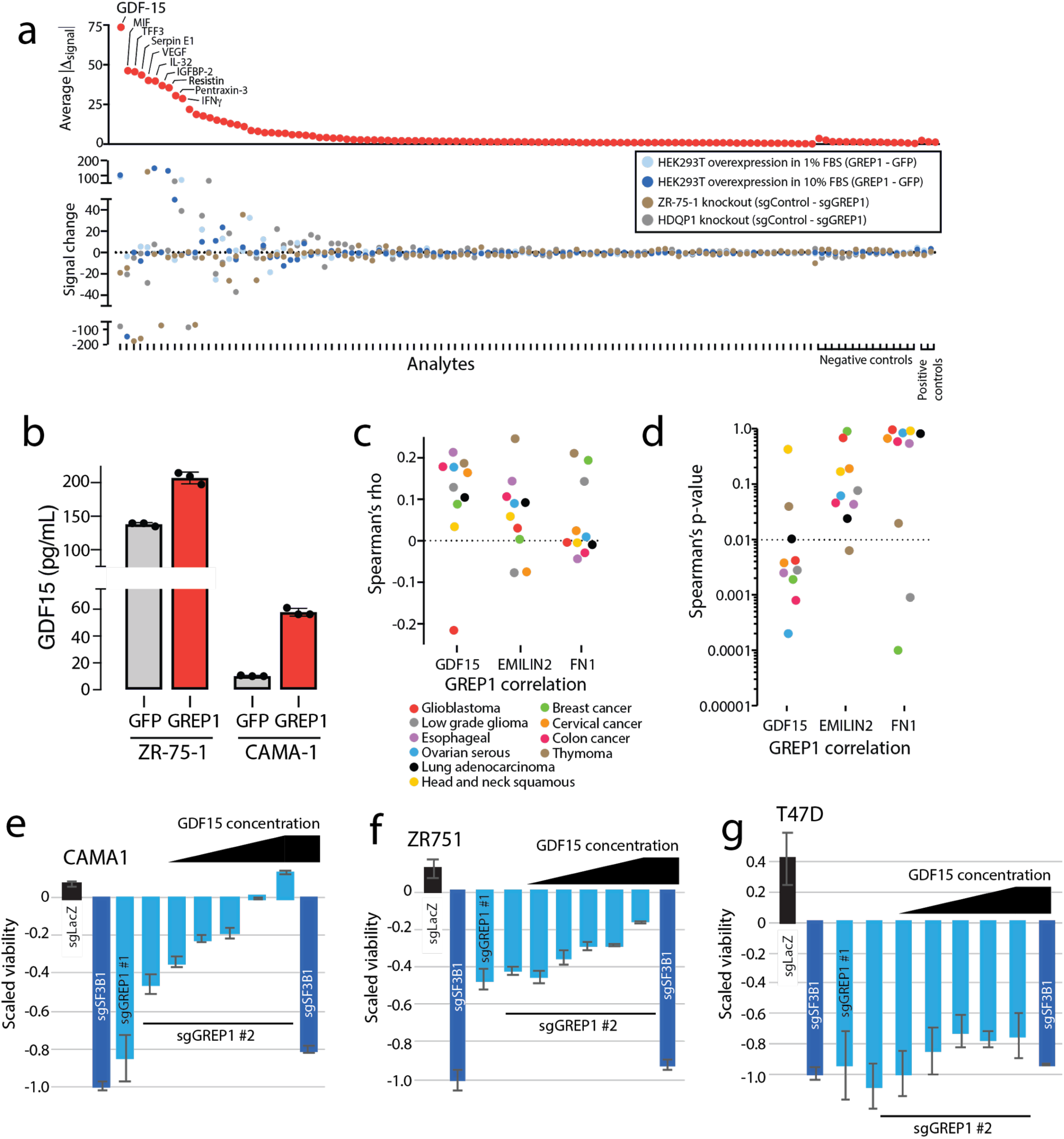
GREP1 regulates GDF15. **a)** Cytokine profiling in HEK293T cells with transient ectopic GREP1 or GFP overexpression, ZR-75-1 cells with stable GREP1 knockout, or HDQP1 cells with stable GREP1 knockout. The change in signal abundance was calculated for each control/GREP1 pair. To rank cytokines, the average of the absolute values for the individual signal changes was plotted. **b)** GDF15 abundance by ELISA in ZR-75-1 and CAMA-1 cells overexpressing a GREP1 or GFP cDNA plasmid. **c)** Spearman’s rho for GREP1 expression correlation with GDF15, EMILIN2, or FN1 in the indicated TCGA datasets. **d)** Spearman’s p value for the GREP1 correlation coefficient for GREP1 correlation with GDF15, EMILIN2, or FN1 in the indicated TCGA datasets. **e-g)** Recombinant GDF15 partially rescues GREP1 knockout. CAMA-1, ZR-75-1 or T47D Cas9 cells were infected with the indicated sgRNAs. 24 hours after infection, cells were treated with vehicle control or increasing concentration of recombinant human GDF15 as shown. Relative abundance was measured 7 days after infection. All error bars represent standard deviation.

**Supplementary Figure 14:**
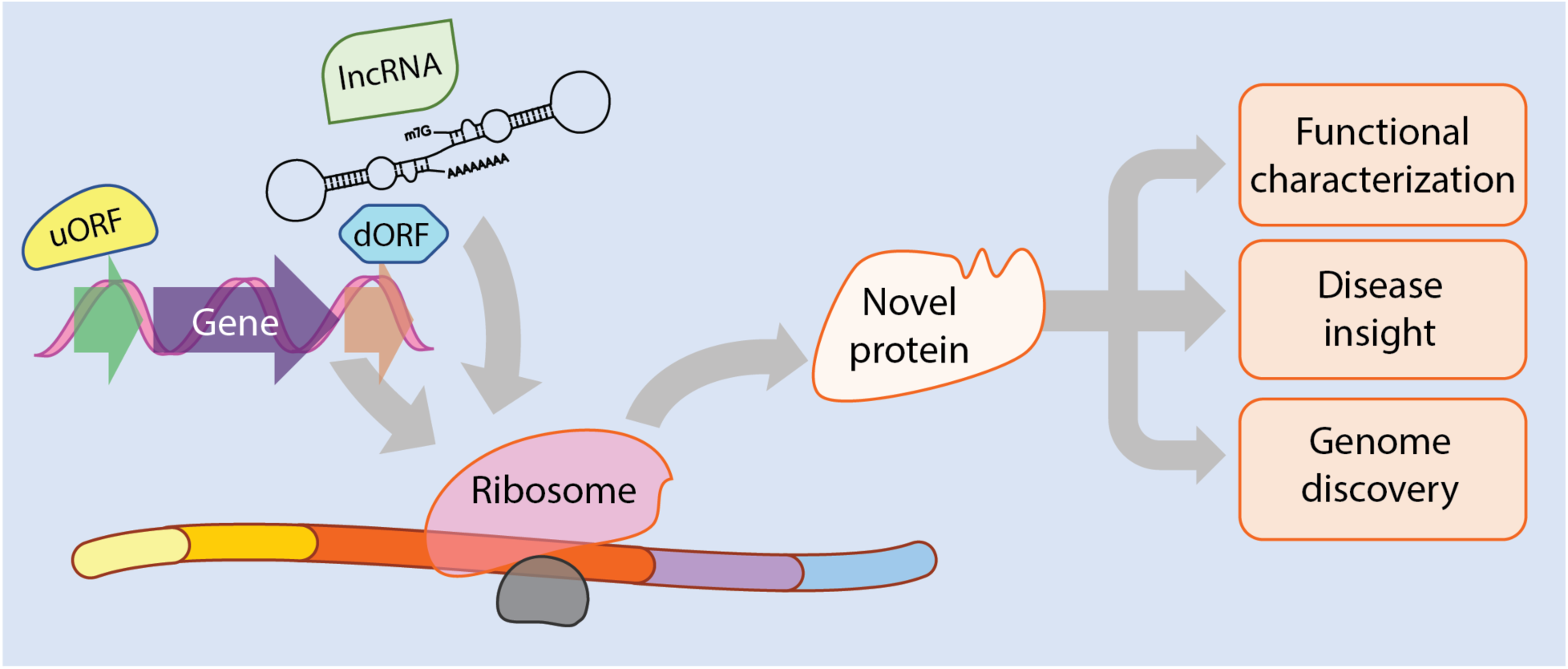
A graphical model.

## List of Supplementary Tables

Supplementary Table 1: Collated datasets for ORF discovery.

Supplementary Table 2: Summarized information about each ORF in the ORFeome library.

Supplementary Table 3: Gene expression for ORFs in the Cancer Cell Line Encyclopedia.

Supplementary Table 4: Distribution of expression in ORFeome compared to annotated mRNAs or annotated ncRNAs.

Supplementary Table 5: Sources of evidence supporting transcriptional dysregulation of uncharacterized ORFs in cancer.

Supplementary Table 6: Putative lncRNAs associated with lineage differentiation of human embryonic stem cells.

Supplementary Table 7: Murine homologs of proteins included in the ORFeome.

Supplementary Table 8: Conversed domains in the ORFeome.

Supplementary Table 9: Predicted structural models for the ORFeome peptide sequences using Phyre2.

Supplementary Table 10: Prediction of signal localization sequences in the ORFeome.

Supplementary Table 11: ORFeome genes implicated in loss of cell viability by CRISPRi data.

Supplementary Table 12: Peptides from mass spectrometry data.

Supplementary Table 13: Summarized peptide support for ORFeome genes.

Supplementary Table 14: 96 well plate data for ORFeome in cell western blot data.

Supplementary Table 15: Phylostratigraphy analysis of ORFeome.

Supplementary Table 16: L1000 expression profiling data.

Supplementary Table 17: Percentile rank connectivity scores of ORF L1000 signatures to reference Perturbational Classes (PCLs).

Supplementary Table 18: The number of PCL connections for ORF wild type/mutant pairs, stratified by percentile rank level.

Supplementary Table 19: CRISPR guide RNAs used in the primary CRISPR screen.

Supplementary Table 20: Primary screen CRISPR data for 8 cell lines at day 7 and 21 post-infection.

Supplementary Table 21: Log fold change data for each sgRNA in the primary screen CRISPR data for 8 cell lines at day 21 post-infection.

Supplementary Table 22: CRISPR guide RNAs used in the secondary CRISPR screen.

Supplementary Table 23: Distribution of sgRNAs in the secondary CRISPR screen.

Supplementary Table 24: Number of sgRNAs taking each coding ORF exon and total ORF region in the secondary screen.

Supplementary Table 25: Secondary screen CRISPR data at day 7 and 21 post-infection.

Supplementary Table 26: Summarized concordance results from primary and secondary CRISPR screens.

Supplementary Table 27: Individual sgRNA sequences used for validation experiments.

Supplementary Table 28: Summarized results from CRISPR tiling experiments for 41 ORFs.

Supplementary Table 29: Pooled CRISPR log fold change data for all cell lines without filtering.

Supplementary Table 30: Pooled CRISPR log fold change data for cell lines after filtering.

Supplementary Table 31: Identification of glycosylation sites on GREP1 through mass spectrometry.

Supplementary Table 32: Abundance of GREP1 peptides in the full length protein and cleavage product.

Supplementary Table 33: Gene Ontology analysis for proteins with >= 2 fold enrichment in GREP1 IP compared to GFP IP in the conditioned media of ectopically-overexpressed HEK293T cells.

## Supplementary Materials

### Supplementary discussion

#### Historical perspectives on the human genome annotation

The human genome is now generally felt to have ∼19,029 protein-coding genes (*Homo sapiens* CCDS release 22 as of October 10, 2019). The single largest gene discovery project was the Human Genome Project (HGP). The RefSeq database included approximately ∼10,000 genes prior to publication of the HGP^1, 2^, which had doubled from the 4,270 genes in the July 1995 GenBank Release 89.9^3^. Many of these genes were known from positional cloning and other techniques.

The initial HGP in 2001 postulated 30,000 - 40,000 human genes. By itself, this was a dramatic reduction in the ∼50,000 - ∼100,000 anticipated genes^4–6^. However, by the revision of the HGP in 2004, this number had been decreased to 20,000 - 25,000^7^. It was subsequently reduced to ∼19,000, with ∼17,600 confidently observed by mass spectrometry^8^. This number has been the current estimate for the past 10 years and the number used as the basis of all exome sequencing studies.

#### Assumptions made during gene discovery

In the HGP, mRNAs were queried for the presence of an open reading frame that was >= 100 amino acids and began with a methionine start codon. If present, this ORF was reported as a novel protein in the HGP. Such methods had basis in precedent, but were not without challenges: the established noncoding RNA Xist was initially reported to have a 894 bp ORF^9^ until it was determined that this ORF was not actually coded^10^.

Proteins less than 100 amino acids were included in the HGP only if they had been previously known, as such ORFs were difficult to predict due to noise from existing cDNA fragments at that time. Therefore, ORFs less than 100 amino acids were not nominated solely based on computational analyses^11–13^.

#### Protein size and function

There is no specific scientific rationale for why smaller proteins would be less real. An analysis by John Mattick and colleagues suggested that an ORF of >100 amino acids was approximately two standard deviations above the average random ORF size in a random 1kb segment of genome sequence^14^. This is statistic, though, is not particularly meaningful as most genes are much longer than 1000bp due to extended untranslated regions (UTRs). However, it highlights the challenge in computationally separating signal from noise.

It is not clear whether there is a minimum size required for peptide/protein function. The smallest known functional unit is a zinc finger, which is an aggregation of the Cys2-His2 four amino acid motif. It is typically thought that a minimum of four or five such motifs are required for functional zinc finger DNA binding, thus suggesting that a peptide of 20 amino acids or greater may be eligible for this function. Secondary structure for a peptide may exist with as few as four or five amino acids^15^, and enkephalins are five amino acid peptides found in the central nervous system and thought to be functional^16^. An alpha helical peptide can be stably produced with 14 amino acids^17^. There are also now known proteins less than 50 amino acids^18, 19^.

#### Skew in the size distribution of annotated proteins

Most annotated proteins are >100 amino acids in most organisms. As shown below, the fraction of the annotated proteome for humans, *C. elegans*, *D. melanogaster* and *D. rerio*.

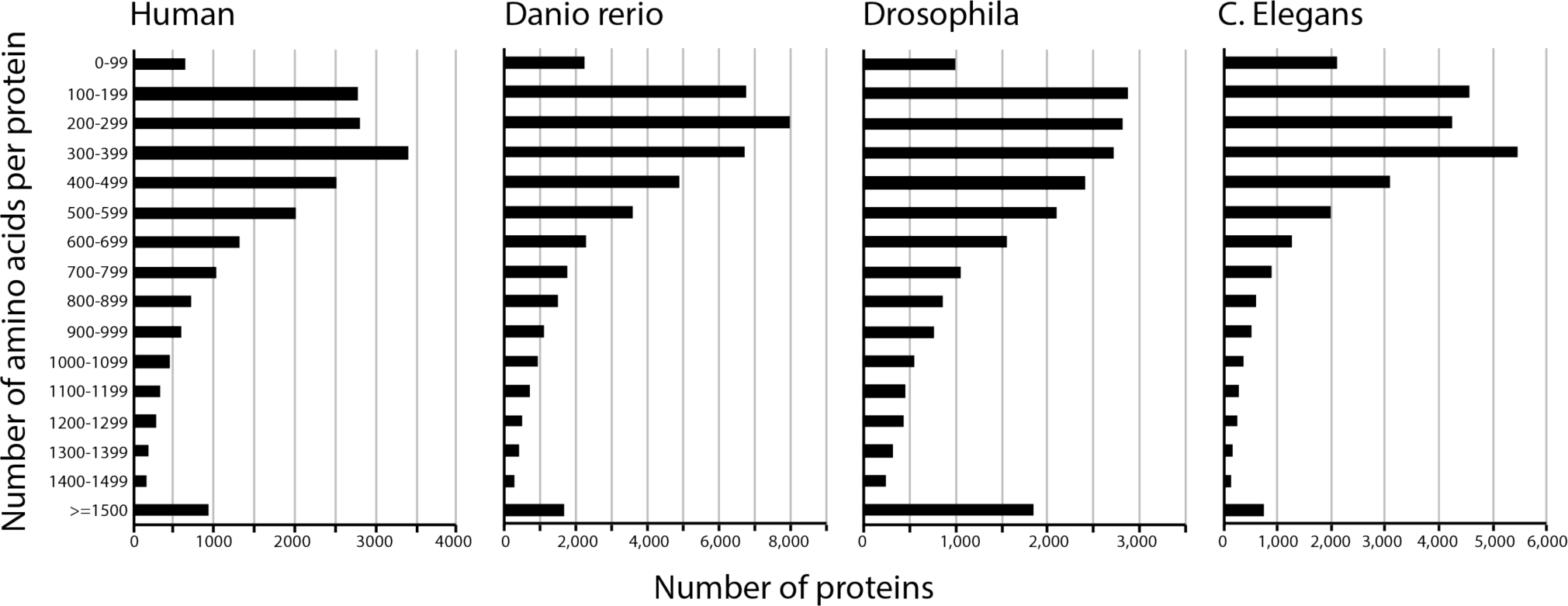

Among human proteins <100 amino acids, 61% are 90 - 99 amino acids large, and thus proteins < 90 amino acids are very rare in annotated databases. Below these data are shown in figure formation for *H. sapiens*.

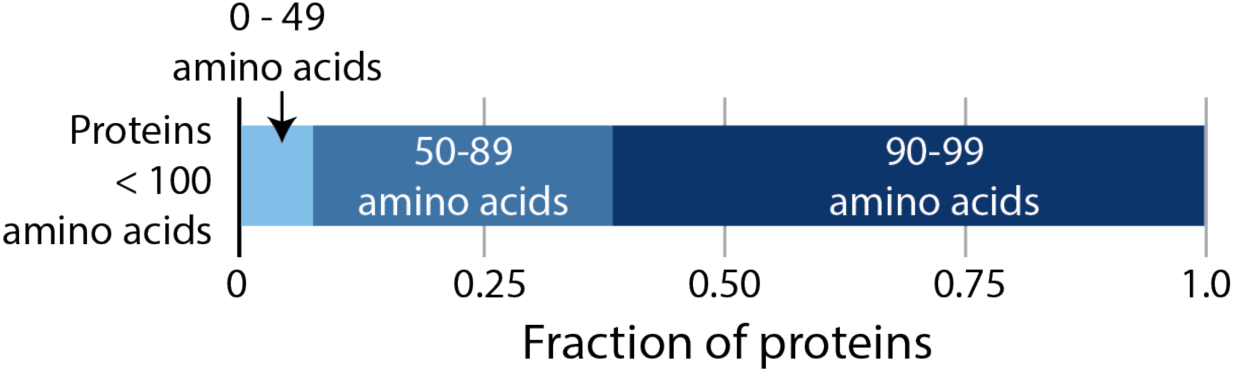

#### Methods to validate a putative protein

Once a potential protein is identified, there are many possible ways to demonstrate its existence. Mass spectrometry of endogenous peptides can provide evidence, though small proteins often have few trypic sites and may not perform well by mass spectrometry. Also, many unannotated proteins are likely lineage-restricted and may not be historically well represented in the mature tissues profiled by mass spectrometry.

Tag-free biochemical transcription/translation with rabbit or wheat germ lysates can be used, but these assays have a high false negative rate and are biochemical assays only. *In vitro* studies can include ribosome profiling/polysome association to see if the mRNA is bound by ribosomes, though this is not direct evidence of translation. Other *in vitro* studies are exogenously expressing an epitope-tagged plasmid construct. However, the epitope tags may destabilize small proteins, leading to protein elimination.

Other approaches include development of a new antibody for a protein for experimental use. This approach is limited as it is expensive and takes a significant amount of time. A genetic knock-in of a fusion-tagged cDNA is also possible, but again costly and time-intensive.

#### Expression of the ORFeome compared to other lncRNAs

It is well-established that, in general, so-called lncRNAs are more tissue-restricted and lower expressed than annotated human proteins^20, 21^. To evaluate the expression level of the ORFs in our ORFeome, we were able to extract gene expression data for 13,049 ncRNA, 18,165 mRNA, and 446 of our ORFs in the Cancer Cell Line Encyclopedia dataset^22^. We found that the ORFs were significantly higher expressed than baseline ncRNAs, though less highly expressed than canonical proteins. See figure below (p values by the Kolmogorov-Smirnov test):

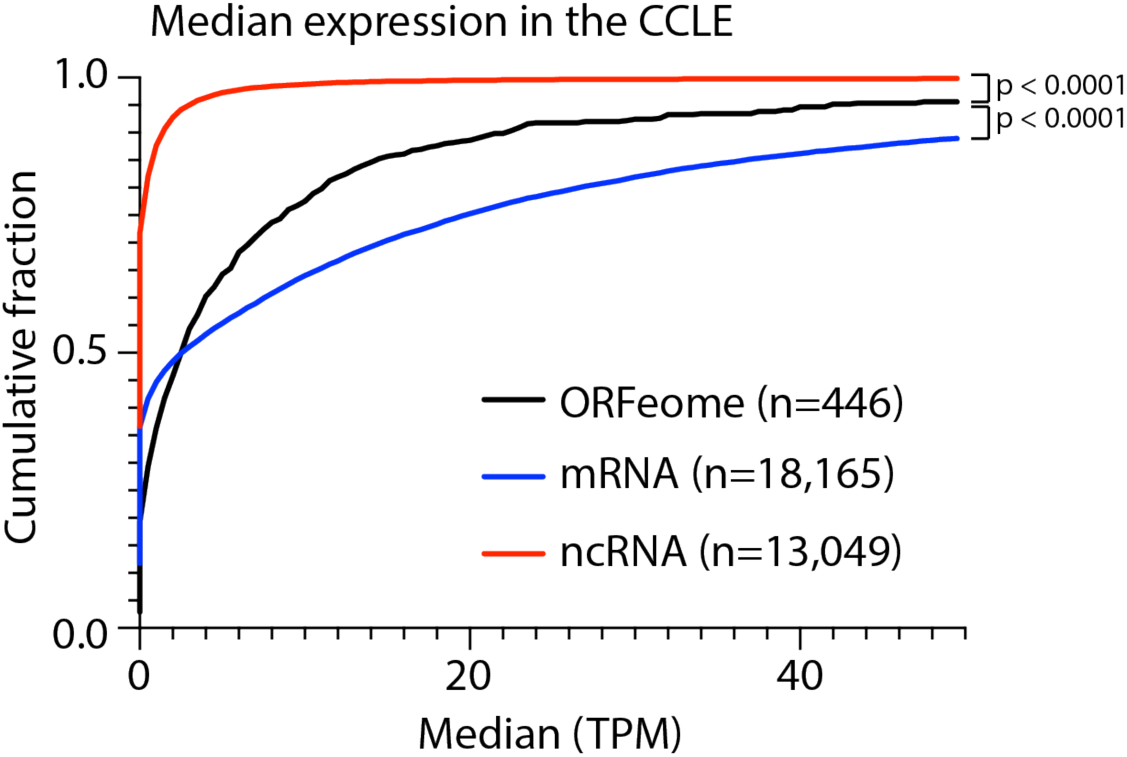

#### Features of the ORFeome amino acid sequences

For the 490 ORFeome ORFs with predicted amino acid sequences longer than 40 amino acids, we evaluated several biophysical properties, including protein sequence length, number of protein binding-sites, aggregation propensity, disorder and number of Pfam-annotated protein domains. First, the amino acid sequences of these ORFs suggest that they have a large proportion of their outer surface exposed to water (73% ± 0.4%), have a high number of predicted protein-binding sites (12.79 ± 0.2 per 100 aa) and disorder (0.98 ± 0.04 per 100 aa), and that have few Pfam-annotated protein domains (0.08 ± 0.01 per 100 aa). In contrast, average mammalian genes, including human genes, encode much longer proteins of ∼500 aa that have a low amount of disorder and high aggregation propensity^23, 24^.

#### Distinguishing a predictive structural model from background signal

Predicting protein structure was performed with the PHYRE2 server^25^ for the 530 ORFs that were >= 40 amino acids in length. To control for the chance of randomly predicted protein structure, we created a score to distinguish background signal. Each amino acid sequence was given a percent confidence score and an alignment coverage percentage by the PHYRE2 server. We multiplied these two numbers together to create a protein structure score. We then computationally generated a list of 500 random 150 amino acid peptides with a methionine start site, and analyzed these in the same manner. We used the distribution of these datasets to define a threshold for determining the presence of a robust structural prediction. See figure below (p value by the Kolmogorov-Smirnov test):

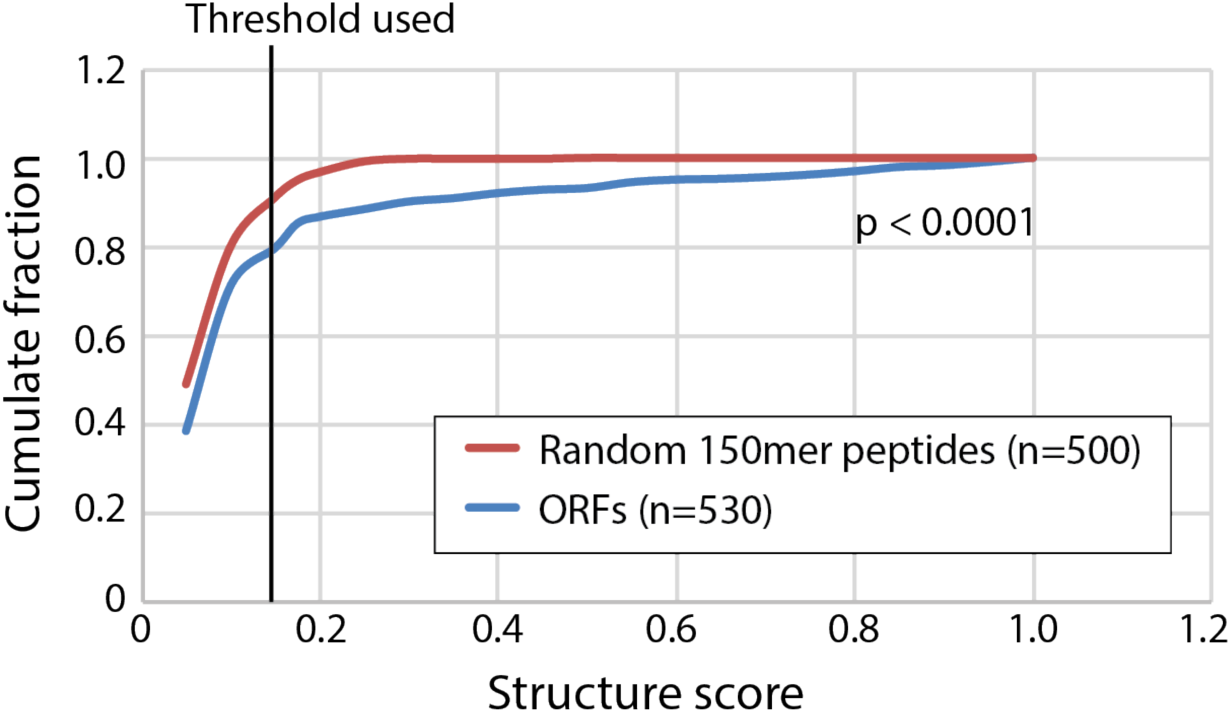

#### Updated annotation status of the ORFs in this manuscript

This project was initiated in January 2016 and therefore we employed databases available at that time. Over the past several years, these gene annotation databases have been updated, but our study was not able to accommodate changes in annotation status due to the nature of large-scale ORF and CRISPR library generation for functional genomics. Therefore, a subset of the genes included in this study are now annotated in the recent versions of GENCODE. A few of the ORFs in this study have now been functionally characterized and published in other studies as well.

We have now re-evaluated the annotation status of our ORFs in GENCODE v31. There are 61 ORFs that are now annotated as protein-coding in GENCODE v31. 43 of these 61 (70.5%) are annotated as the same ORF in GENCODE v31 as in our ORFeome. 2 of the 61 are annotated as different ORFs in the two databases. 44 of the 61 (74%) validated in our V5 western blot assay as a translated protein. The table below shows a list of ORFs that are now annotated as protein-coding, along with the current transcript name and a publication investigating that ORF, if available.

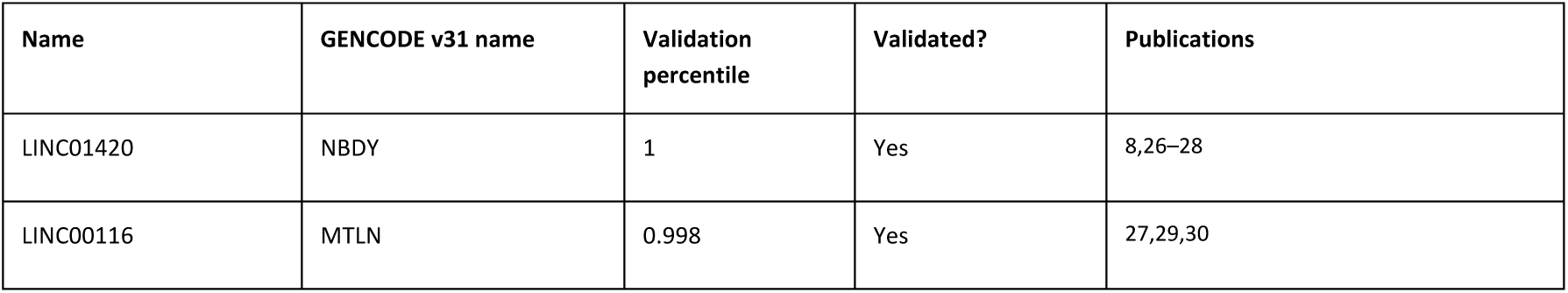
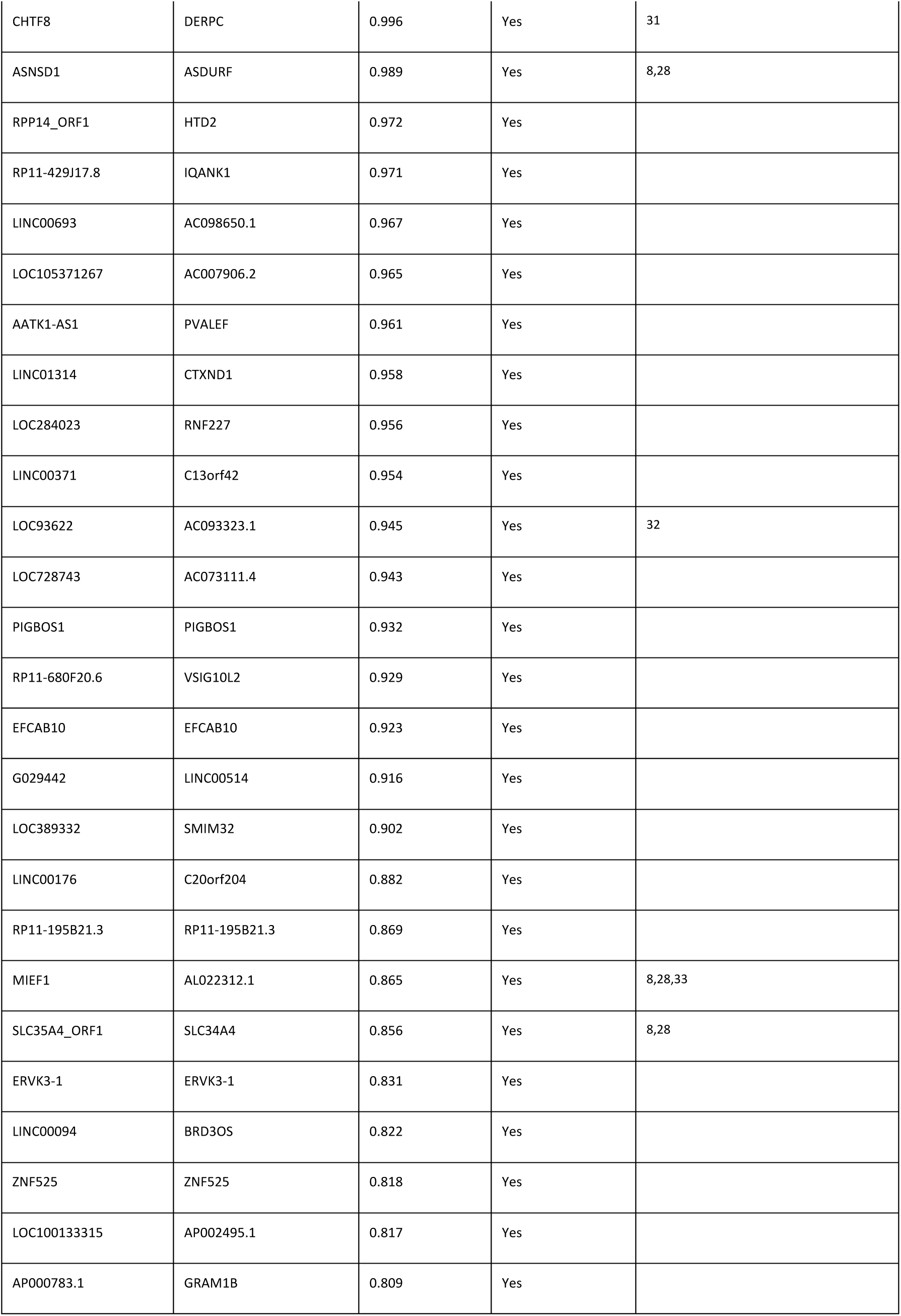
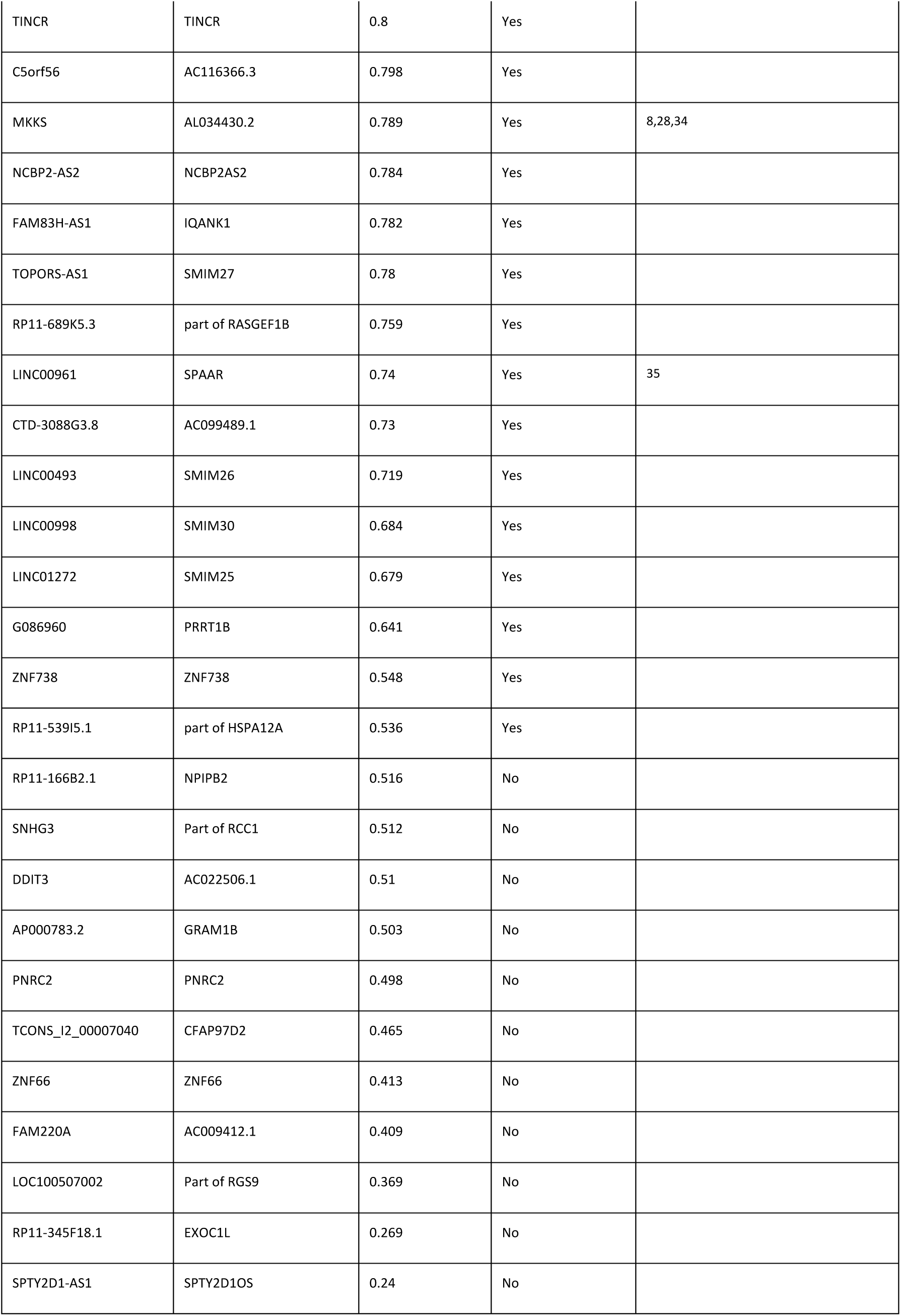
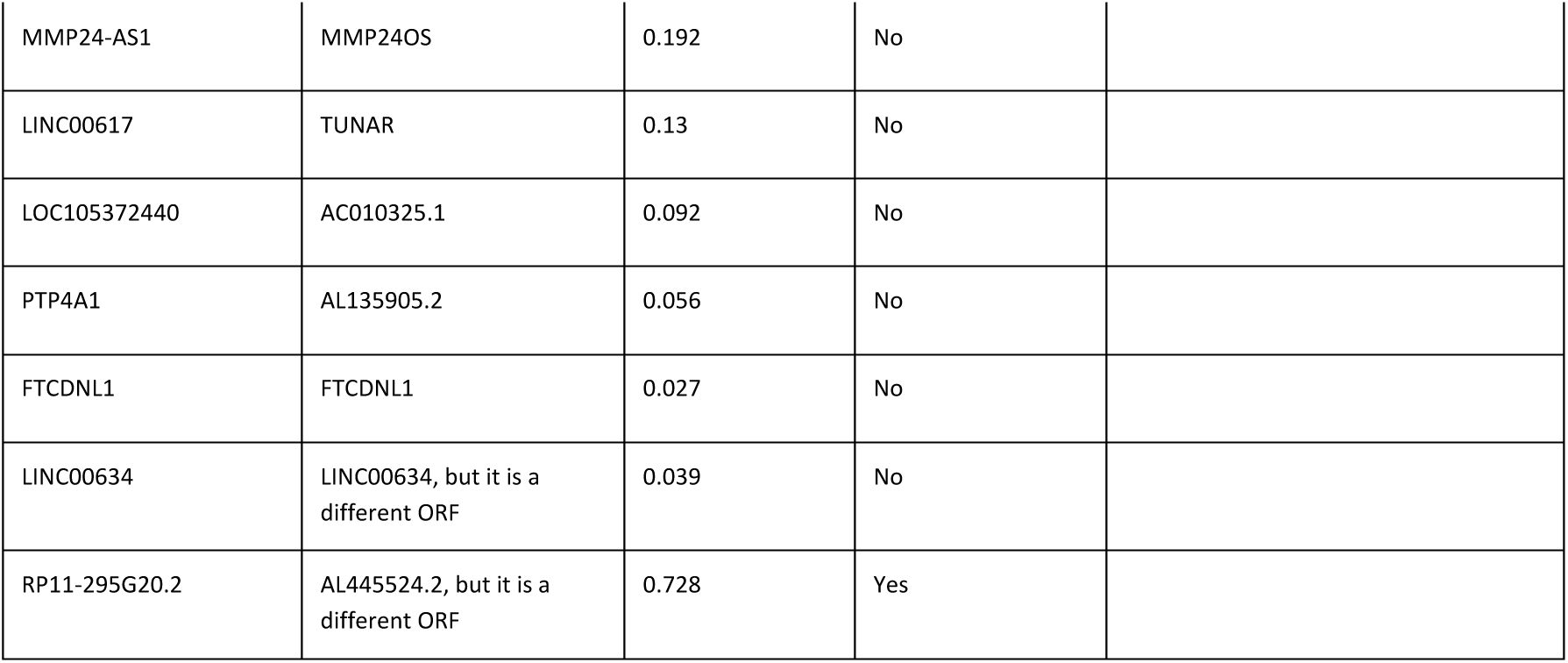

#### Implications of this research

The human genome houses the basis for much of human biology and disease. Understanding the full complement of human protein-coding genes and their functions is a critical component of biomedical science. If a large fraction of human proteins remains undiscovered or unexplored, this provides a tremendous opportunity to expand the purview of research, with further opportunities to search for both the biological causes of disease as well as potential therapeutic targets.

### Immunoblot figures

**Immunoblot Figure 1:**
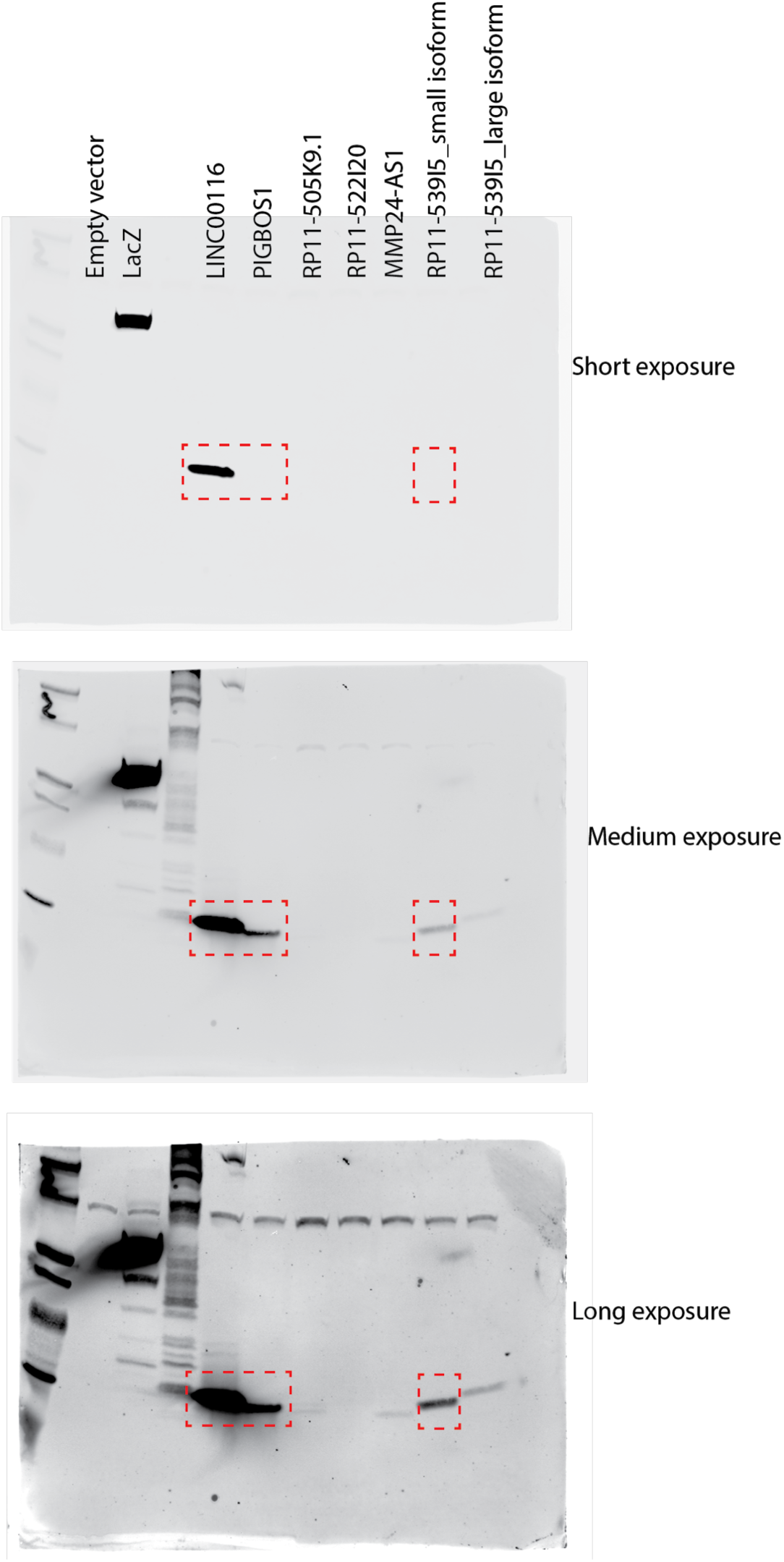
Uncropped images for ORF proteins in Figure 1. The dotted red box indicates the images used in the figure.

**Immunoblot Figure 2:**
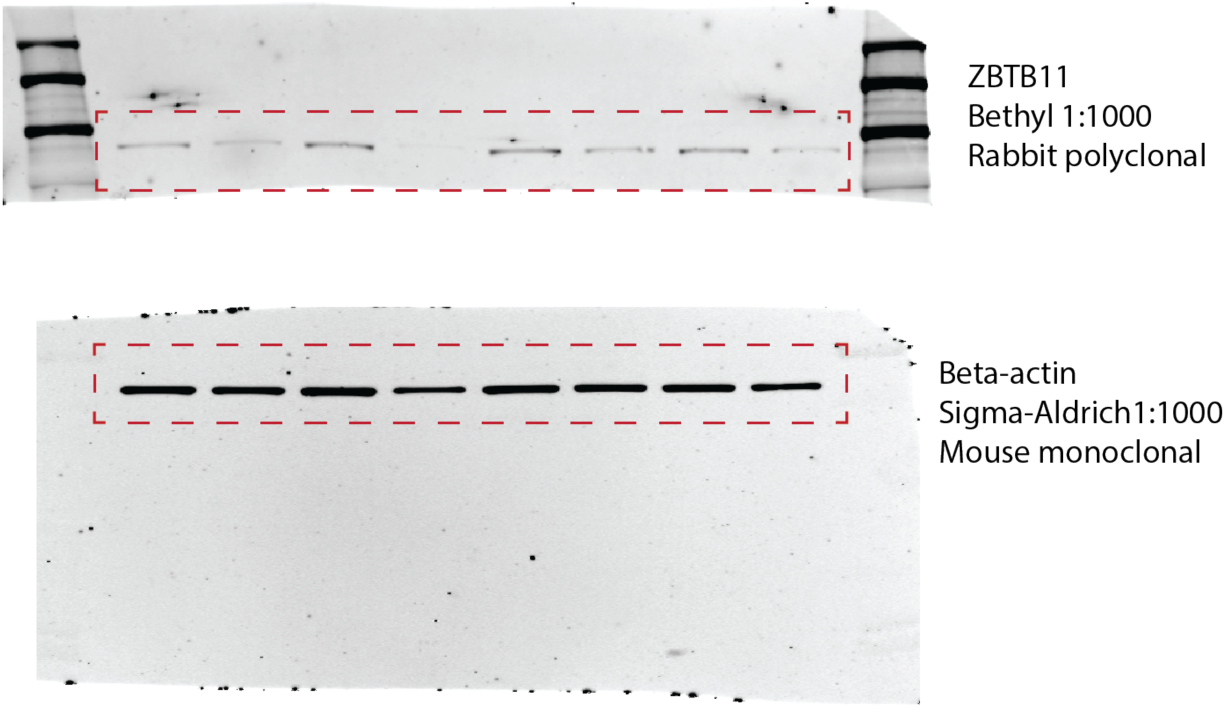
Uncropped images of ZBTB11 and Beta-actin immunoblots. The dotted red box indicates the images used in the figure.

**Immunoblot Figure 3:**
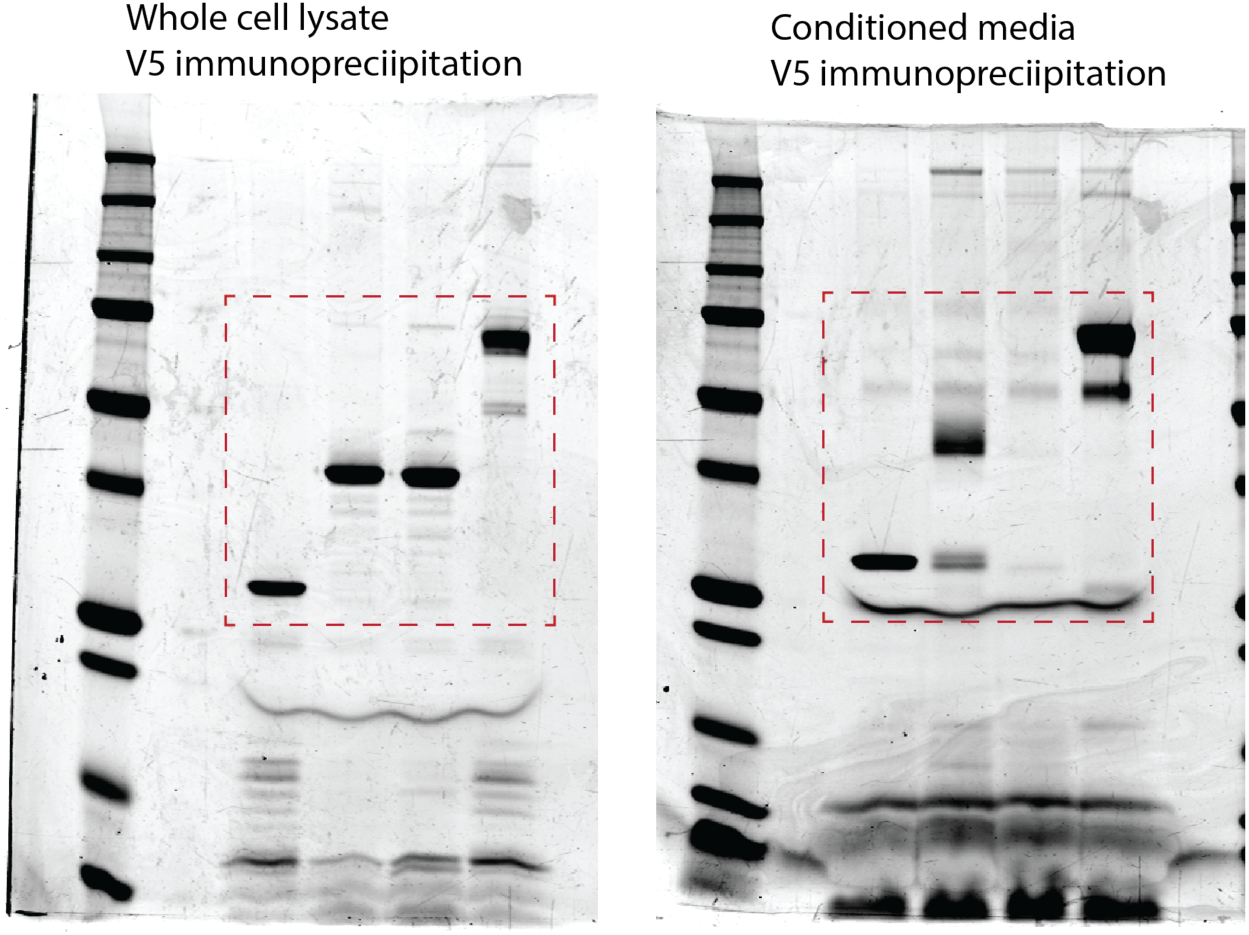
Uncropped images of GREP1 secretion Commassie stains. The dotted red box indicates the images used in the figure.

**Immunoblot Figure 4:**
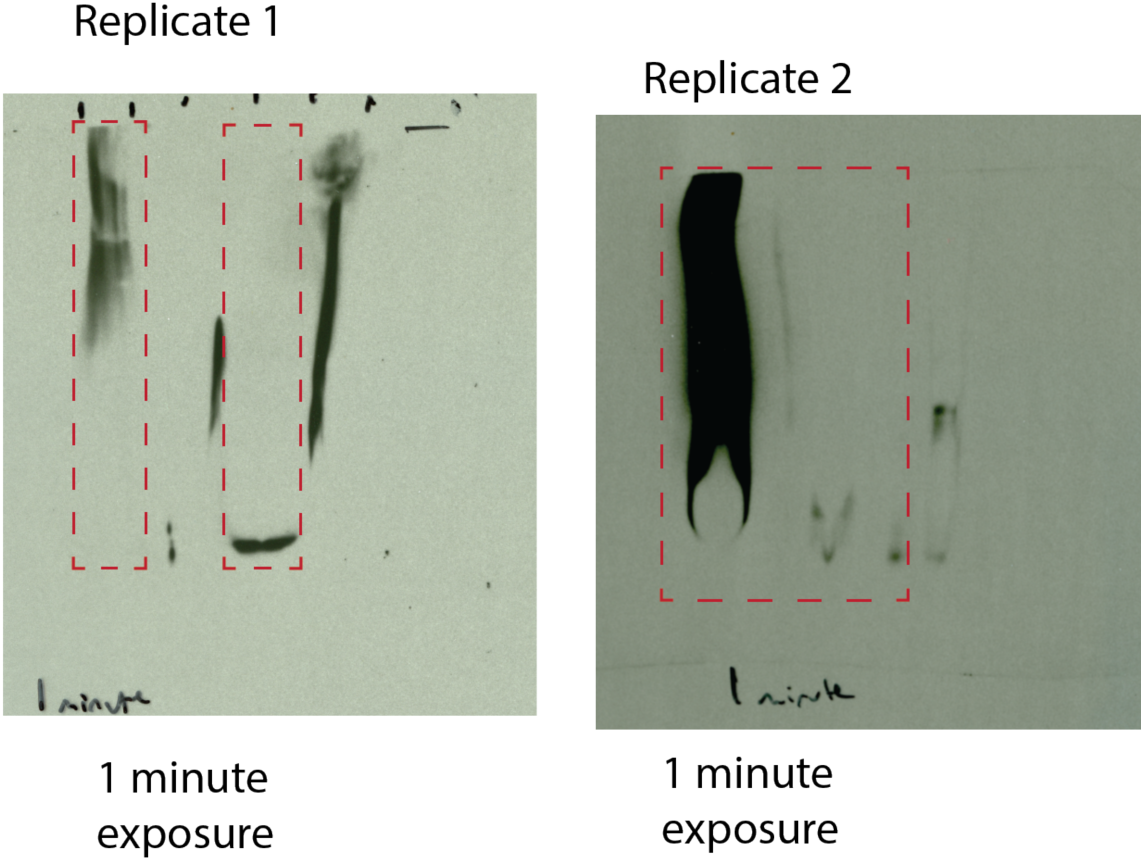
Uncropped images of GREP1 non-denaturing western blots. The dotted red box indicates the images used in the figure.

**Immunoblot Figure 5:**
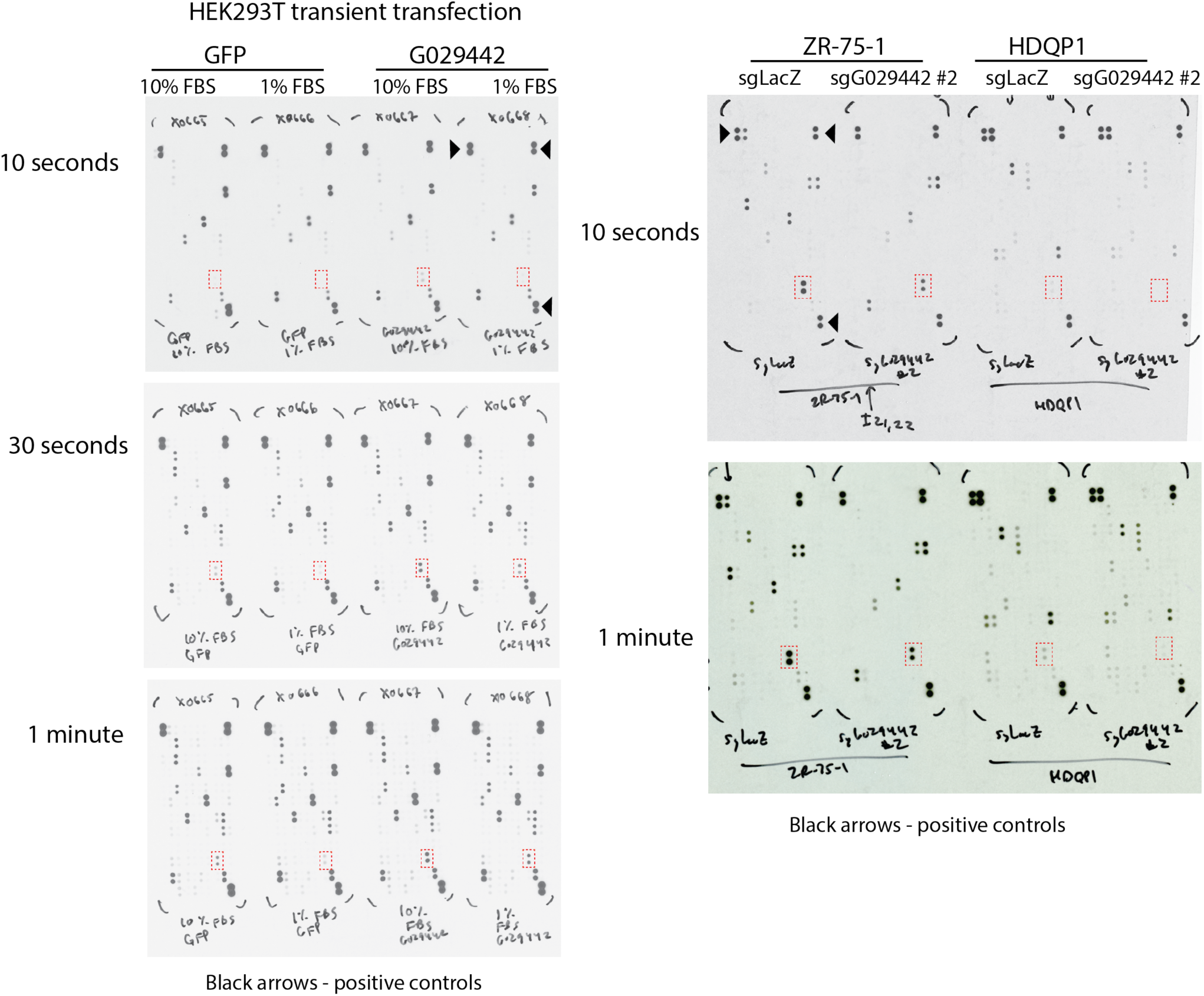
Uncropped images of cytokine array profiling. The dotted red box indicates GDF15.

## References

1. Ewing, B. & Green, P. Analysis of expressed sequence tags indicates 35,000 human genes. Nat. Genet. 25, 232–234 (2000).

2. Fields, C., Adams, M. D., White, O. & Venter, J. C. How many genes in the human genome? Nat. Genet. 7, 345–346 (1994).

3. Liang, F. et al. Gene Index analysis of the human genome estimates approximately 120,000 genes. Nat. Genet. 25, 239–240 (2000).

4. Omenn, G. S. et al. Progress on Identifying and Characterizing the Human Proteome: 2018 Metrics from the HUPO Human Proteome Project. J. Proteome Res. 17, 4031–4041 (2018).

5. Ji, Z., Song, R., Regev, A. & Struhl, K. Many lncRNAs, 5’UTRs, and pseudogenes are translated and some are likely to express functional proteins. Elife 4, e08890 (2015).

6. Ingolia, N. T. et al. Ribosome profiling reveals pervasive translation outside of annotated protein-coding genes. Cell Rep. 8, 1365–1379 (2014).

7. van Heesch, S. et al. The Translational Landscape of the Human Heart. Cell 178, 242– 260.e29 (2019).

8. Pertea, M. et al. CHESS: a new human gene catalog curated from thousands of large-scale RNA sequencing experiments reveals extensive transcriptional noise. Genome Biol. 19, 208 (2018).

9. Lander, E. S. et al. Initial sequencing and analysis of the human genome. Nature 409, 860– 921 (2001).

10. Consortium, M. G. S. & Mouse Genome Sequencing Consortium. Initial sequencing and comparative analysis of the mouse genome. Nature 420, 520–562 (2002).

11. Burge, C. & Karlin, S. Prediction of complete gene structures in human genomic DNA. J. Mol. Biol. 268, 78–94 (1997).

12. Dinger, M. E., Pang, K. C., Mercer, T. R. & Mattick, J. S. Differentiating Protein-Coding and Noncoding RNA: Challenges and Ambiguities. PLoS Comput. Biol. 4, e1000176 (2008).

13. Jungreis, I., et al. Nearly all new protein-coding predictions in the CHESS database are not protein-coding. Preprint at https://www.biorxiv.org/content/10.1101/360602v1 (2018)

14. Banfai, B. et al. Long noncoding RNAs are rarely translated in two human cell lines. Genome Res. 22, 1646–1657 (2012).

15. Mudge, J. M. et al. Discovery of high-confidence human protein-coding genes and exons by whole-genome PhyloCSF helps elucidate 118 GWAS loci. Genome Res. 29, 2073–2087 (2019).

16. Mackowiak, S. D. et al. Extensive identification and analysis of conserved small ORFs in animals. Genome Biol. 16, 179 (2015).

17. Iyer, M. K. et al. The landscape of long noncoding RNAs in the human transcriptome. Nat. Genet. 47, 199–208 (2015).

18. Bazzini, A. A. et al. Identification of small ORFs in vertebrates using ribosome footprinting and evolutionary conservation. EMBO J. 33, 981–993 (2014).

19. Calviello, L. et al. Detecting actively translated open reading frames in ribosome profiling data. Nat. Methods 13, 165–170 (2016).

20. Branca, R. M. M. et al. HiRIEF LC-MS enables deep proteome coverage and unbiased proteogenomics. Nat. Methods 11, 59–62 (2014).

21. Cabili, M. N. et al. Integrative annotation of human large intergenic noncoding RNAs reveals global properties and specific subclasses. Genes Dev. 25, 1915–1927 (2011).

22. Gao, X. et al. Quantitative profiling of initiating ribosomes in vivo. Nat. Methods 12, 147– 153 (2015).

23. Gascoigne, D. K. et al. Pinstripe: a suite of programs for integrating transcriptomic and proteomic datasets identifies novel proteins and improves differentiation of protein-coding and non-coding genes. Bioinformatics 28, 3042–3050 (2012).

24. Kim, M.-S. et al. A draft map of the human proteome. Nature 509, 575–581 (2014).

25. Koch, A. et al. A proteogenomics approach integrating proteomics and ribosome profiling increases the efficiency of protein identification and enables the discovery of alternative translation start sites. Proteomics 14, 2688–2698 (2014).

26. Ma, J. et al. Discovery of human sORF-encoded polypeptides (SEPs) in cell lines and tissue. J. Proteome Res. 13, 1757–1765 (2014).

27. Ruiz-Orera, J., Messeguer, X., Subirana, J. A. & Alba, M. M. Long non-coding RNAs as a source of new peptides. Elife 3, e03523 (2014).

28. Slavoff, S. A. et al. Peptidomic discovery of short open reading frame-encoded peptides in human cells. Nat. Chem. Biol. 9, 59–64 (2013).

29. Sun, H. et al. Integration of mass spectrometry and RNA-Seq data to confirm human ab initio predicted genes and lncRNAs. Proteomics 14, 2760–2768 (2014).

30. Vanderperre, B. et al. Direct detection of alternative open reading frames translation products in human significantly expands the proteome. PLoS One 8, e70698 (2013).

31. Wilhelm, M. et al. Mass-spectrometry-based draft of the human proteome. Nature 509, 582–587 (2014).

32. Schwaid, A. G. et al. Chemoproteomic Discovery of Cysteine-Containing Human Short Open Reading Frames. J. Am. Chem. Soc. 135, 16750–16753 (2013).

33. Zhang, C. et al. Systematic analysis of missing proteins provides clues to help define all of the protein-coding genes on human chromosome 1. J. Proteome Res. 13, 114–125 (2014).

34. Subramanian, A. et al. A Next Generation Connectivity Map: L1000 Platform and the First 1,000,000 Profiles. Cell 171, 1437–1452.e17 (2017).

35. Nassa, M. et al. Analysis of human collagen sequences. Bioinformation 8, 26–33 (2012).

36. Mullican, S. E. & Rangwala, S. M. Uniting GDF15 and GFRAL: Therapeutic Opportunities in Obesity and Beyond. Trends Endocrinol. Metab. 29, 560–570 (2018).

37. Breit, S. N., Tsai, V. W.-W. & Brown, D. A. Targeting Obesity and Cachexia: Identification of the GFRAL Receptor–MIC-1/GDF15 Pathway. Trends Mol. Med. 23, 1065–1067 (2017).

38. Baroni, M. et al. Distinct response to GDF15 knockdown in pediatric and adult glioblastoma cell lines. J. Neurooncol. 139, 51–60 (2018).

39. Huang, C.-Y. et al. Molecular alterations in prostate carcinomas that associate with in vivo exposure to chemotherapy: identification of a cytoprotective mechanism involving growth differentiation factor 15. Clin. Cancer Res. 13, 5825–5833 (2007).

40. Ratnam, N. M. et al. NF-κB regulates GDF-15 to suppress macrophage surveillance during early tumor development. J. Clin. Invest. 127, 3796–3809 (2017).

41. Corre, J. et al. Bioactivity and prognostic significance of growth differentiation factor GDF15 secreted by bone marrow mesenchymal stem cells in multiple myeloma. Cancer Res. 72, 1395–1406 (2012).

42. Peake, B. F., Eze, S. M., Yang, L., Castellino, R. C. & Nahta, R. Growth differentiation factor 15 mediates epithelial mesenchymal transition and invasion of breast cancers through IGF-1R-FoxM1 signaling. Oncotarget 8, 94393–94406 (2017).

43. Martinez, T. F. et al. Accurate annotation of human protein-coding small open reading frames. Nat. Chem. Biol. (2019) doi:10.1038/s41589-019-0425-0.

44. Chen, J. et al. Evolutionary analysis across mammals reveals distinct classes of long non-coding RNAs. Genome Biol. 17, 19 (2016).

45. Xie, W. et al. Epigenomic analysis of multilineage differentiation of human embryonic stem cells. Cell 153, 1134–1148 (2013).

46. Liu, S.J. et al. CRISPRi-based genome-scale identification of functional long noncoding RNA loci in human cells. Science 355, eaah7111 (2017).

47. Petersen, T. N., Brunak, S., von Heijne, G. & Nielsen, H. SignalP 4.0: discriminating signal peptides from transmembrane regions. Nat. Methods 8, 785–786 (2011).

48. Kelley, L. A., Mezulis, S., Yates, C. M., Wass, M. N. & Sternberg, M. J. E. The Phyre2 web portal for protein modeling, prediction and analysis. Nat. Protoc. 10, 845–858 (2015).

49. Domazet-Lošo, T., Brajković, J. & Tautz, D. A phylostratigraphy approach to uncover the genomic history of major adaptations in metazoan lineages. Trends Genet. 23, 533–539 (2007).

50. Domazet-Lošo, T. et al. No Evidence for Phylostratigraphic Bias Impacting Inferences on Patterns of Gene Emergence and Evolution. Mol. Biol. Evol. 34, 843–856 (2017).

51. Kumar, S., Stecher, G., Suleski, M. & Blair Hedges, S. TimeTree: A Resource for Timelines, Timetrees, and Divergence Times. Mol. Biol. Evol. 34, 1812–1819 (2017).

52. Yang, X. et al. A public genome-scale lentiviral expression library of human ORFs. Nat. Methods 8, 659–661 (2011).

53. Subramanian, A. et al. Gene set enrichment analysis: a knowledge-based approach for interpreting genome-wide expression profiles. Proc. Natl. Acad. Sci. USA 102, 15545– 15550 (2005).

54. Ross, Z., Wickham, H. & Robinson, D. Declutter your R workflow with tidy tools. Preprint at: https://peerj.com/preprints/3180.pdf (2017).

55. Enache, O. M. et al. The GCTx format and cmap{Py, R, M, J} packages: resources for optimized storage and integrated traversal of annotated dense matrices. Bioinformatics 35, 1427–1429 (2019).

56. Doench, J. G. et al. Optimized sgRNA design to maximize activity and minimize off-target effects of CRISPR-Cas9. Nat. Biotechnol. 34, 184–191 (2016).

57. Piccioni, F., Younger, S. T. & Root, D. E. Pooled Lentiviral-Delivery Genetic Screens. Curr. Protoc. Mol. Biol. 32.1.1–32.1.21 (2018) doi:10.1002/cpmb.52.

58. Meyers, R. M. et al. Computational correction of copy number effect improves specificity of CRISPR-Cas9 essentiality screens in cancer cells. Nat. Genet. 49, 1779–1784 (2017).

59. Hart, T., Brown, K. R., Sircoulomb, F., Rottapel, R. & Moffat, J. Measuring error rates in genomic perturbation screens: gold standards for human functional genomics. Mol. Syst. Biol. 10, 733 (2014).

60. Yu, C. et al. High-throughput identification of genotype-specific cancer vulnerabilities in mixtures of barcoded tumor cell lines. Nat. Biotechnol. 34, 419–423 (2016).

61. Niknafs, Y. S. et al. MiPanda: A Resource for Analyzing and Visualizing Next-Generation Sequencing Transcriptomics Data. Neoplasia 20, 1144–1149 (2018).

62. Shevchenko, A., Wilm, M., Vorm, O. & Mann, M. Mass spectrometric sequencing of proteins silver-stained polyacrylamide gels. Anal. Chem. 68, 850–858 (1996).

63. Peng, J. & Gygi, S. P. Proteomics: the move to mixtures. J. Mass Spectrom. 36, 1083– 1091 (2001).

64. Eng, J. K., McCormack, A. L. & Yates, J. R. An approach to correlate tandem mass spectral data of peptides with amino acid sequences in a protein database. J. Am. Soc. Mass Spectrom. 5, 976–989 (1994).

65. Beausoleil, S. A., Villén, J., Gerber, S. A., Rush, J. & Gygi, S. P. A probability-based approach for high-throughput protein phosphorylation analysis and site localization. Nat. Biotechnol. 24, 1285–1292 (2006).

66. Jones, D. T. & Cozzetto, D. DISOPRED3: precise disordered region predictions with annotated protein-binding activity. Bioinformatics 31, 857–863 (2015).

## Supplementary references

1. Zoubak, S., Clay, O. & Bernardi, G. The gene distribution of the human genome. Gene 174, 95–102 (1996).

2. Pruitt, K. D. & Maglott, D. R. RefSeq and LocusLink: NCBI gene-centered resources. Nucleic Acids Res. 29, 137–140 (2001).

3. Benson, D. A., Boguski, M., Lipman, D. J. & Ostell, J. GenBank. Nucleic Acids Research vol. 22 3441–3444 (1994).

4. Fields, C., Adams, M. D., White, O. & Venter, J. C. How many genes in the human genome? Nat. Genet. 7, 345–346 (1994).

5. Schuler, G. D. et al. A gene map of the human genome. Science 274, 540–546 (1996).

6. Liang, F. et al. Gene Index analysis of the human genome estimates approximately 120,000 genes. Nat. Genet. 25, 239–240 (2000).

7. Consortium, I. H. G. S. & International Human Genome Sequencing Consortium. Finishing the euchromatic sequence of the human genome. Nature vol. 431 931–945 (2004).

8. Omenn, G. S. et al. Progress on Identifying and Characterizing the Human Proteome: 2019 Metrics from the HUPO Human Proteome Project. Journal of Proteome Research (2019) doi:10.1021/acs.jproteome.9b00434.

9. Borsani, G. et al. Characterization of a murine gene expressed from the inactive X chromosome. Nature vol. 351 325–329 (1991).

10. Brockdorff, N. et al. The product of the mouse Xist gene is a 15 kb inactive X-specific transcript containing no conserved ORF and located in the nucleus. Cell vol. 71 515–526 (1992).

11. Clamp, M. et al. Distinguishing protein-coding and noncoding genes in the human genome. Proceedings of the National Academy of Sciences vol. 104 19428–19433 (2007).

12. Burge, C. & Karlin, S. Prediction of complete gene structures in human genomic DNA. Journal of Molecular Biology vol. 268 78–94 (1997).

13. Consortium, M. G. S. & Mouse Genome Sequencing Consortium. Initial sequencing and comparative analysis of the mouse genome. Nature vol. 420 520–562 (2002).

14. Dinger, M. E., Pang, K. C., Mercer, T. R. & Mattick, J. S. Differentiating Protein-Coding and Noncoding RNA: Challenges and Ambiguities. PLoS Computational Biology vol. 4 e1000176 (2008).

15. Kumar, N. & Kishore, R. Determination of an unusual secondary structural element in the immunostimulating tetrapeptide rigin in aqueous environments: insights via MD simulations, 1H NMR and CD spectroscopic studies. Journal of Peptide Science (2010) doi:10.1002/psc.1260.

16. Marcotte, I., Separovic, F., Auger, M. & Gagné, S. M. A Multidimensional 1H NMR Investigation of the Conformation of Methionine-Enkephalin in Fast-Tumbling Bicelles. Biophysical Journal vol. 86 1587–1600 (2004).

17. Imura, T. et al. Minimum Amino Acid Residues of an α-Helical Peptide Leading to Lipid Nanodisc Formation. Journal of Oleo Science vol. 63 1203–1208 (2014).

18. Anderson, D. M. et al. A Micropeptide Encoded by a Putative Long Noncoding RNA Regulates Muscle Performance. Cell vol. 160 595–606 (2015).

19. Nelson, B. R. et al. A peptide encoded by a transcript annotated as long noncoding RNA enhances SERCA activity in muscle. Science 351, 271–275 (2016).

20. Iyer, M. K. et al. The landscape of long noncoding RNAs in the human transcriptome. Nat. Genet. 47, 199–208 (2015).

21. Cabili, M. N. et al. Integrative annotation of human large intergenic noncoding RNAs reveals global properties and specific subclasses. Genes & Development vol. 25 1915–1927 (2011).

22. Barretina, J. et al. The Cancer Cell Line Encyclopedia enables predictive modelling of anticancer drug sensitivity. Nature 483, 603–607 (2012).

23. Neme, R. & Tautz, D. Phylogenetic patterns of emergence of new genes support a model of frequent de novo evolution. BMC Genomics 14, 117 (2013).

24. Wilson, B. A., Foy, S. G., Neme, R. & Masel, J. Young genes are highly disordered as predicted by the preadaptation hypothesis of de novo gene birth. Nature Ecology & Evolution vol. 1 (2017).

25. Kelley, L. A., Mezulis, S., Yates, C. M., Wass, M. N. & Sternberg, M. J. E. The Phyre2 web portal for protein modeling, prediction and analysis. Nat. Protoc. 10, 845–858 (2015).

26. D’Lima, N. G. et al. A human microprotein that interacts with the mRNA decapping complex. Nat. Chem. Biol. 13, 174–180 (2017).

27. Paik, Y.-K. et al. Launching the C-HPP neXt-CP50 Pilot Project for Functional Characterization of Identified Proteins with No Known Function. J. Proteome Res. 17, 4042–4050 (2018).

28. Omenn, G. S. et al. Progress on Identifying and Characterizing the Human Proteome: 2018 Metrics from the HUPO Human Proteome Project. J. Proteome Res. 17, 4031–4041 (2018).

29. Makarewich, C. A. et al. MOXI Is a Mitochondrial Micropeptide That Enhances Fatty Acid β-Oxidation. Cell Rep. 23, 3701–3709 (2018).

30. Chugunova, A. et al. LINC00116 codes for a mitochondrial peptide linking respiration and lipid metabolism. Proc. Natl. Acad. Sci. U. S. A. 116, 4940–4945 (2019).

31. Sun, M. et al. A Human Novel Gene DERPC Located on 16q22.1 Inhibits Prostate Tumor Cell Growth and Its Expression Is Decreased in Prostate and Renal Tumors. Molecular Medicine vol. 8 655–663 (2002).

32. Liu, L. et al. Interaction between p12CDK2AP1 and a novel unnamed protein product inhibits cell proliferation by regulating the cell cycle. Mol. Med. Rep. 9, 156–162 (2014).

33. Rathore, A. et al. MIEF1 Microprotein Regulates Mitochondrial Translation. Biochemistry 57, 5564–5575 (2018).

34. Akimoto, C. et al. Translational repression of the McKusick–Kaufman syndrome transcript by unique upstream open reading frames encoding mitochondrial proteins with alternative polyadenylation sites. Biochimica et Biophysica Acta (BBA) - General Subjects 1830, 2728– 2738 (2013).

35. Matsumoto, A. et al. mTORC1 and muscle regeneration are regulated by the LINC00961-encoded SPAR polypeptide. Nature vol. 541 228–232 (2017).

